# Unravelling the functional diversity of type III polyketide synthases in fungi

**DOI:** 10.1101/2024.11.25.625244

**Authors:** Nika Sokolova, Stepan S. Denisov, Thomas Hackl, Kristina Haslinger

**Affiliations:** Department of Chemical and Pharmaceutical Biology, University of Groningen, The Netherlands; Institute of Biological Chemistry, University of Vienna, Vienna, Austria; RDM Cardiovascular Medicine, University of Oxford, Oxford, The United Kingdom; Groningen Institute for Evolutionary Life Sciences, University of Groningen, Groningen, The Netherlands

## Abstract

Type III polyketide synthases (T3PKSs) are enzymes that produce diverse compounds of ecological and clinical importance. While well-studied in plants and bacteria, only a handful of T3PKSs from fungi have been characterised to date. Here, we developed a comprehensive workflow for kingdom-wide characterisation of T3PKSs. Using publicly available genomes, we mined more than 1000 putative enzymes and analysed their active site architecture and genomic neighbourhood. From there, we selected 37 representative PKS candidates for cell-free expression and prototyping with a diverse set of Coenzyme A activated substrates, revealing unique patterns in substrate and cyclisation specificity, as well as the preferred number of malonyl-Coenzyme A extensions. Using the 341 enzyme-substrate pairs generated in this study, we trained a machine learning model to predict T3PKS substrate specificity and experimentally validated it with an extended panel of non-natural substrates. We anticipate that the model will be useful for *in silico* screening of T3PKSs, while the insight into the product scope of these enzymes offers interesting starting points for further exploration.

## Introduction

The production of bulk and fine chemicals relies heavily on fossil fuels and rare earth catalysts. Biocatalysis, capitalising on renewable starting materials and enzyme catalysts, holds the potential to shift the paradigm towards greener synthesis of drugs^1^, industrial chemicals^2^ and polymers^3^. Despite the growing interest of pharmaceutical and fragrance companies in enzymatic processes, the wider use of biocatalysis is fundamentally hindered by our ability to select enzymes for reactions and substrates of interest^4,5^. To be able to compete with the organic synthesis toolbox refined over several centuries, biocatalysis needs more promiscuous enzymes that are active on non-native substrates^6^.

Enzymes catalysing the formation of molecular scaffolds that are commonly found in marketed drugs are particularly interesting^7^. One example are type III polyketide synthases (T3PKSs) giving rise to distinct polyketide scaffolds (Fig. 1a). These enzymes use coenzyme A (CoA)-bound substrates for iterative decarboxylation, elongation, cyclisation, and aromatisation – all within a single active site – with variable number of elongations (ketide number) and three distinct cyclisation mechanisms^8^. Thanks to their relaxed substrate specificity, broad product range, and structural simplicity, T3PKSs comprise a rich pool for the selection and evolution of powerful biocatalysts.

**Figure 1:**
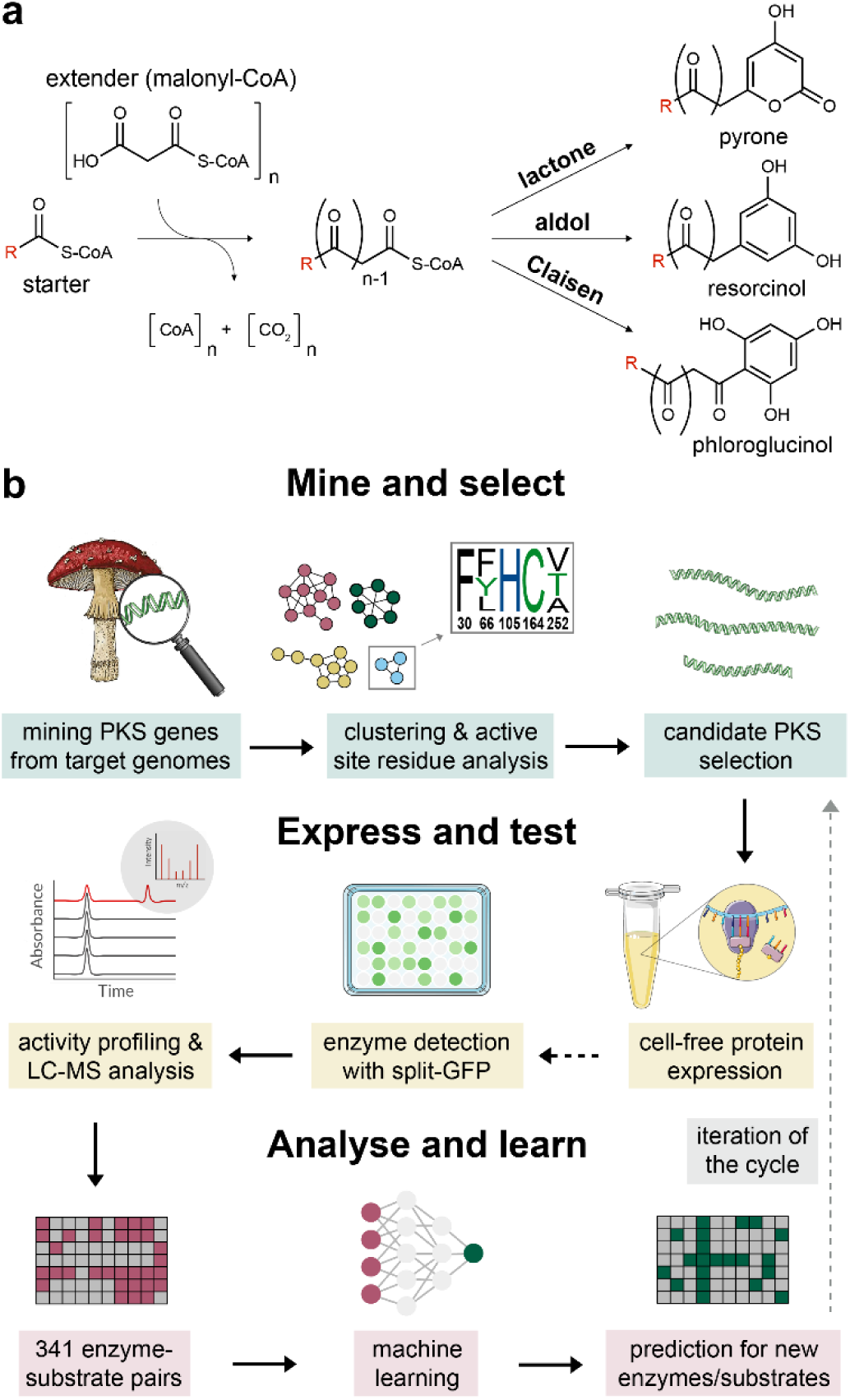
**Scope of this study.** a) Typical reactions catalysed by T3PKSs. The extender unit (usually malonyl-CoA) is iteratively decarboxylated and condensed with the starter unit (variable). The linear polyketide is then cyclised through one of the three possible mechanisms yielding pyrone (lactone), resorcinol (aldol), or phloroglucinol (Claisen) products. b) The workflow for kingdom-wide characterisation of T3PKS. Optional steps are represented with dotted arrows.

T3PKSs are best studied in plants, where they synthesise more than 10 different groups of natural products^9^. For instance, plant T3PKSs have been employed in microbial production routes towards medicinally relevant cannabinoid, quinolone, and tropane alkaloid scaffolds^9^. In bacteria, T3PKSs synthesise naphtalene pigments, spore germination inhibitors, and phenolic lipids required for cyst formation^10^. The first fungal representative, 2-oxoalkylresorcylic acid synthase from *Neurospora crassa*, was characterised in 2007^11^. Since then, several more T3PKSs were cloned from filamentous fungi^12–21^ and yeast^22^, and genome mining projects hint at the presence of many more^23^. Most fungal T3PKSs can synthesise tri-, tetra- or pentaketide pyrones and/or resorcinols from a range of fatty acyl-CoA substrates *in vitro* but lack activity on ring-type substrates typical for plant T3PKSs^8^. However, reports on fungal T3PKSs remain somewhat anecdotal to date, and our understanding of their functional diversity and native functions is lagging compared to their plant and bacterial counterparts.

Here, we devised a comprehensive workflow for the characterisation of fungal T3PKSs based on computational and experimental methods (Fig. 1b). We closely examined the entire fungal T3PKS sequence space and selected 37 representative candidates for cell-free expression and experimental characterisation. We then mapped the activity landscape of candidate T3PKSs using a panel of CoA- bound substrates, revealing unexpected products and highlighting enzymes worth further investigation. With this unique set of enzyme-substrate pairs at hand, we trained and experimentally validated the first machine learning model for T3PKS substrate specificity prediction. We also applied machine learning to propose residues important for substrate specificity. Lastly, we examined the genomic context of these enzymes to derive hypotheses on their elusive natural functions in fungi.

## Results

### Kingdom-wide genome mining suggests functionally diverse T3PKSs

First, we set out to explore the prevalence and diversity of T3PKSs in the fungal kingdom. To mine for enzyme candidates, we collected sequences of experimentally characterised T3PKSs from plants and fungi (Supplementary Data 1) and used them to query 2096 fungal genomes (on 2021-12-22) available in the JGI MycoCosm database^24^. A total of 1148 putative T3PKSs were detected in 806 fungal strains (38% of all included genomes). Of these, 582 carried a single sequence and 224 had between 2 and 7 copies of T3PKS genes (Supplementary Data 2). In parallel, we also retrieved all putative T3PKS sequences from the InterPro database^25^ (domain IPR011141), since we had noticed that the MycoCosm and InterPro databases are not mutually redundant. The merged and dereplicated dataset contained 1640 sequences (Supplementary Data 3).

To identify potential functional groups of fungal T3PKSs, we constructed a sequence similarity network of the mined T3PKSs using the EFI-EST webtool^27^ (Supplementary Data 4). In this network, a sequence identity cutoff of around 57% resulted in cluster separation that roughly reflected the phylogenetic clades defined in a recent evolutionary analysis of fungal T3PKSs^23^ (Supplementary Fig. 1). However, we further increased the cutoff to 80% (Fig. 2a) in view of literature evidence that plant T3PKSs can employ distinct cyclisation mechanisms despite sharing 74% sequence identity^28^. Even at such a high cutoff, the clusters were not exclusively monogeneric, and cluster 11 contained sequences from up to three different fungal classes. Previously characterised T3PKSs localised to smaller clusters on the network or appeared as singletons, suggesting that they may not be representative of the typical fungal T3PKSs. As is commonly observed with biosynthetic genes in fungi^29^, sequences mined from *Aspergillus* separated into more than 10 clusters in the network, hinting at distinct functional groups of T3PKSs in this genus.

**Figure 2:**
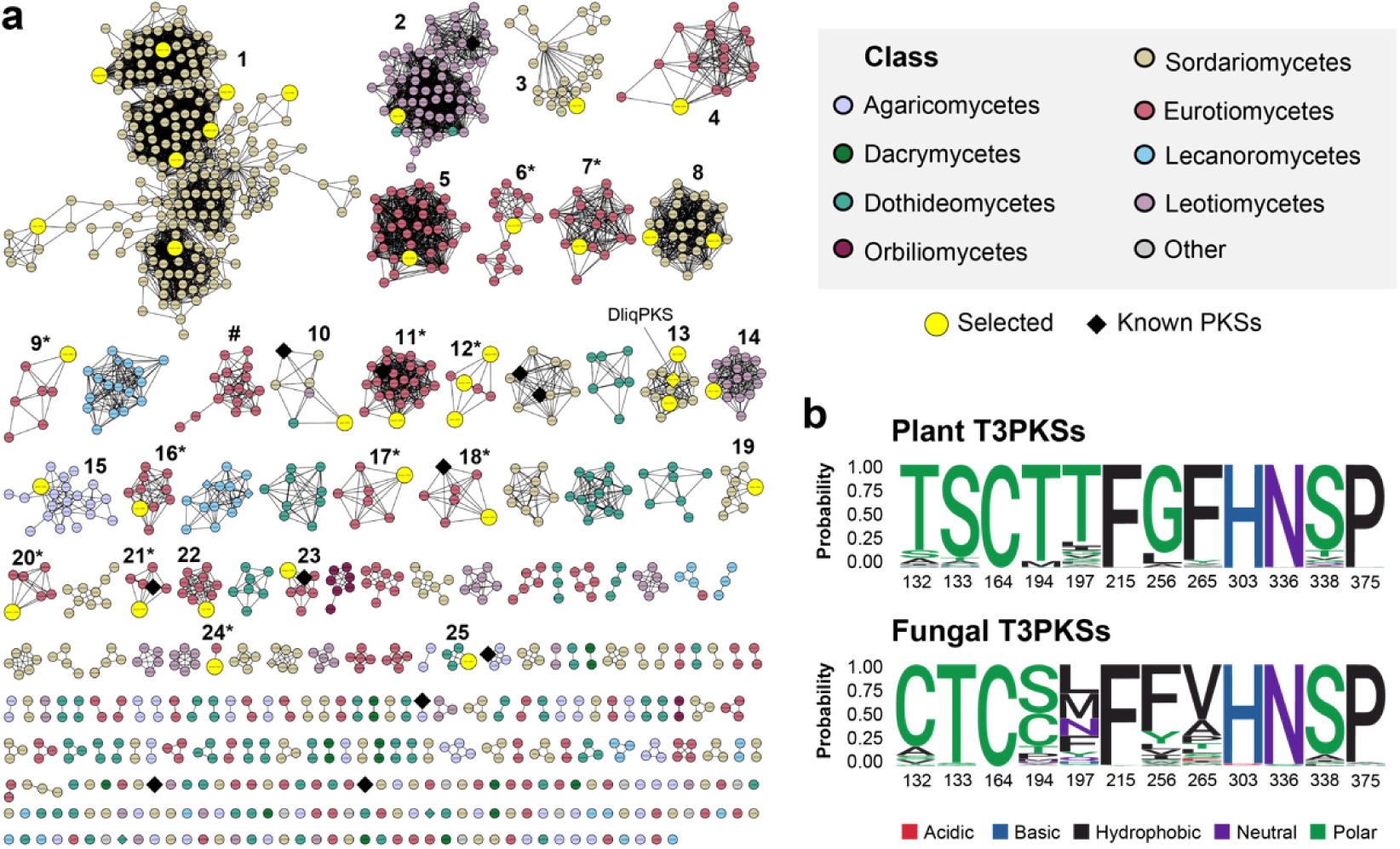
**Clustering and active site residue analysis of fungal T3PKSs.** a) Sequence similarity network of the mined enzyme candidates at 80% sequence identity cutoff. Each circle is a representative node grouping protein sequences with >95% sequence identity. Diamond-shaped nodes represent enzymes with literature precedent. Clusters from which sequences were selected for experimental characterisation are numbered. Clusters composed exclusively of sequences from *Aspergillus* are marked with an asterisk. The only major cluster where the catalytic triad is not conserved is marked with # (amino acid substitutions: H303D and N336S). b) Comparison of the active site residues between plant T3PKSs (184 reviewed sequences from InterPro) and the mined fungal T3PKSs. The numbering of the residues corresponds to that in MsCHS (model chalcone synthase from *Medicago sativa*). The active site residue logo was generated using the R package ggseqlogo^26^.

We zoomed into each cluster and compared the predicted active site residues of fungal sequences to their well-studied plant counterparts (Fig. 2b). The catalytic cysteine (residue 164 in MsCHS, the model chalcone synthase from *Medicago sativa*^30^) is conserved across all fungal T3PKSs, while the other residues of the catalytic triad, H303 and N336, were conserved in all but one major cluster. However, one of the residues critical for substrate recognition, the "gatekeeper" F265^30^, is variable in fungal T3PKSs (Fig. 3b). The F265V substitution, observed in half of the sequences, is also characteristic of divergent plant T3PKSs that produce quinolone and acridone alkaloids from anthranilic acid derivatives^31^. The conserved G256 of the substrate-binding pocket, replaced with leucine in some divergent plant T3PKSs^32^, is substituted with bulky aromatic residues in most fungal enzymes. In addition, position 197, which is known to influence substrate^33^ and product specificity^28^ in plant chalcone synthases, is variable in fungal enzymes and comprises mostly hydrophobic residues. Overall, the observed variations in the active site residues suggested potential functional differences among fungal T3PKS enzymes.

**Figure 3:**
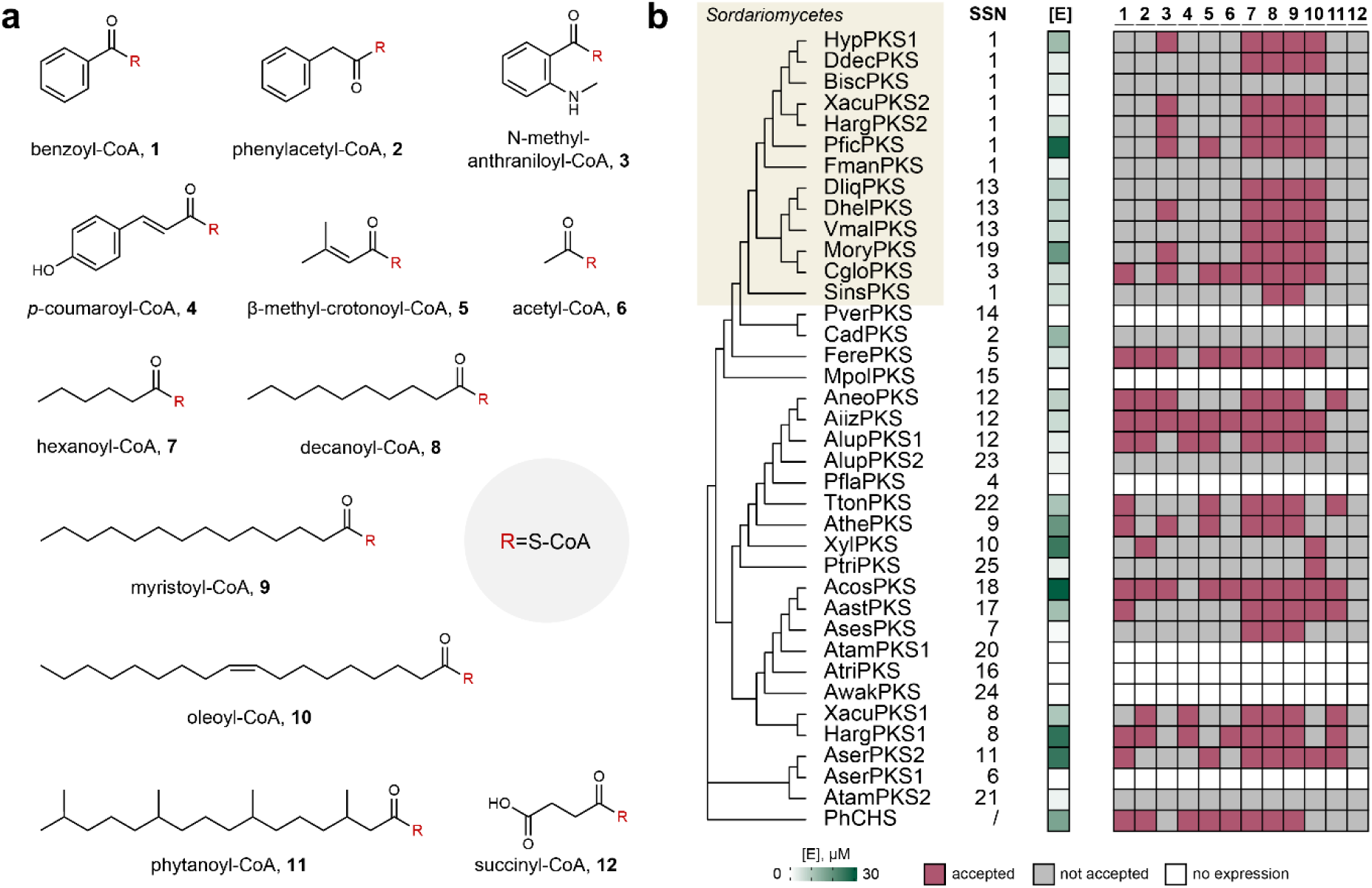
**Activity profiling of the cell-free expressed T3PKSs.** a) Panel of CoA thioesters **1**-**12** used as substrates for activity profiling. b) Maximum-likelihood phylogenetic tree based on the multiple sequence alignment of 37 fungal T3PKSs (outgroup: PhCHS) combined with a heatmap of enzyme concentrations in the cell-free lysate and a binary representation of enzyme activity in cell-free lysates supplemented with substrates **1**-**12** and malonyl-CoA. The "SSN" column specifies the corresponding cluster in the sequence similarity network. Enzyme concentrations [E] (0 to 30 µM) were determined with a split-GFP complementation assay.

We then proceeded to sample sequences for experimental characterisation. During selection, we aimed to capture the taxonomic and ecological diversity of the host organisms, as well as the putative functional diversity hinted at by the active site composition. We also oversampled cluster 1 due to its large size and apparent separation into several subclusters reflecting the taxonomic orders of the donor organisms (Supplementary Fig. 2). When selecting sequences within each cluster, we prioritsed those with active site residues corresponding to the cluster consensus and those stemming from highly contiguous, high-quality genome assemblies. Our positive controls included a chalcone synthase from *Petunia hybrida* (PhCHS, UniProt: P08894) and a reported fungal chalcone synthase from *Diaporthe liquidambaris* (DliqPKS, UniProt: A0A7T8G346)^34^. In total, we selected 37 fungal genes for experimental characterisation (Supplementary Table 1, Supplementary Data 5).

### Activity profiling of the cell-free expressed T3PKSs

We then turned to functional characterisation of the candidate enzymes. To facilitate the express-test workflow, we used the recently established myTXTL® cell-free expression platform from linear DNA templates^35^. We designed each DNA fragment to contain the codons for split-GFP (green fluorescent protein) and a hexahistidine tag in-frame with the 3’ end of the target gene. The split-GFP tag was used to quantify the expression levels of the target proteins via split-GFP complementation assay^36^. We first verified the express-test workflow using our positive control enzyme PhCHS. The myTXTL® lysate containing 0.78 mg/mL cell-free expressed PhCHS successfully converted coumaroyl-CoA and malonyl-CoA to naringenin chalcone, a part of which then spontaneously cyclised to naringenin (Supplementary Fig. 3).

Next, we collected a panel of substrates for enzyme activity profiling (Fig. 3a). During selection, we attempted to capture the diversity of known T3PKS substrates across all phylogenetic groups. Thus, in addition to **4**, which is the canonical substrate of plant chalcone synthases, we included substrates of other plant T3PKSs such as benzophenone and quinolone synthases (**1** and **3**, respectively). Since previously characterised fungal and bacterial T3PKSs tend to prefer aliphatic over aromatic substrates^8^, we chose a selection of fatty acyl-CoA substrates of different chain length, level of branching and degree of unsaturation (**6-10**). Finally, we picked several more acyl-CoA thioesters that, to the best of our knowledge, were never tested with T3PKSs before (**11-12**).

We screened the resulting panel of 12 substrates with malonyl-CoA as a co-substrate against 38 cell-free expressed enzymes and analysed the products with liquid chromatography–coupled mass spectrometry (LC-MS). An enzyme was deemed active if we detected one or several new peaks with *m/z* values consistent with the elongation of the carbon chain with up to six CH_2_CO units followed by cyclisation. We also cross-compared reactions with the same enzyme but different substrates to eliminate any possible background peaks that show up in multiple samples. Based on the results of the split-GFP assay, 31 enzymes were expressed at detectable levels (Fig. 3b). Of these, 26 enzymes were active on at least one substrate. The most promiscuous enzyme, AiizPKS, accepted 10 out of 12 substrates, while the plant enzyme PhCHS accepted 8. The reported fungal chalcone synthase DliqPKS did not produce naringenin chalcone from coumaroyl-CoA under our assay conditions but was active on fatty acyl substrates **6-10**. **12** was the only substrate not accepted by any enzyme, possibly due to electronic effects, or because it was diverted to other reactions in the cell-free lysate. The most broadly accepted substrates in the panel were **8** and **9**, for which 24 out of 26 active enzymes yielded product, followed by **6** (23 enzymes) and **10** (18 enzymes). We observed some specificity for the fatty acyl-CoA substrates **7-10** in the clade consisting of sequences from Sordariomycetes (Fig. 3b). The ability to accept aromatic substrates did not correlate with the phylogeny of the enzymes seen in the multiple sequence alignment or the SSN and was scattered across the sequence space. Among the pairs of enzymes that originate from the same organism, in only two both enzymes were active (XacuPKS1 and 2, and HargPKS1 and 2), and the activity profiles within the pairs differed in the ability to accept aromatic and highly branched substrates.

### Machine learning can predict substrate specificity of fungal T3PKSs

Our activity profiling showed that the substrate specificity varied even across enzymes from closely related organisms and could not be predicted with human intelligence from phylogeny alone. This led us to hypothesise that machine learning might capture the complex patterns of substrate preference more efficiently. Thus, we used the binary activity data of 31 expressed enzymes with 11 substrates (excluding **12**, which was not accepted by any of the enzymes) to train a machine learning model for substrate specificity prediction in fungal T3PKSs (Fig. 4a). In total, 341 enzyme/substrate pairs were included in the data set, of which 146 (43%) were active. Enzymes were represented by calculating and averaging their per-residue embeddings whereas substrate features were represented by extended-connectivity fingerprints (ECFPs). The final representation of each enzyme/substrate pair was obtained by concatenating enzyme and substrate features.

**Figure 4:**
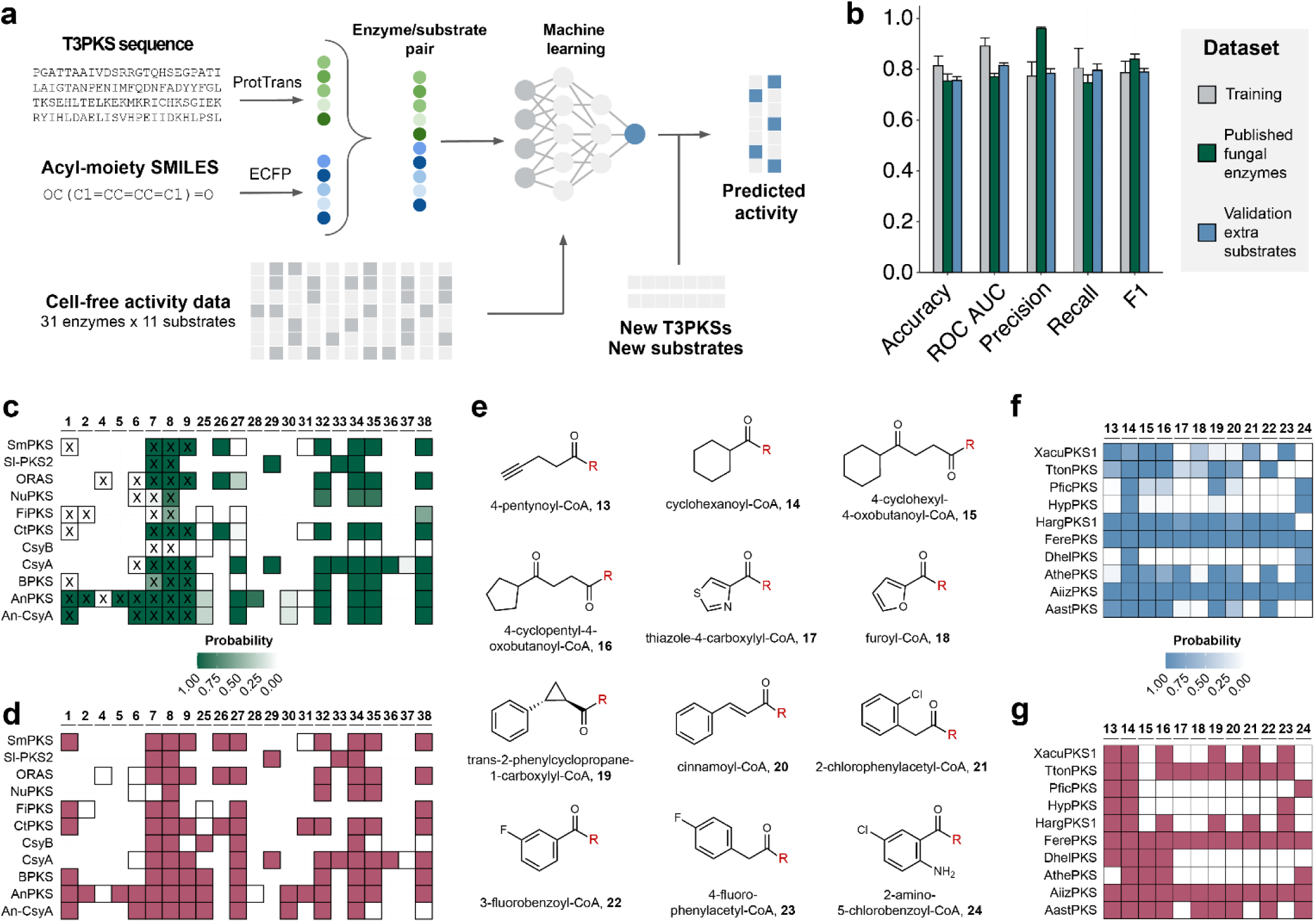
A machine learning model for T3PKS substrate specificity prediction. a) The workflow used to train the predictive machine learning model in this study. b) Performance metrics of the Multilayer Perceptron model for different datasets. The model was trained on the cell-free activity profiling dataset with ProtTrans-X5/MACCS Keys feature vectors; error bars indicate standard deviation. c) Heatmap of the predicted substrate specificity for 11 previously characterised T3PKSs from fungi. The structures of substrates 25-38 are depicted in Supplementary Fig. 6. Substrates that were present in the training dataset of our model are marked with X. d) The published data on substrate specificity for the same group of enzymes. We used the activity data from in vitro enzymatic assays reported for SmPKS from Sordaria macrospora17, Sl-PKS2 from Sporotrichum laxum16, ORAS from Neurospora crassa11, NuPKS from Naganishia uzbekistanensis22, FiPKS from Fusarium incarnatum20, CtPKS from Chaetomium thermophilum17, CsyB37 and CsyA38 from Aspergillus oryzae, BPKS from Botrytis cinerea15, AnPKS from Aspergillus niger CBS 513.8812, and An-CsyA from A. niger NRRL 32839. e) Structures of substrates 13-24 used for experimental validation of the model; R = S-CoA. f) Heatmap of the predicted substrate specificity for 10 fungal T3PKSs with non-natural substrates 13-24, and g) the substrate specificity observed through in vitro assays. All prediction values are averaged from 100 model runs with a random seed.

We assessed three machine learning algorithms – Decision tree, Random Forest and feedforward neural network Multilayer Perceptron (MLP) – for binary classification of enzyme activity with three different enzyme/substrate representations (Supplementary Fig. 4). The model accuracy and F1 scores across all tested combinations averaged at 79% and 0.764, respectively. The best-performing combinations were the MLP model with ProtTrans-X5/Morgan fingerprints (accuracy 82 ± 4%, F1 score 0.8 ± 0.05) and ProtTrans-X5/MACCS Keys representations (accuracy 81 ± 4%, F1 score 0.79 ± 0.05) (Supplementary Fig. 4). As no statistically significant difference was found between these models, the shorter ProtTrans-X5/MACCS Keys combination was used for the following experiments (Fig. 4b).

We then evaluated the performance of the model on enzymes which were not included in the training dataset. For that purpose, we collected published data on substrate specificity of 11 T3PKSs from fungi, yielding a total of 110 enzyme/substrate pairs (Fig. 4c). The dataset exhibited strong positivity bias, as 95 enzyme/substrate pairs (86% of the total number) were active. In addition, more than half of the data points (65) were composed of both the enzyme and the substrate that were not included in our training dataset. When tested on this dataset, the calculated accuracy and F1 score for our model reached 75 ± 3% and 0.84 ± 0.02, respectively (Fig. 4d). Additionally, to assess the phylogenetic bias of our model, we gathered 122 enzyme/substrate pairs for 6 plant and 9 bacterial T3PKSs (Supplementary Table 2), of which 88 were active (72%). When tested on this dataset, the model had lower performance with 56 ± 4% accuracy and 0.7 ± 0.04 F1 score (Supplementary Fig. 5).

### Experimental validation of the machine learning model on additional substrates

To further test the limits of our model’s applicability, we performed experimental validation with a panel of substrates that were not present in the training dataset. We predicted activity for all our PKSs with 12 chemically diverse carboxylic acids that we found were amenable to enzymatic CoA ligation using Os4CL from *Oryza sativa*^40^ or PqsA from *Pseudomonas aeruginosa*^41^ (Fig. 4e, Supplementary Fig. 7). From there, we chose 10 enzymes with varying degrees of predicted substrate tolerance for experimental testing. The validation dataset thus consisted of 120 enzyme-substrate pairs, of which 69 (57.5%) were predicted to be active (Fig. 4f). We cloned the genes with a C-terminal hexahistidine-tag, expressed them in *E. coli* and purified them to homogeneity alongside the two CoA ligases. We then tested the substrate specificity of the T3PKSs in a two-step cascade reaction and analysed the products with LC-MS (Fig. 4g, Supplementary Fig. 8). The accuracy of the model was marginally lower than that within the training set (76 ± 2% accuracy and 0.79 ± 0.01 F1 score), indicating its suitability for predictions of substrate specificity with new substrates (Fig. 4b). Of note, the model made accurate predictions even for substrates **13-16** that are relatively distant from those present in the training set (Supplementary Fig. 9).

### Machine learning proposes amino acid residues important for substrate specificity

While our model makes accurate predictions of activity for new enzymes and substrates, it does not provide insight into the decision-making process. Therefore, we employed a descriptive machine learning approach to pinpoint the residues that might be important for substrate specificity (Fig. 5a). Instead of featurising the whole protein sequence, we retrieved active site residues based on the naringenin-bound structure of MsCHS (PDB: 1CGK), as well as those lining the fatty acid-binding tunnel according to the structure of ORAS (PDB: 3UET). For all 31 active fungal PKSs, we featurised these 41 amino acids and used them together with the activity data for the ML model training. For this purpose, the Decision Tree approach was used as a simplistic model that is easier to interpret compared to more sophisticated algorithms. The resulting model achieved 79.4 ± 4.0% accuracy and 0.76 ± 0.045 F1 score (Fig. 5b). The analysis of feature importances in the model showed that tunnel residues L26, A30 and N65, as well as the active site residue H257, were the most important for classification accuracy (Fig. 5c-e).

**Figure 5:**
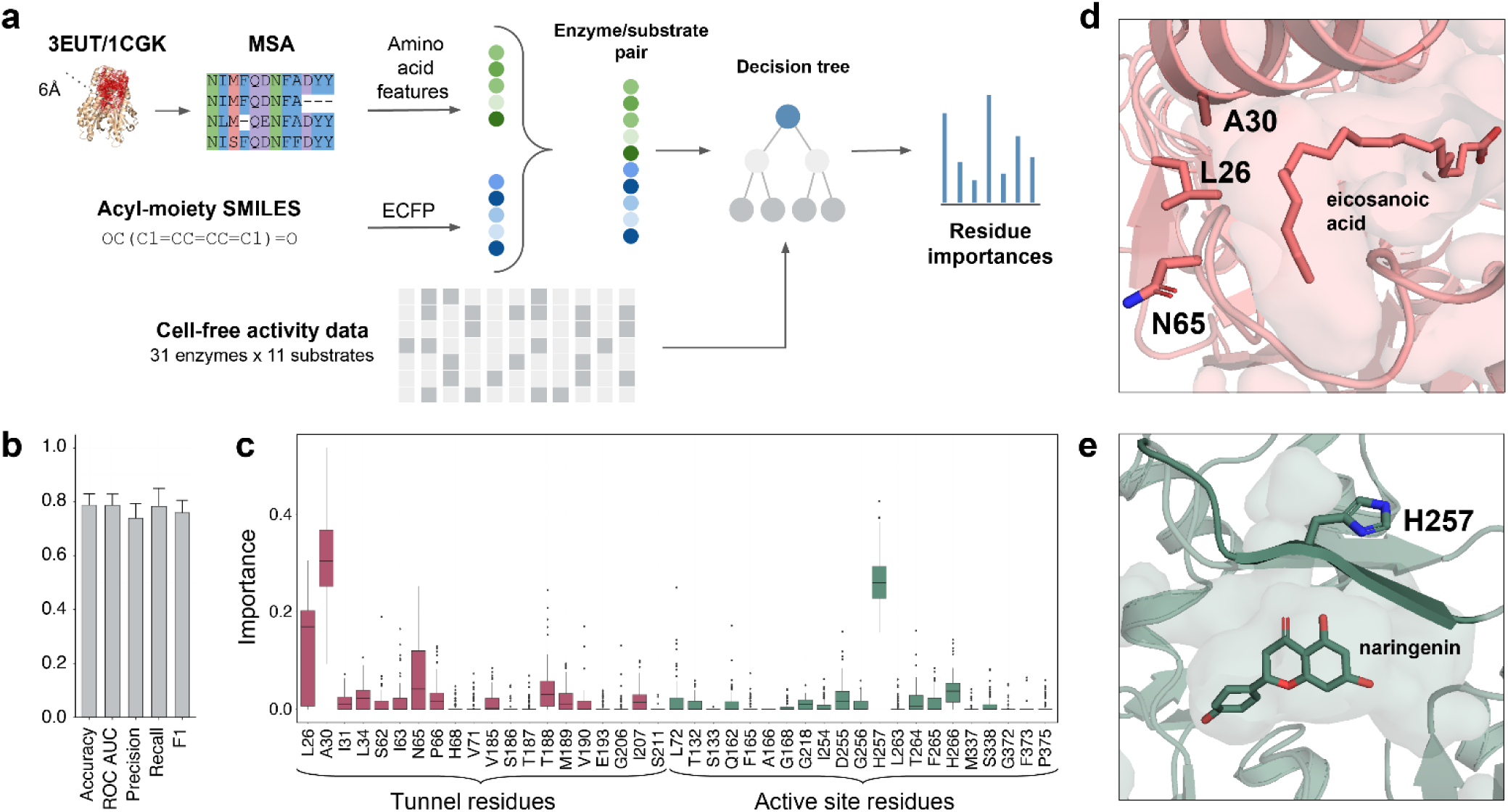
The descriptive machine learning model. a) The workflow used to train the descriptive machine learning model in this study. MSA - multiple sequence alignment; ECFP - extended-connectivity fingerprints. 3EUT – crystal structure of ORAS complexed with eicosanoic acid^42^; 1CGK – crystal structure of MsCHS complexed with naringenin^30^. b) Performance metrics of the Decision Tree model trained on the cell-free activity profiling dataset with per-residue molecular descriptor and MACCS Keys feature vectors; error bars indicate standard deviation. c) The importances of the active site and substrate tunnel residues represented as boxplots. Importance values were averaged from 100 model runs with random seeds. Numbering of the tunnel residues is taken from ORAS; numbering of the active site residues is taken from MsCHS. Cartoon representation of d) the substrate tunnel in ORAS and e) active site in MsCHS with the three most important residues shown as sticks (blue: nitrogen atoms, red: oxygen atoms).

L26, A30 and N65 are the residues capping the surface end of the substrate tunnel in ORAS. The role of these residues is currently understudied, but it is conceivable that they are important for the orientation of longer-chain fatty acyl substrates. Interestingly, position 26 is mostly occupied by the hydrophobic amino acids leucine, isoleucine, valine and methionine among the tested T3PKS with the exception of two enzymes, XylPKS (cysteine) and AtamPKS2 (arginine). XylPKS was only active on two substrates (**7** and **10**), whereas AtamPKS2 was inactive. This may indicate that sequence variation in this position leads to a stricter substrate specificity, possibly outside the scope of the tested substrates. The sequence diversity at positions 30 and 65 is quite high across the selected T3PKS and we were not able to rationalise their individual contributions to the substrate specificities.

While the active site residue H257 is too distant to directly interact with the substrate, it likely participates in salt bridge formation between the T3PKS monomers^43^. Its mutation to lysine was shown to contribute to a shift in the cyclisation specificity of MsCHS towards a resorcinol product^44^ but there are no literature reports on its role in substrate specificity. Within our set of fungal T3PKS, this position is occupied by a negatively charged glutamate residue in the enzymes stemming from Sordariomycetes, as well as FerePKS, PtriPKS, and the inactive AtamPKS2 (Supplementary Fig. 10). Functionally, these enzymes appear to prefer aliphatic starter units (**7**-**10**) and substrate **3**, although CgloPKS and FerePKS can also accept substrate **1**.

Overall, the descriptive ML model does not allow the immediate rationalisation of the individual contributions of the selected amino acid positions to substrate selection, since there might also be synergistic effects of these positions at play. It may, however, provide an interesting starting point for future enzyme engineering campaigns.

### Pyrone-, resorcinol- and quinolone-type polyketides in the activity profile of fungal T3PKSs

Until now, we have focussed solely on substrate specificity of the PKSs. However, during activity profiling, many enzymes exhibited remarkable product promiscuity and yielded up to four products from a single substrate (Supplementary Table 3). To determine the cyclisation mode and the ketide number of the fungal T3PKS products, we analysed representative reactions using high-resolution tandem mass spectrometry (HR-MS/MS) and compared fragmentation patterns with those of known T3PKS reaction products (Supplementary Fig. 11-39). We discriminated between different cyclisation modes by the characteristic *m/z* value of the ring fragment (125.0244 for pyrone, calculated for C_6_H_5_O_3_^-^; 123.0452 for resorcinol, calculated for C_7_H_7_O_2_^-^; and 153.0193 for phloroglucinol, calculated for C_7_H_5_O_4_^-^). The identity of triketide pyrones, where the ring fragment is typically absent, was confirmed by observing the mass difference of 1.979 in the precursor ion *m/z* compared to tetraketide resorcinols.

Figure 6a depicts the product profile of 37 PKSs. Lactonisation appeared to be the preferred cyclisation mechanism for ring-type substrates **1-4**, as well as short-chain (**5-7**) and branched-chain (**11**) fatty acyl substrates. Fragmentation patterns corresponding to resorcinol products were observed only with fatty acyl substrates **8-10**, and the propensity for aldol condensation increased with chain length. Reactions with **8-10** typically yielded a mixture of pyrone and resorcinol products (Fig. 6b-f). However, several enzymes exhibited stricter cyclisation specificity: AiizPKS, AlupPKS1, AcosPKS and AserPKS2 were selective for resorcinol products with substrates **9-10**, while AastPKS was the only enzyme that yielded pyrones even from the longest substrate in the panel (**10**). Within the T3PKS pairs stemming from the same organisms, HargPKS1 and XacuPKS1 were specific for lactonisation, while HargPKS2 and XacuPKS2 produced pentaketide resorcinols from **10**. We did not detect *m/z* values corresponding to the fragmentation of phloroglucinol-type products in any of the fungal PKS reactions.

**Figure 6:**
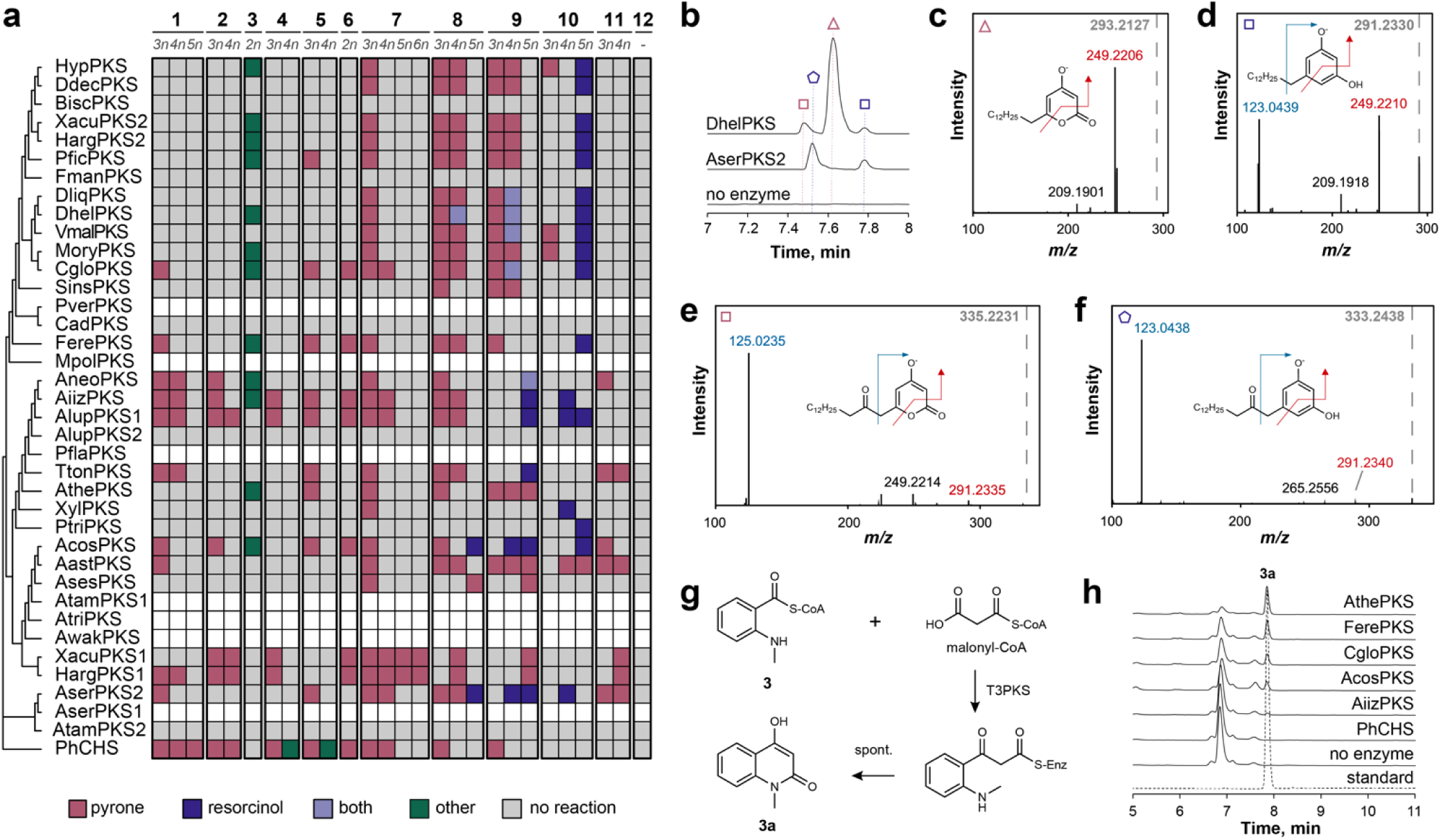
Product scope of the 37 fungal T3PKSs. a) Product profile of the reactions of cell-free lysates with substrates 1-12. Ketide number: 2n – diketide; 3n – triketide; 4n – tetraketide; 5n – pentaketide; 6n – hexaketide. b) Extracted ion chromatograms of the reaction products of DhelPKS and AserPKS2 with substrate 9 compared to the "no enzyme" control. Putative tri- and tetraketide pyrone products are marked with pink triangle and rectangle, respectively. Putative tetra- and pentaketide resorcinol products are marked with blue rectangle and pentagon, respectively. c-f) HR-MS/MS spectra and the proposed fragmentation patterns of the above-mentioned reaction products. The precursor ion is indicated with a dashed grey line labelled with its *m/z* value. g) The proposed^45^ scheme of diketide quinolone formation from 3. h) HPLC-based identification of reaction products of purified T3PKSs with 3. Dotted line represents the authentic standard of 4-hydroxy-1-methyl-2-quinolone. Detection was performed at 273 nm.

The *m/z* values of most products corresponded to tri-, tetra- and pentaketides, with HargPKS2 and XacuPKS2 also yielding a hexaketide pyrone from **7**. With substrate **3**, twelve of our enzymes produced a compound with an *m/z* of 174 in the negative ion mode, suggesting a diketide product (Fig. 6g). The UV absorbance spectrum of the compound with two local maxima at 273 nm and 315 nm suggested the formation of a fused ring structure of the quinolin-2-one type^46,47^ (Supplementary Fig. 15b). Indeed, its MS/MS fragmentation pattern (Supplementary Fig. 15c) was identical to that of 4-hydroxy- 1-methyl-2-quinolone produced by plant quinolone synthases^31^, and the authentic standard of 4- hydroxy-1-methyl-2-quinolone co-eluted with the reaction product peak (Fig. 6h). It is worth noting that PhCHS, our control enzyme, did not accept this substrate. Product *m/z* values from the ML validation experiment with substrates **13** to **24** corresponded to tri- and tetraketide pyrones of the respective carboxylic acids (Supplementary Fig. 8).

### Linking genomic context of the PKSs to their putative native products

Lastly, we examined the underlying biosynthetic gene clusters (BGCs) of the genes encoding the T3PKSs tested here to gain insight into their possible native functions. Overall, the immediate neighbourhood of the T3PKS genes is well conserved across most SSN clusters (Supplementary Data 7). However, in some SSN clusters, we observed several distinct groups of BGCs (clusters 1-3 and 1-4) or little conservation in the gene neighbourhood (cluster 15), suggesting that the SSN clusters are often but not always representative of functional similarity.

As expected, T3PKS genes are often neighboured by genes encoding typical tailoring functions – cytochrome p450s, aldo-keto reductases and short-chain dehydrogenases. Furthermore, T3PKS genes from clusters 9, 10, 12, 22 and 23 originate from hybrid biosynthetic gene clusters that also encode a T1PKS (AlupPKS2, AiizPKS, XylPKS, AthePKS), a non-ribosomal peptide synthetase (TtonPKS), or a dimethylallyltryptophane synthase (AiizPKS, AlupPKS1). While the T3PKSs encoded by some of these genes were among the most promiscuous of the set, accepting 6 to 10 substrates, XylPKS and AlupPKS2 hardly accepted any. The co-localisation of their genes with one encoding a putative highly reducing T1PKS might offer an explanation. In two of such hybrid fungal clusters characterised to date, the T3PKS accepts the product of the T1PKS – a polyunsaturated and/or branched fatty acid^19,21^. Through heterologous expression and gene knockout experiments, it was shown that the T3PKSs from these two clusters, FscB and PspB, do not accept endogenous fatty acyl-CoAs and only yield product when co-expressed with their partner T1PKSs^19,21^. XylPKS/FscB and AlupPKS2/PspB share 77% and 95% amino acid sequence identity, respectively. Enzymes encoded by the genes neighbouring the XylPKS- and AlupPKS2-encoding genes also share high sequence identity with those in the scirpilin and soppiline BGCs, respectively, suggesting chemically similar products (Fig. 7a-b). Genomic regions encoding for AthePKS and TtonPKS and their adjacent T1PKS do not share homology with any known BGC.

**Figure 7:**
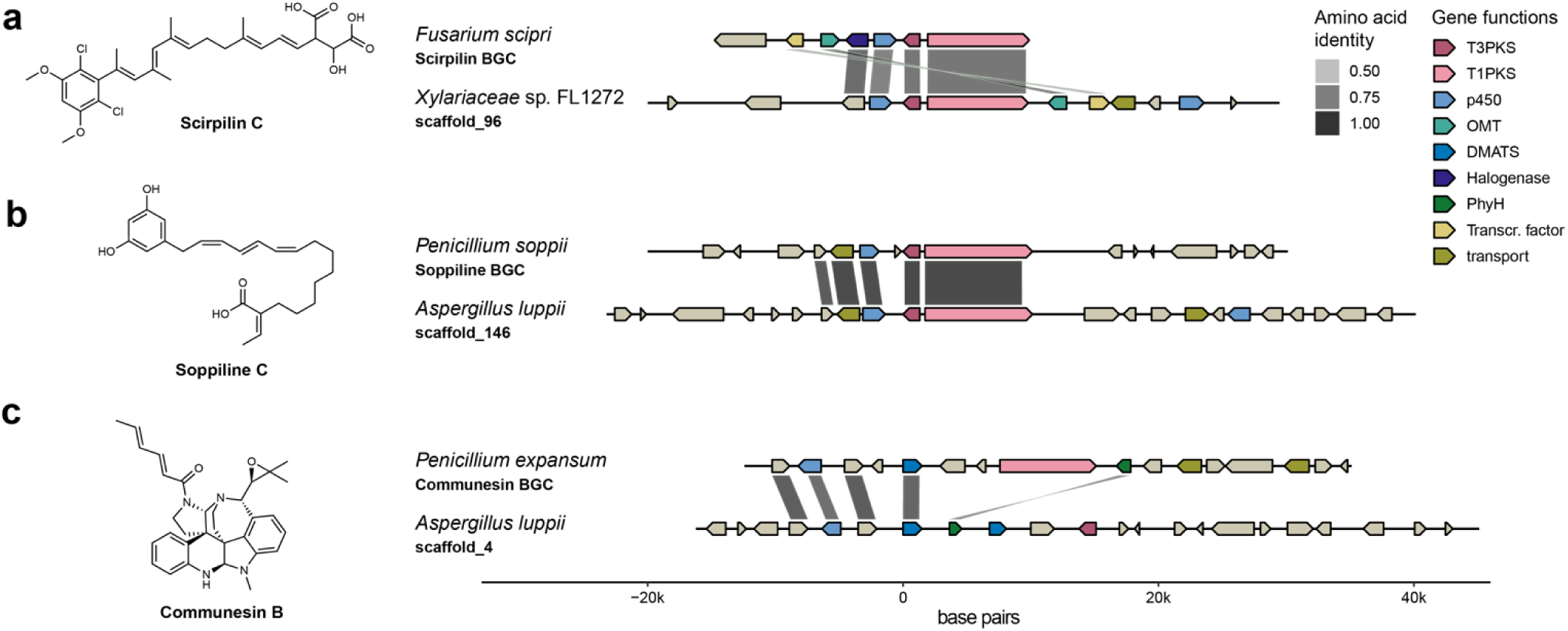
G**e**nomic **neighbourhood of the fungal T3PKSs.** a) Comparison of the scirpilin biosynthetic gene cluster in *Fusarium scirpi* (GenBank: OR551244) and the genomic neighbourhood of the XylPKS-encoding gene in *Xylariaceae sp.* FL1272. b) Comparison of soppiline biosynthetic gene cluster in *Penicillium soppii* (MIBiG^48^: BGC0003032) and the genomic neighbourhood of the AlupPKS2- encoding gene in *Aspergillus luppii*. c) Comparison of the communesin biosynthetic gene cluster in *Penicillium expansum* (MIBiG: BGC0001205) and the genomic neighbourhood of the AlupPKS1-encoding gene in *Aspergillus luppii*. The gene clusters and pairwise amino acid sequence identities between the encoded proteins were visualised using gggenomes^49^.

AiizPKS- and AlupPKS1-genes neighbour one or two dimethylallyltryptophane synthase (DMATS) genes, respectively. These enzymes typically catalyse the prenylation of tryptophan in alkaloid biosynthesis, but some are also active on tyrosine and aromatic polyketides^50^. To our knowledge, co-localisation of DMATS and T3PKS genes has not been observed before and may suggest a hybrid polyketide-indole or an aromatic meroterpenoid product. When examining the gene neighbourhood of the AlupPKS1-encoding gene, we noticed similarities in its gene composition with the BGC of communesin B, an anticancer alkaloid from *Penicillium expansum*^51^ (Fig. 7c). The major difference is that the AlupBGC encodes a T3PKS instead of a T1PKS, and two instead of one DMATS-type enzymes. Given the relaxed substrate specificity of AlupPKS1, functional studies at the BGC level are needed to shed light on the hybrid product encoded here.

Curiously, T3PKS genes from Sordariomycetes (SSN clusters 1, 3, 13 and 19) have fewer typical tailoring genes in their vicinity, but are frequently neighboured by genes encoding putative carbohydrate-active enzymes (CAZymes), such as glycosyl hydrolases and pectate lyases (Supplementary Fig. 40). Many fungal CAZymes are pathogenicity factors^52^, and the presence of a chalcone synthase-type gene was recently identified as one of the genetic determinants of endophytism in *Arabidopsis* mycobiota^53^. Thus, the observed co-localisation may hint at a link between T3PKSs and fungi-plant interactions.

## Discussion

Biocatalysis holds promise to reshape our approach to chemical and pharmaceutical synthesis, but its wider implementation is limited by the number of characterised enzymes at our disposal. In this study, we functionally characterised 37 new T3PKSs, essentially tripling the number of previously studied fungal enzymes in this family. Roughly half of the 24 CoA thioesters used here as substrates had not been tested with fungal – or any – T3PKSs before. As a result, we achieved the production of unnatural polyketides with pharmaceutically valuable alkyne^54^, furan^55^ and thiazole^56^ moieties, as well as several precursors of bioactive natural products. For instance, TtonPKS converted **20** to the direct precursor of kavain, an anticonvulsant from the kava plant, and half of the active PKSs converted derivatives of anthranilic acid to heterocyclic quinolin-2-one alkaloids. The ability to synthesise these biologically active scaffolds was thus far believed to be confined to a small group of divergent T3PKSs from Rutaceous plants^31,45,46^ and PqsD from the biosynthetic pathway of the *Pseudomonas* quinolone signal, PQS^57^. This unexpected insight into substrate and product specificities of fungal T3PKSs opens new possibilities for their recruitment in biocatalytic routes towards pharmaceutically relevant compounds.

In combination with high-throughput enzyme assays, machine learning is gaining traction as a powerful tool for substrate specificity prediction, with several domain-specific^58^, enzyme family-specific^59–62^ and general^63^ models reported recently. Building on our activity profiling data, we trained the first machine learning model for T3PKS substrate specificity prediction, which achieved reasonable accuracy despite the relatively modest size of our dataset (341 data points). This could be attributed to thorough sampling from the whole fungal T3PKS sequence space and a panel of structurally diverse substrates resulting in a balanced training dataset. Although predictive ML models tend to underperform with small molecules that were not present in the training dataset^63^, we found that our model performs well with an extended panel of substrates that are relatively distant from those in the training dataset. We anticipate that this feature can be valuable for narrowing down the number of enzymes to be screened for activity with a given substrate, especially in cases that are counterintuitive or hard to predict with human intelligence alone. For instance, the model correctly predicted that several enzymes would accept the bulky, nonplanar substrate **19** despite being inactive with simpler aromatic substrates **1** or **2**. A current limitation of our model, however, is the fungi-focussed nature of the training dataset. While it adequately predicts the activity of 11 published fungal T3PKSs, its performance declines dramatically with plant and bacterial enzymes. A unified model for phylogenetics-independent prediction of T3PKS substrate specificity would thus be a logical follow-up to this work.

Furthermore, it must be noted that our choice of substrates for activity mapping was inspired by literature precedent and limited by commercial and enzymatic accessibility of CoA thioesters. The physiological substrates (and, by extension, roles) of fungal T3PKSs remain to be uncovered. In our study, most enzymes produced resorcinol-type phenolic lipids from medium- to long-chain fatty acyl substrates *in vitro*. Reports of phenolic lipids from fungi include hydroxylated and sulfoalkylresorcinols from Xylariales^64,65^, glycosylated alkylresorcinols and resorcylic acids from Eurotiales^66,67^, and monounsaturated resorcylic acids from several basidiomycete fungi^68^. Adipostatin A, an alkylresorcinol from grain crops with reported fungistatic properties, has also been isolated from several species of plant-pathogenic fungi^69,70^. Consequently, it has been postulated that subtoxic levels of adipostatin A produced by these fungi might play a role in their higher tolerance to plant alkylresorcinols^69,71^. The Sordariomycete clade of alkylresorcinol-producing T3PKSs (SSN clusters 1, 3, 13 and 19) stems predominantly from endophytic fungi. It is thus conceivable that a similar resistance mechanism is also used by non-pathogenic endophytes to evade the plant defence system. Co-localisation of the genes encoding these T3PKSs with putative CAZyme genes further calls for investigation of their role in fungi-plant interactions.

Unprecedentedly for fungal T3PKSs, several enzymes from our set favoured lactonisation over aldol condensation even with longer fatty acyl substrates. We also observed functional differences between pairs of T3PKSs from the same organism: HargPKS1 and XacuPKS1 produced strictly pyrones from all accepted substrates, while HargPKS2 and XacuPKS2, like most other enzymes, yielded pentaketide resorcinols from **10**. A similar pattern has been observed with bacterial T3PKSs ArsB and ArsC from *Azotobacter vinelandii*: the former is specific for aldol condensation, while the latter prefers lactonisation^72^. However, it is also possible that the pyrones are shunt products caused by the feeding of non-physiological substrates, as commonly observed with plant chalcone synthases *in vitro*^73^. The true substrates of the PKSs tested here may be encoded within the underlying BGCs: either through the action of a dedicated CoA ligase on an exotic carboxylic acid substrate, or as a product of the partner T1PKS in hybrid clusters. The latter might explain why several PKSs encoded in hybrid clusters were expressed, but were not active on any substrate. On the other hand, the genes of some of the most promiscuous enzymes in our study co-localise with genes encoding other core biosynthetic enzymes – NRPS or DMATS. The natural product scaffolds produced by these unprecedented hybrid BGCs remain to be explored.

This study was conceived amid the controversy regarding flavonoid production in fungi. Although flavonoids have been reported in fungal extracts^74–76^, no dedicated routes for their biosynthesis had been described. While ORAS and AnPKS failed to produce naringenin chalcone from coumaroyl-CoA and malonyl-CoA *in vitro*^11,12^, the studies on other fungal T3PKSs did not include flavonoid precursors in the substrate panel. We therefore hypothesised that a comprehensive search across the fungal sequence space might uncover the enzymes responsible for flavonoid production. However, our kingdom-wide activity profiling did not yield proof for the formation of phloroglucinol products with any of the substrates, suggesting that fungal T3PKSs are unlikely to synthesise flavonoid scaffolds. This conclusion is corroborated by the recent discovery of fungal NRPS-PKS hybrids that produce flavonoids naringenin^77^ and chlorflavonin^78^. On the other hand, half of the active T3PKSs in this study produced plant-like quinolone alkaloids from anthranilic acid derivatives. While fungal 6,6-quinolones are known to originate from an NRPS pathway^79^, the potential involvement of T3PKSs in the synthesis of simple fungal 2-quinolones, such as penicinolone^80^, remains to be investigated.

In summary, we mined, expressed and characterised 37 new fungal T3PKSs in terms of their substrate and product scope. We then used the biochemical data to train and experimentally validate the first machine learning model for predicting T3PKS substrate specificity. We also trained a descriptive machine learning model that identified several residues as important for substrate specificity that we would not have labelled as such based on literature precedent or multiple sequence alignment. Therefore, this analysis may guide future efforts to elucidate the mechanism underlying substrate selection and to shift the substrate specificity by enzyme engineering. From the biocatalytic perspective, our approach identified highly promiscuous enzymes that enabled precursor-directed biosynthesis of several pharmaceutical precursors and unnatural polyketides. Future experimental efforts will focus on harnessing this promiscuity for chemoenzymatic and *in vivo* biosynthetic applications and elucidating the native functions of these enzymes.

## Methods

### Materials

CoA thioesters for T3PKS assays were purchased from Sanbio B.V. (Uden, The Netherlands), TransMIT (Gießen, Germany) or Endotherm Life Science Molecules (Saarbrücken, Germany). Malonyl-CoA, CoA and carboxylic acids for CoA ligation were purchased from Sigma-Aldrich (St. Louis, USA) or BLD Pharm (Shanghai, China). The myTXTL® kit for cell-free expression was obtained from Daicel Arbor Biosciences (Ann Arbor, USA). Linear gene fragments used as templates for cell-free expression were ordered as gBlocks™ from Integrated DNA Technologies, Inc. (IDT; Coralville, USA).

### Selection of enzymes of interest and bioinformatic analyses

Two different multiple sequence alignments (MSAs) of known T3PKSs from fungi and plants (Supplementary Data 1) were generated with mafft v7.505^81^ (parameters: --thread 8 --maxiterate 1000 --genafpair --reorder) and converted to Hidden Markov Model (HMM) profiles using the msa2profile function of MMseqs2^82^. The resulting HMM profiles were used to query the nucleotide sequences of 2096 fungal genomes from the Joint Genome Institute (JGI) MycoCosm repository^24^ using the easy-search function of MMseqs2 (parameters: -s 7). Hits representing exons of the same open reading frame were concatenated and converted to amino acid sequences. The hits were combined with 613 fungal proteins annotated as belonging to the IPR011141 family of the InterPro database^25^. The combined and dereplicated set of 1640 sequences was used as input for calculating a sequence similarity network with a webtool of the Enzyme Function Initiative (EFI-EST)^27^; edge selection cut-off: alignment score threshold 40. To reduce complexity of the sequence similarity network (SSN), we opted for the "representative node" mode, where sequences sharing 95% sequence identity are grouped and visualised as a single node. The finalised network was visualised in Cytoscape 3.8.2^83^ with the yFiles organic layout. The sequences were filtered by length (300 to 600 amino acids) to exclude fragments, and the final sequence identity cutoff was set to 80%.

Sequences from each cluster were retrieved and aligned with a set of reference T3PKSs from plants (Supplementary Data 1) using mafft (parameters: --genafpair). Residues from positions 132, 133, 164, 194, 197, 215, 256, 265, 303, 336, 338 and 375 (residue numbering from MsCHS, Uniprot: P30074) were extracted and compared across fungal and plant T3PKSs. An MSA of all fungal T3PKSs and 184 reviewed plant T3PKSs from the InterPro database was performed in the same way and visualised as a sequence logo using the R package ggseqlogo^25^. From the thus analysed clusters, enzymes were chosen for experimental characterisation (Supplementary Table 1). In total, 25 clusters representing taxonomic and ecological diversity of the host organisms, as well as the putative functional diversity hinted at by the active site composition, were selected. For sequence selection within the clusters, we prioritised those with active site residues corresponding to the cluster consensus and those stemming from highly contiguous, high-quality genome assemblies.

To examine the T3PKS-encoding gene neighbourhoods within each SSN cluster, 50,000 base pairs both up- and downstream of the T3PKS-encoding gene were extracted from the corresponding genome using the subseq function of seqkit^84^ (parameters: -d 50000 -u 50000). The neighbourhoods were visualised using the R package gggenomes v0.9.12.9000^49^. For biosynthetic gene cluster comparisons, the BGCs were first predicted with the command line version of antiSMASH 7.0.0^85^ (parameters: -- taxon fungi --cassis --clusterhmmer --genefinding-gff3 --genefinding-tool none). BGC comparison and visualisation was performed using gggenomes v0.9.12.9000^49^.

### Design and synthesis of DNA templates for cell-free expression

The selected enzymes were back-translated into DNA sequences, codon-optimised for expression in *E. coli* K12 with the IDT optimisation algorithm and manually modified to exclude recognition sites for NcoI, HindIII and XhoI restriction enzymes. The 5’ end of all DNA fragments was designed with an identical overhang of 100 bp, the p70a promoter sequence, a ribosome-binding site and an NcoI recognition site to facilitate cloning into the pET28a(+) (Novagen) expression vector. The 3’ end of each gene was designed to include the recognition site for HindIII, the in-frame codons for the Tobacco Edge Virus protease cleavage site, a hexahistidine-tag, and a GFP11 tag. After the stop codon, each gene fragment included an XhoI recognition site, the T500 terminator sequence and a 100 bp overhang. The synthetic DNA was obtained as gBlocks Gene Fragments from IDT. The sequences of all synthetic DNA constructs are provided in Supplementary Data 5, and gene fragment annotations are included in Supplementary Data 6.

### Construction of plasmids

Genes coding for T3PKSs selected for expression in *E. coli* were amplified by polymerase chain reaction (PCR) from the synthetic DNA fragments and cloned by restriction and ligation (NcoI/HindIII) into pET28a(+) for expression under the T7 promoter. The gene encoding the standard protein PhCHS- GFP11 for split-GFP assays was cloned into pET28a(+) by restriction and ligation with NcoI/XhoI to retain the GFP11 tag. All constructs were verified by Sanger sequencing (Macrogen Europe, Amsterdam). The list of all plasmids used in this study is provided in Supplementary Table 4.

### Cell-free expression

Cell-free expression was performed with the myTXTL® Linear DNA kit from Daicel Arbor Biosciences according to the manufacturer’s instructions. Briefly, the synthesised DNA fragments were reconstituted in ultrapure water to a final concentration of 80 nM and stored at -20 °C between experiments. myTXTL® lysate was thawed on ice and aliquoted at 9 µL into 1.5 mL tubes or 96 well plates. 3 µL of DNA template were added to each tube or well and mixed by pipetting (final concentration 20 nM). The reactions were briefly centrifuged to collect the liquid at the bottom and transferred to an incubator at 29 °C for 20 h. As a negative control (’no enzyme’), one reaction was performed with 3 µL of ultrapure water instead of DNA. After incubation, the reactions were diluted 2.5-fold with ultrapure water and used directly for the split-GFP and T3PKS assays.

### Split-GFP complementation assay

T3PKS expression in the cell-free system was quantified using a split-GFP complementation assay. The GFP fluorescence complementation fragment GFP1–10 was expressed in *E. coli* BL21(DE3) from the plasmid pAGM22082_sfGFP1-10^36^, prepared as inclusion body pellets according to a published protocol^86^ and stored at –70 °C between experiments. For the measurement, a pellet of GFP1–10 was fully dissolved in 9 M urea and resuspended in 25 mL of TNG buffer (100 mM Tris-HCl pH 7.4, 100 mM NaCl, 10% (v/v) glycerol). 2.5 µL of the diluted myTXTL® reactions were aliquoted into a 96-well µClear® white plate (Greiner Bio-One, Kremsmünster, Austria) and mixed with 7.5 µL of TNG buffer. In parallel, serial dilutions of the standard protein (purified PhCHS-GFP11, see section “Expression and purification of T3PKSs”) were aliquoted in triplicate into the same plate. 90 µL of the GFP1–10 solution was then added to each well. Immediate fluorescence values were measured using a FLUOstar Omega (BMG LABTECH, Ortenberg, Germany) microplate reader (excitation wavelength: 485 nm; emission wavelength: 520 nm; bottom read mode). The plate was incubated at 4 °C overnight, and the final fluorescence values were measured with the same parameters. Complementation fluorescence ΔF was calculated from equation 1,

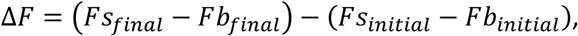

where Fs_final_ and Fs_initial_ are the final and initial fluorescence values for a sample, and Fb_final_ and Fb_initial_ are the final and initial fluorescence values for the blank (myTXTL + ultrapure water). ΔF values were converted to protein concentrations in μM based on the calibration curve with the standard protein.

## *In vitro* T3PKS assays

The standard T3PKS reaction mixture consisted of 0.3 mM starter-CoA, 0.3 mM extender-CoA (malonyl-CoA) and 5 µL of the diluted myTXTL® lysate or purified enzyme (final concentration 3 µM) in 50 mM Tris/HCl pH 7.5. The total reaction volume was 50 µL, and the reaction was initiated by the addition of the enzyme. After 24 h of incubation at 30 °C, 300 µL of 9:1 ethyl acetate-methanol (v/v) supplemented with 0.1% formic acid were added and the plates were shaken for 20 min. After centrifuging at 3,428 × g for 15 min, 250 µL of the organic layer were transferred to another plate with a JANUS liquid handler (PerkinElmer, Waltham, United States) and dried under a stream of N_2_. The dried extracts were resuspended in 25 µL methanol supplemented with 0.1% formic acid and used for chromatographic analysis. Reactions with purified PhCHS or ultrapure water served as the positive and negative controls, respectively.

### Analysis of T3PKS reaction products

The extracted PKS reactions were first analysed using a low-resolution Waters Acquity Arc UHPLC-MS system equipped with a 2998 PDA detector and a QDa single-quadrupole mass detector. The samples were separated over a Waters XBridge BEH C18 3.5 μm 2.1×50 mm column at 40 °C with a concentration gradient (solvent A: water +0.1 % formic acid, and solvent B: acetonitrile + 0.1 % formic acid) at a flow rate of 0.5 mL/min (2 μL injections). The following gradient was used: 5 % B for 2 min, 5–90 % B over 3 min; 90 % B for 2 min; 5 % B for 3 min. For reactions with oleoyl-CoA and phytanoyl-CoA, a different gradient was employed: 50 % B for 2 min, 50–70 % B over 4 min; 70–90 % B over 2 min; 90 % B for 2 min; 90–50% B over 2 min; 50 % B over 2 min. MS analysis was carried out in both positive and negative ion modes with the following parameters: probe temperature of 600 °C; capillary voltage of 1.0 kV; cone voltage of 15 V; scan range 100-1250 *m/z*. The acquired data were analysed using the proprietary software MassLynx.

High-resolution and MS/MS analyses of the extracted PKS reactions were performed using a Shimadzu LC20-XR system (Shimadzu Benelux, Den Bosch, The Netherlands) coupled to a Q Exactive Plus mass spectrometer (Thermo Fisher Scientific, USA). The samples were separated over a Waters XBridge BEH C18 reversed-phase column at 50 °C with a concentration gradient (solvent A: water + 0.1 % formic acid, and solvent B: acetonitrile + 0.1 % formic acid) at a flow rate of 0.5 mL/min (2 μL injections). The following gradient was used: 5 % B for 2 min, 5–90 % B over 3 min; 90 % B for 2 min; 5 % B for 3 min. MS and MS/MS analyses were performed with electrospray ionisation (ESI) in positive or negative ion mode at a spray voltage of 3.5, and sheath and auxiliary gas flow set at 48 and 11, respectively. The ion transfer tube temperature was 255 °C. Spectra were acquired in data-dependent mode with a survey scan at m/z 80 − 1200 at a resolution of 70,000 followed by MS/MS fragmentation of the top 5 precursor ions at a resolution of 17,500. A stepped collision energy of 30-40-55 was used for fragmentation, and fragmented precursor ions were dynamically excluded for 10 s. The acquired data were analysed using MZmine 4.0.3^87^.

### CoA ligation reactions

Os4CL from *Oryza sativa*^40^ or PqsA from *Pseudomonas aeruginosa*^41^ were expressed and purified according to the original protocols and used for CoA ligation reactions with carboxylic acid substrates. The reaction mixture consisted of 5 mM MgCl_2_, 6.25 mM ATP, 1 mM carboxylic acid, 1.5 mM CoA and 10 mM enzyme in 100 mM Tris/HCl pH 7.5. The total volume of the reaction was 100 µL or 1,000 µL for analytical and preparative scale, respectively. The reactions were incubated at 30 °C for 16 h and analysed by HPLC as described by Rautengarten *et al.*^40^. The reactions were used directly for T3PKS assays or aliquoted and stored at -20 °C until use.

### Expression and purification of T3PKSs

Plasmids harbouring the T3PKS genes were transformed into chemically competent *E. coli* BL21(DE3) and maintained on selective LB agar containing 50 μg/ mL kanamycin. A starter culture was inoculated from a single colony (5 mL, LB with ampicillin) and incubated at 37 °C (180 rpm, overnight). The main culture was inoculated from the starter culture (1 : 100) into 200 mL of TB autoinduction medium (20 g/L tryptone, 24 g/L yeast extract, 4 ml/L glycerol, 17 mM KH_2_PO_4_, 72 mM K_2_HPO_4_, 0.2 % w/v lactose and 0.05 % w/v glucose) and incubated at 37 °C (180 rpm, 2 h), after which the temperature was lowered to 18 °C (180 rpm, overnight). All following steps were performed with chilled buffers. The cells were harvested by centrifugation (15 min, 3,428 × g) and the pellet was resuspended in 5 volumes of lysis buffer (buffer A including one EDTA-free protease inhibitor tablet (Roche, Basel, Switzerland); buffer A: 50 mM Tris/HCl pH 7.5, 500 mM NaCl, 20 mM imidazole). The cell suspension was lysed by sonication (50 % duty cycle, 7 cycles of 35 s ON/60 s OFF) and cleared by centrifugation for 60 min at 17,000 × g. The supernatant was loaded onto the Ni-NTA affinity matrix (Qiagen, Hilden, Germany) equilibrated with buffer A by gravity flow. The column was washed with 20 column volumes of wash buffer (50 mM Tris/HCl pH 7.5, 500 mM NaCl, 30 mM imidazole) and eluted stepwise with one column volume of buffers B1 to B6 (buffers B1-B6: 50 mM Tris/HCl pH 7.4, 500 mM NaCl, 50 mM imidazole/ 100 mM imidazole/ 150 mM imidazole/ 200 mM imidazole/ 250 mM imidazole or 500 mM imidazole, respectively). The eluates of each step were collected in separate fractions and analysed by SDS-PAGE. T3PKS-containing fractions with low protein background were pooled and transferred into the storage buffer (10 mM HEPES/NaOH pH 7.5, 50 mM NaCl, 2 mM dithiothreitol, 5% (v/v) glycerol) by three cycles of concentration (Amicon® Ultra Centrifugal Filter; 3 kDa molecular weight cutoff) and dilution (1:30). The protein concentrations were determined by absorbance at 280 nm (NanoDrop, ThermoFisher Scientific, USA) before the purified enzymes were aliquoted and flash-frozen with liquid nitrogen for storage at −70 °C.

### Machine learning

To generate molecular descriptors for the polyketide starter units, the canonical SMILES for the acyl-moieties were retrieved from PubChem and three types of Extended-Connectivity Fingerprints (ECFPs) - Morgan Fingerprints (2048 bits), RDKit topological fingerprints and MACCS keys - were calculated using RDKit^88^. Embeddings for enzymes were generated from full-length sequences using the ProtT5- XL-UniRef50 model by ProtTrans^89^. ML models were trained using scikit-learn 1.5.0^90^ with default hyperparameters using 0.7/ 0.3 *StratifiedShuffleSplit* for training/ testing purposes. The model performance was assessed by calculating the average accuracy, area under the receiver operating characteristic curve (AUROC), recall, precision, and F1 score in 100 runs with a random seed.

For building a descriptive ML model, the crystal structures of ORAS (type III pentaketide synthase from *Neurospora crassa,* PDB: 3EUT) and MsCHS (chalcone synthase from *Medicago sativa* (PDB: 1CGK) were used as templates. Residues within 5 Å of eicosanoic acid in 3EUT and within 6 Å of the catalytic cysteine and naringenin in 1CGK were selected, and the corresponding positions in the 31 active fungal T3PKSs were retrieved from an MSA. 12 fully conserved positions were removed, and 88 molecular descriptors were calculated using the peptides.py package – a python realisation of the Peptides R package^91^. The final feature vectors for each enzyme/ substrate pair were obtained by concatenating amino acid molecular descriptors with substrate MACCS Keys and had the following dimensions: {41 amino acids x (88 molecular descriptors) + 167 MACCS Keys}. The Decision tree model in scikit-learn 1.5.0^90^ was trained and assessed as described above for the predictive model. The model feature importances were extracted using the default *feature_importances_* function and summed up for each residue. The structure models of fungal T3PKSs were obtained using the local installation of AlphaFold 2^92^ at the Hábrók high performance computing cluster (University of Groningen). Data visualisation was performed using *ggplot2*^93^ *ComplexHeatmap*^94^ R packages, and PyMOL.

## Data availability

The data generated in this study are provided in the manuscript and in the Supplementary Information.

## Code availability

The code used to train machine learning models and visualise gene neighbourhoods of the T3PKSs can be found at: https://github.com/thackl/type_III_PKS and https://doi.org/10.5281/zenodo.14066149.

## Acknowledgements

This project has received funding from the European Union’s Horizon 2020 research and innovation programme under the Marie Skłodowska-Curie grant agreement 893122 to K.H. S.S.D was supported in part by Netherlands Organisation for Scientific Research Rubicon grant 019.202EN.038 and the European Union’s Horizon 2020 Marie Skłodowska-Curie grant 101152560.

pAGM22082_sfGFP1-10 was a gift from Sylvestre Marillonnet (Addgene plasmid #153515). We thank the Center for Information Technology of the University of Groningen for their support and for providing access to the Hábrók high performance computing cluster. The authors are grateful to the Interfaculty Mass Spectrometry Center of the University of Groningen and the University Medical Center Groningen for their services in high resolution tandem mass spectrometry, Ronald van Merkerk for help with establishing the robotic extraction system, and Dr. Angelina Osipyan for providing carboxylic acids of substrates **14** and **15**.

## Author contributions

N.S. and K.H. conceived the study. N.S. selected the putative T3PKSs with the help of T.H., and performed all biochemical experiments. S.S.D. trained the machine learning models. N.S. and S.S.D. analysed the data and created figures. T.H. provided bioinformatics support. N.S. wrote the manuscript with input from S.S.D. and guidance from K.H. All authors read and approved the final version of the manuscript.

## Materials & Correspondence

Correspondence and material requests should be addressed to Dr. Kristina Haslinger

