## Supplementary Figures and Tables for "Unravelling the functional diversity of type III polyketide synthases in fungi"

### Supplementary Tables

**Supplementary Table 1**. List of putative fungal T3PKSs selected for experimental characterisation

| Name | JGI accession | Uniprot | Organism | Genome reference |
|---|---|---|---|---|
| AastPKS | Aspaste1\|20971 | - | *Aspergillus astellatus* | * |
| AcosPKS | Aspcos1\|217105 | - | *Aspergillus costaricaensis* CBS 115574 | 1 |
| AiizPKS | Aspiiz1\|328085 | - | *Aspergillus iizukae* CBS 541.69 | * |
| AlupPKS1 | Asplup1\|182290 | - | Aspergillus luppii CBS 653.74 | * |
| AlupPKS2 | Asplup1\|227097 | - | *Aspergillus luppii* CBS 653.74 | * |
| AneoPKS | Aspneof1\|275638 | - | *Aspergillus neoflavipes* CBS 260.73 | * |
| AserPKS1 | Aspser1\|206060 | - | *Aspergillus sergii* CBS 130017 | 2 |
| AserPKS2 | Aspser1\|218351 | - | *Aspergillus sergii* CBS 130017 | 2 |
| AsesPKS | Aspses1\|335127 | - | *Aspergillus sesamicola* CBS 137324 | * |
| AtamPKS1 | Asptam1\|307157 | A0A5N6VBQ0 | *Aspergillus tamarii* CBS 117626 | 2 |
| AtamPKS2 | Asptam1\|310896 | A0A5N6V0I9 | *Aspergillus tamarii* CBS 117626 | 2 |
| AthePKS | Aspth1\|366720 | - | *Aspergillus thesauricus* IBT 34227 | * |
| AtriPKS | Asptri1\|208758 | - | *Aspergillus trinidadensis* IBT 32571 | * |
| AwakPKS | Aspwak1\|463812 | - | *Aspergillus waksmanii* IBT 31900 | * |
| BiscPKS | Biscog1\|570470 | - | *Biscogniauxia* sp. FL1348 | 3 |
| CadPKS | Cadsp1\|525675 | A0A2V1CPT9 | *Cadophora* sp. DSE1049 | 4 |
| CglobPKS | Chagl1\|381215 | - | *Chaetomium tenue* MPI-SDFR-AT-0079 v1.0 | 5 |
| DdecPKS | Daldec1\|343198 | - | *Daldinia decipiens* CBS 113046 | 3 |
| DhelPKS | Diahe1\|3921 | A0A2P5I7D7 | *Diaporthe helianthi* str. 7/96 | 6 |
| DliqPKS | - | A0A7T8G346 | *Diaporthe liquidambaris* | - |
| FerePKS | Foner1\|4724 | A0A178ZV18 | *Fonsecaea erecta* CBS 125763 | 7 |
| FmanPKS | Fusma1\|10567 | - | *Fusarium mangiferae* MRC7560 | 8 |
| HargPKS1 | Hyparg1\|454749 | - | *Hypoxylon argillaceum* CBS 527.63 | 3 |
| HargPKS2 | Hyparg1\|272596 | - | *Hypoxylon argillaceum* CBS 527.63 | 3 |
| HypPKS | HyNC0597_1\|672645 | - | *Hypoxylon* sp. NC0597 | 3 |
| MoryPKS | Magor1\|2848 | G4MSX3 | *Pyricularia oryzae* 70-15 | 9 |
| MpolPKS | Mycpol1\|1099419 | - | *Mycena polygramma* CBHHK137 | 10 |
| PficPKS | Pesfi1\|10058 | W3WYM6 | *Pestalotiopsis fici* W106-1 | 11 |
| PflaPKS | Penfla1\|6890 | A0A1V6SK38 | *Penicillium flavigenum* IBT 14082 | 12 |
| PtriPKS | - | A0A2W1FIY5 | *Pyrenophora tritici-repentis* | 13 |
| PverPKS | Psever1\|4162 | A0A1B8GPR4 | *Pseudogymnoascus verrucosus* UAMH 10579 | 14 |
| SinsPKS | Spoin1\|227 | A0A162JF80 | *Sporothrix insectorum* RCEF 264 | 15 |
| TtonPKS | Trito1\|844 | - | *Trichophyton tonsurans* CBS 112818 | 16 |
| VmalPKS | - | A0A194VDY8 | *Valsa mali* var. pyri | 17 |
| XacuPKS1 | Xylacu1\|452307 | - | *Xylaria acuta* CBS 122032 | 3 |
| XacuPKS2 | Xylacu1\|516844 | - | *Xylaria acuta* CBS 122032 | 3 |
| XylPKS | XyFL1272_2\|442241 | - | *Xylariaceae* sp. FL1272 | 3 |

* = these whole genome sequencing projects were executed by the US Department of Energy Joint Genome Institute https://www.jgi.doe.gov/ in collaboration with the user community.

**Supplementary Table 2**. List of T3PKSs from fungi, plants and bacteria with published activity data that were used in the ML validation experiments in this study.

| Enzyme | Origin | Donor organism | Reference |
|---|---|---|---|
| ORAS | fungal | *Neurospora crassa* | 18 |
| AnPKS | fungal | *Aspergillus niger CBS 513.88* | 19 |
| An-CsyA | fungal | *Aspergillus niger NRRL 328* | 20 |
| CsyA | fungal | *Aspergillus oryzae* | 21 |
| BPKS | fungal | *Botrytis cinerea* | 22 |
| Sl-PKS2 | fungal | *Sporotrichum laxum* | 23 |
| SmPKS | fungal | *Sordaria macrospora* | 24 |
| CtPKS | fungal | *Chaetomium thermophilum* | 24 |
| CsyB | fungal | *Aspergillus oryzae* | 24 |
| NuPKS | fungal | *Naganishia uzbekistanensis* | 25 |
| FiPKS | fungal | *Fusarium incarnatum* | 26 |
| QNS-Marmelos | plant | *Aegle marmelos* | 27 |
| QNS-Microcarpa | plant | *Citrus microcarpa* | 28 |
| ANS-Microcarpa | plant | *Citrus microcarpa* | 28 |
| BNS-Palmatum | plant | *Rheum palmatum* | 29–31 |
| ARAS1 | plant | *Oryza sativa* | 32 |
| ARAS2 | plant | *Oryza sativa* | 32 |
| Sg-RppA | bacterial | *Streptomyces griseus* | 33 |
| ArsB | bacterial | *Azotobacter vinelandii* | 34 |
| ArsC | bacterial | *Azotobacter vinelandii* | 34 |
| Ncs | bacterial | *Streptomyces clavuligerus* | 35 |
| BpsA | bacterial | *Bacillus subtilis* | 36 |
| PhlD | bacterial | *Pseudomonas fluorescens* | 37 |
| gcs | bacterial | *Streptomyces coelicolor* | 38 |
| Se-RppA | bacterial | *Saccharopolyspora erythraea* | 39 |
| Sc-RppA | bacterial | *Streptomyces coelicolor* | 40 |

**Supplementary Table 3**. Low-resolution LC-MS analysis of the T3PKS reaction products detected during activity profiling with substrates **1-12**. High-resolution LC-MS/MS analysis of the products is shown in Supplementary Figures 11-39.

| Starter substrate | Product | $t_R$, min | Ion mode | Detected *m/z* | Ketide number* | Cyclisation type* |
|---|---|---|---|---|---|---|
| benzoyl-CoA (**1**) | **1a** | 5.46 | ESI- | 187 | 3 | lactone |
| | **1b** | 5.36 | ESI- | 229 | 4 | lactone |
| phenylacetyl-CoA (**2**) | **2a** | 5.57 | ESI- | 201 | 3 | lactone |
| | **2b** | 5.47 | ESI- | 243 | 4 | lactone |
| N-methylanthraniloyl-CoA (**3**) | **3a** | 5.25 | ESI- | 174 | 2 | quinolone |
| p-coumaroyl-CoA (**4**) | **4a** | 5.25 | ESI- | 229 | 3 | lactone |
| | **4b** | 5.18; 5.61 | ESI- | 271 | 4 | phloroglucinol |
| β-methylcrotonoyl-CoA (**5**) | **5a** | 5.36 | ESI+ | 167 | 3 | lactone |
| | **5b** | 5.19 | ESI+ | 209 | 4 | lactone |
| acetyl-CoA (**6**) | **6a** | 0.97 | ESI+ | 127 | 3 | lactone |
| hexanoyl-CoA (**7**) | **7a** | 5.83 | ESI- | 181 | 3 | lactone |
| | **7b** | 5.76 | ESI- | 223 | 4 | lactone |
| | **7c** | 6.15 | ESI- | 265 | 5 | lactone |
| | **7d** | 5.41 | ESI- | 307 | 6 | lactone |
| decanoyl-CoA (**8**) | **8a** | 6.72 | ESI- | 237 | 3 | lactone |
| | **8b** | 6.64 | ESI- | 279 | 4 | lactone |
| | **8c** | 6.95 | ESI- | 321 | 5 | lactone |
| | **8d** | 6.89 | ESI- | 235 | 4 | resorcinol |
| | **8e** | 6.68 | ESI- | 277 | 5 | resorcinol |
| myristoyl-CoA (**9**) | **9a** | 7.62 | ESI- | 293 | 3 | lactone |
| | **9b** | 7.48 | ESI- | 335 | 4 | lactone |
| | **9c** | 7.80 | ESI- | 377 | 5 | lactone |
| | **9d** | 7.79 | ESI- | 291 | 4 | resorcinol |
| | **9e** | 7.52 | ESI- | 333 | 5 | resorcinol |
| oleoyl-CoA (**10**) | **10a** | 6.14 | ESI- | 347 | 3 | lactone |
| | **10b** | 5.86 | ESI+ | 391 | 4 | lactone |
| | **10c** | 6.57 | ESI+ | 433 | 5 | lactone |
| | **10d** | 6.52 | ESI+ | 347 | 4 | resorcinol |
| | **10e** | 5.96 | ESI- | 387 | 5 | resorcinol |
| phytanoyl-CoA (**11**) | **11a** | 7.38 | ESI- | 377 | 3 | lactone |
| | **11b** | 7.06 | ESI- | 419 | 4 | lactone |

$t_R$ = retention time; * = putative

**Supplementary Table 4**. List of plasmids used in this study.

| Construct name | Backbone | Enzyme encoded | Source |
|---|---|---|---|
| pET28a::6His-Os4CL | pET-28a(+) | Os4CL | [41] |
| pAGM22082_sfGFP1-10 | pAGM22082 | sfGFP1-10 | [42] |
| p70a::mcbR | p70a | mcbR | [43] |
| p70a::PhCHS-6His-GFP11 | p70a | PhCHS-GFP11 | This study |
| pET28a::PhCHS-6His-GFP11 | pET-28a(+) | PhCHS-GFP11 | This study |
| pET28a::AastPKS-6His | pET-28a(+) | AastPKS | This study |
| pET28a::AiizPKS-6His | pET-28a(+) | AiizPKS | This study |
| pET28a::AthePKS-6His | pET-28a(+) | AthePKS | This study |
| pET28a::CgloPKS-6His | pET-28a(+) | CgloPKS | This study |
| pET28a::DhelPKS-6His | pET-28a(+) | DhelPKS | This study |
| pET28a::FerePKS-6His | pET-28a(+) | FerePKS | This study |
| pET28a::HargPKS1-6His | pET-28a(+) | HargPKS1 | This study |
| pET28a::PficPKS-6His | pET-28a(+) | PficPKS | This study |
| pET28a::PhCHS-6His | pET-28a(+) | PhCHS | This study |
| pET28a::XacuPKS1-6His | pET-28a(+) | XacuPKS1 | This study |
| pET28a::AcosPKS-6His | pET-28a(+) | AcosPKS | This study |
| pET28a::TtonPKS-6His | pET-28a(+) | TtonPKS | This study |
| pET28a::HypPKS-6His | pET-28a(+) | HypPKS | This study |
| pET28a::6His-PqsA | pET-28a(+) | PqsA | This study |

#### Supplementary Figures

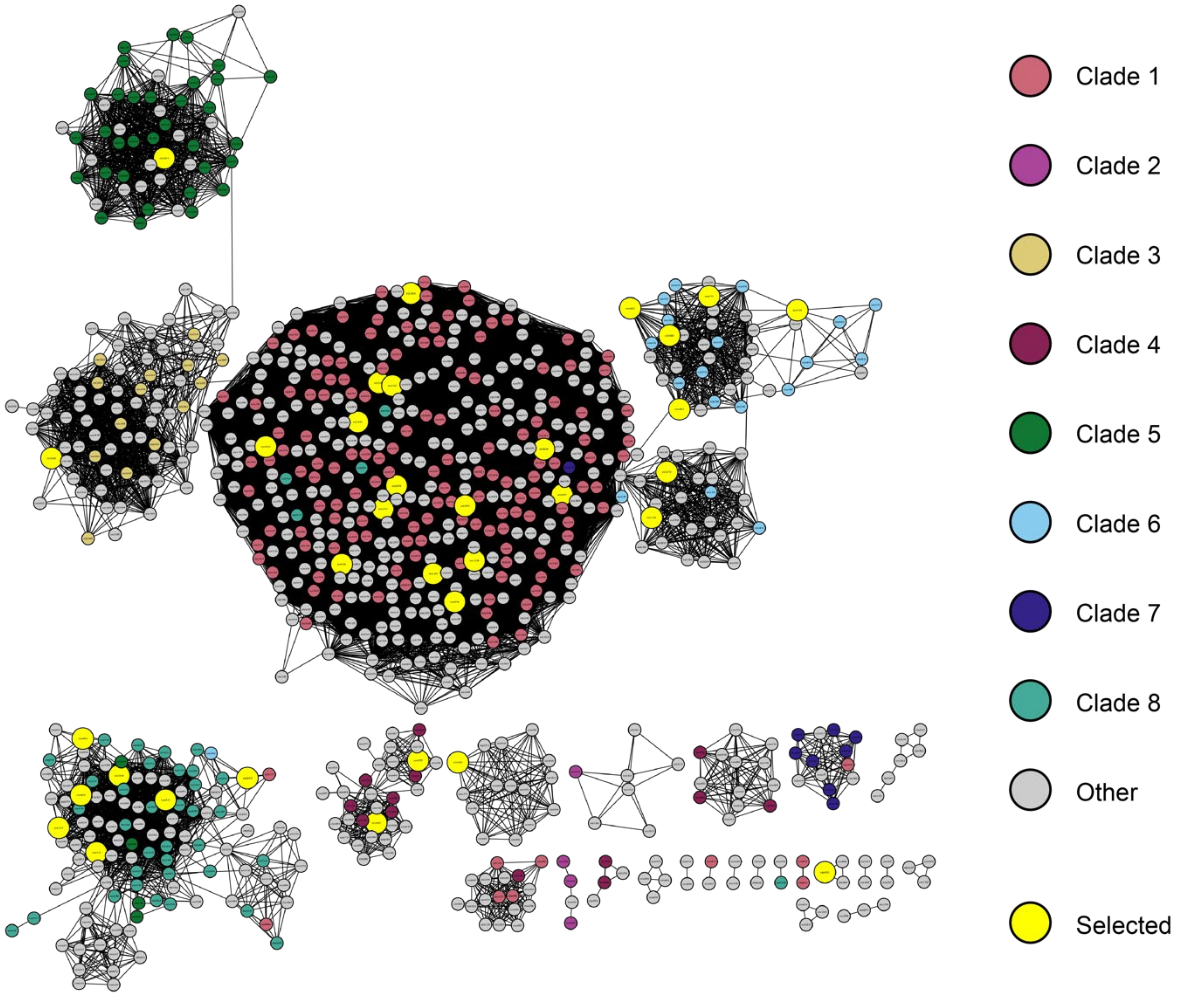

**Supplementary Figure 1:** A sequence similarity network of the putative fungal T3PKSs mined from the Mycocosm database at 57% sequence identity cutoff. Each circle is a representative node grouping protein sequences with >95% sequence identity. The colours reflect phylogenetic clades proposed by Navarro-Muños and Collemare in a recent evolutionary analysis[44].

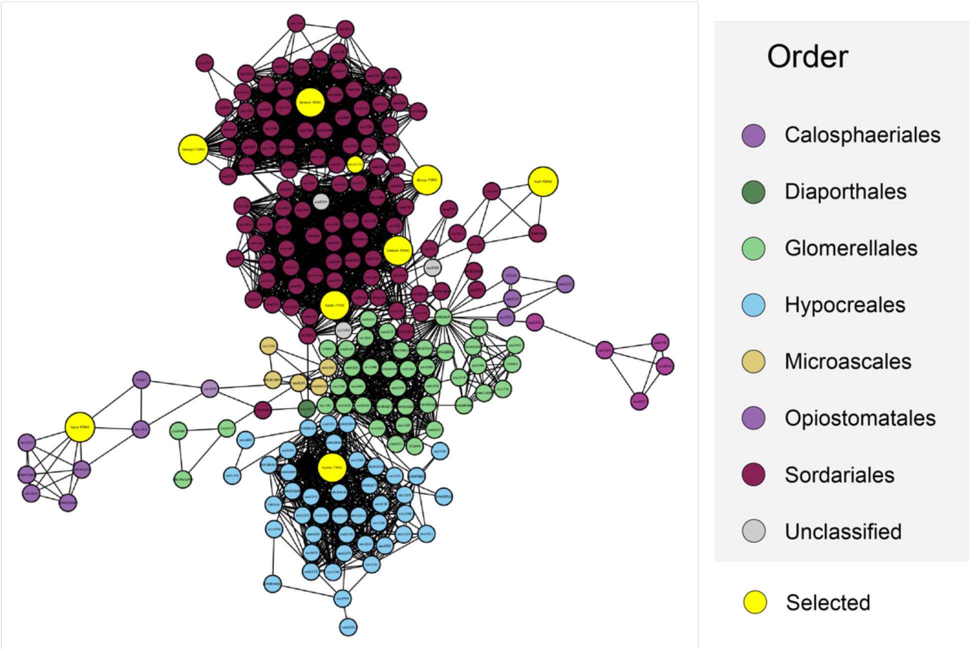

**Supplementary Figure 2:** Cluster 1 of the sequence similarity network of fungal T3PKSs. The nodes are coloured based on the taxonomic order of the donor organisms.

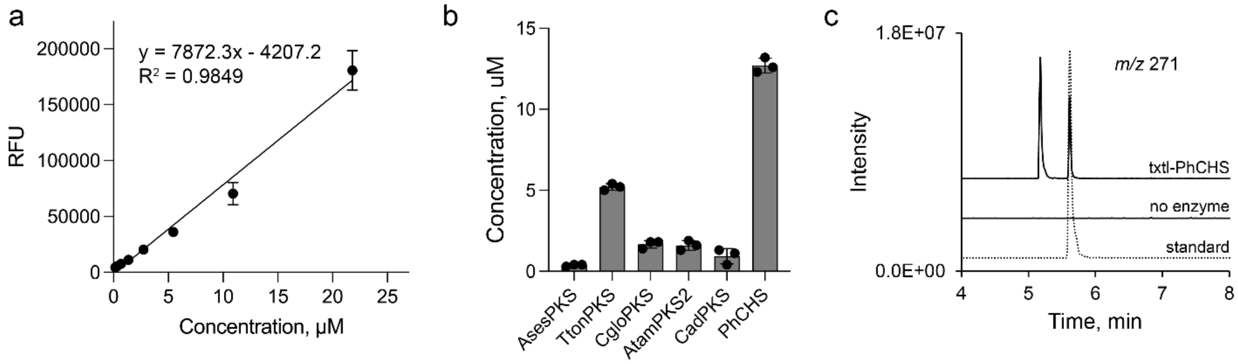

**Supplementary Figure 3.** Benchmarking of the express-test workflow with PhCHS. a) Calibration plot of the purified PhCHS-GFP11 fusion standard protein in TNG buffer used for quantifying soluble cell-free expressed protein; RFU - relative fluorescence units; data points and error bars reflect mean ± SD; n=3. b) Comparison of enzyme concentrations across different batches of cell-free expression reactions using PCR-amplified linear templates. Enzyme concentrations were determined using the split-GFP assay; histograms and error bars reflect mean ± SD; n=3. c) Low-resolution LC-MS analysis of the EtOAc extract of the enzymatic reaction of the cell-free expressed PhCHS (txtl-PhCHS) with substrate **4**; extracted ion chromatogram of the predicted *m/z* of naringenin chalcone and naringenin (271, negative mode). Y-axis shows relative ion abundance. Dotted line represents the authentic standard of naringenin.

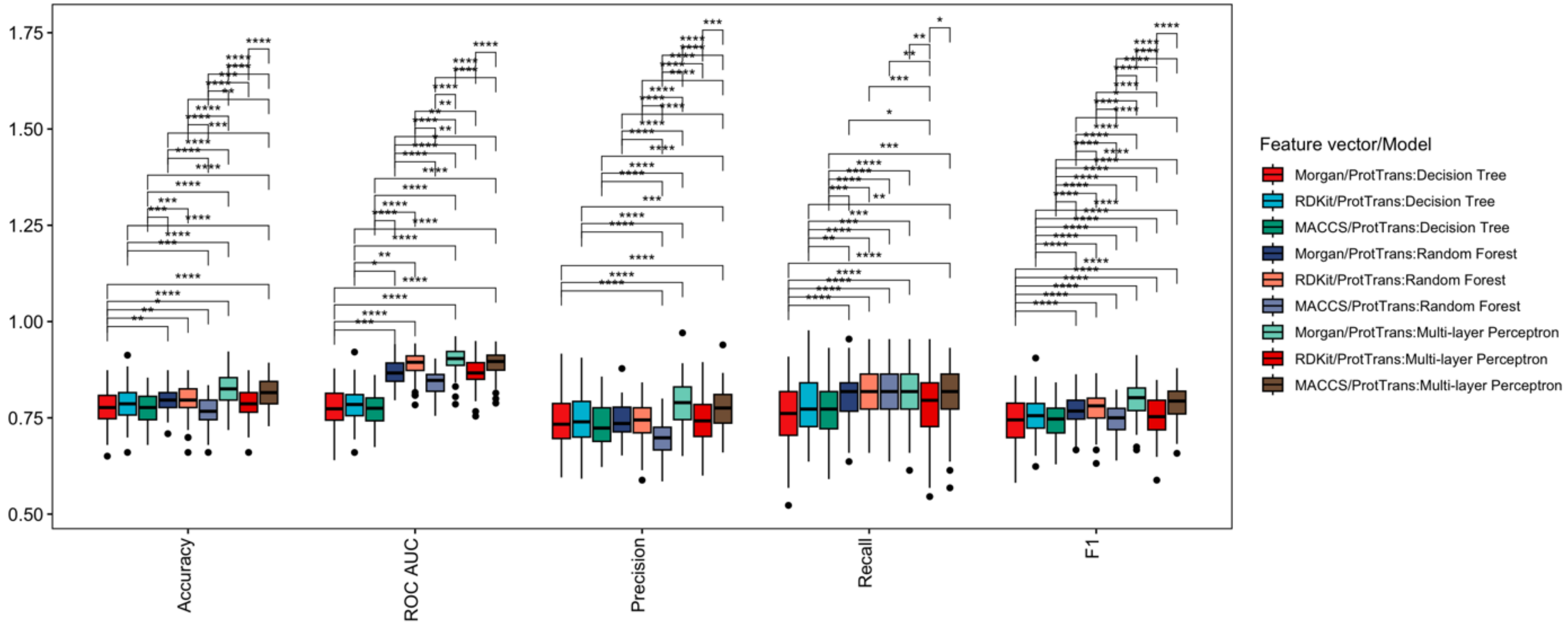

**Supplementary Figure 4**. Boxplots of ML performance metrics for the tested ML algorithms and feature vectors obtained on the cell-free substrate activity dataset. Values are averaged from 100 datasets with random splits. * - $p \le 0.05$, ** - $p \le 0.01$, *** - $p \le 0.001$, **** - $p \le 0.0001$, ns is not shown for visibility.

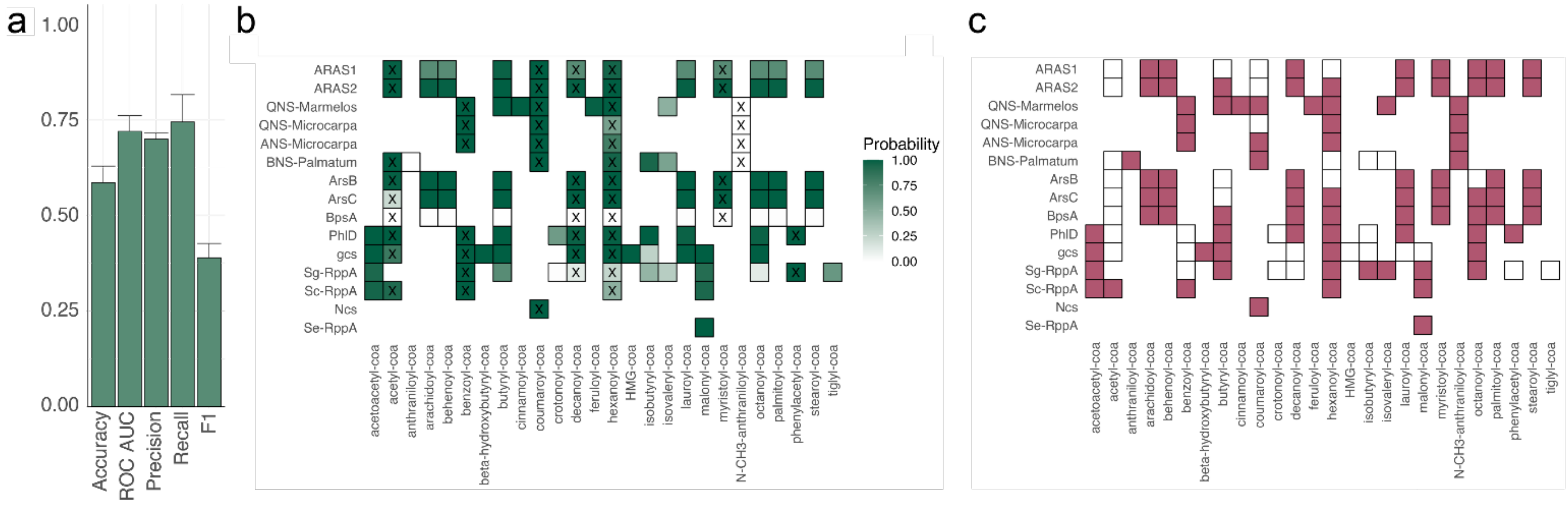

**Supplementary Figure 5.** Testing of phylogenetic bias of the predictive ML model. a) ML performance metrics of the Multi-Layer Perceptron algorithm with ProtTrans-X5/MACCS Keys feature vectors obtained for enzyme-substrate pairs from plant and bacteria from literature (see Supplementary Table 2). Predicted (b) and experimental (c) enzyme/substrate specificity of plant and bacterial T3PKSs. Substrates which were present in the training dataset are marked with X. Performance metrics and predicted enzyme/substrate specificity are averaged from 100 model runs with a random seed.

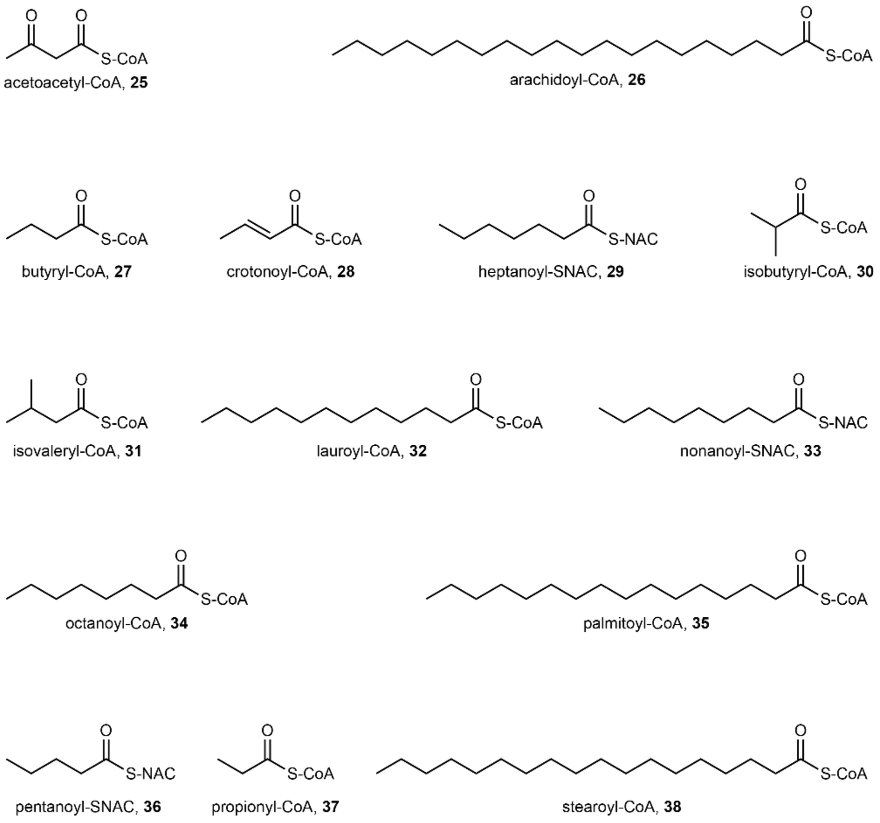

**Supplementary Figure 6.** Structures of compounds **25**-**38** previously reported to be accepted by fungal T3PKSs; NAC – N-acetylcysteamine, a CoA analogue.

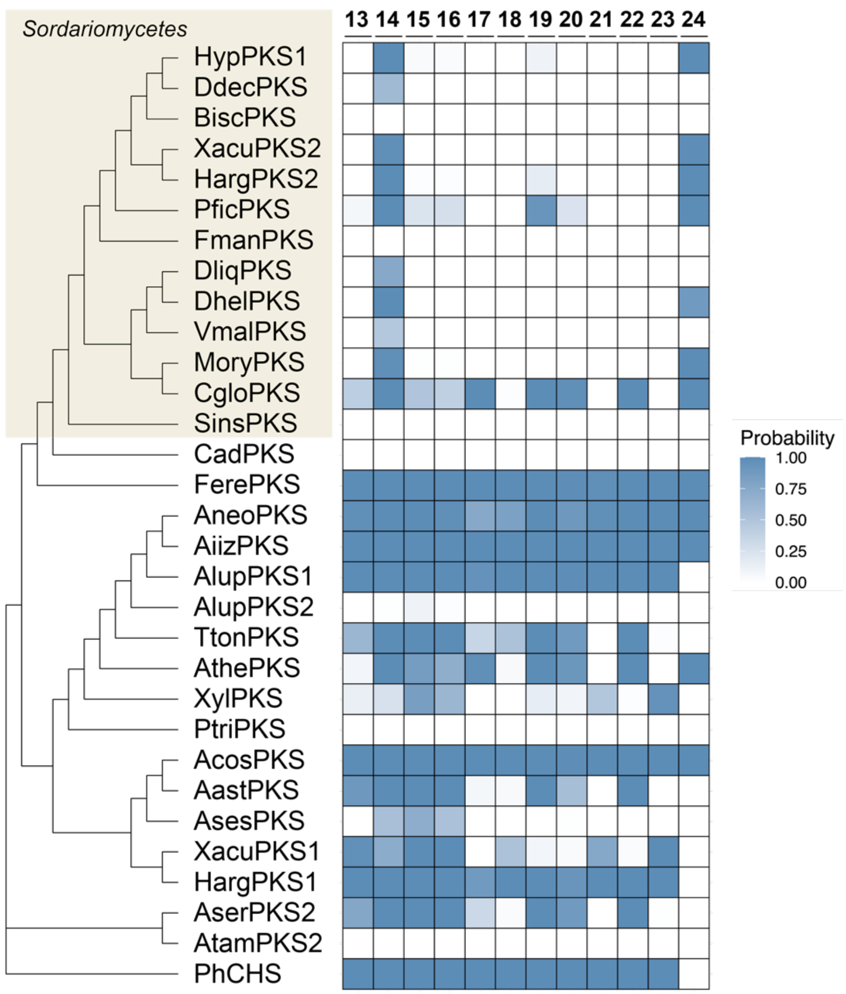

**Supplementary Figure 7.** Enzyme/substrate specificity prediction for 31 active T3PKSs with substrates **13-24** by the Multi-Layer Perceptron algorithm with the ProtTrans-X5/MACCS Keys feature vectors. Predictions are averaged from 100 model runs with a random seed. Seven T3PKSs that were not expressed at detectable levels were excluded from the analysis.

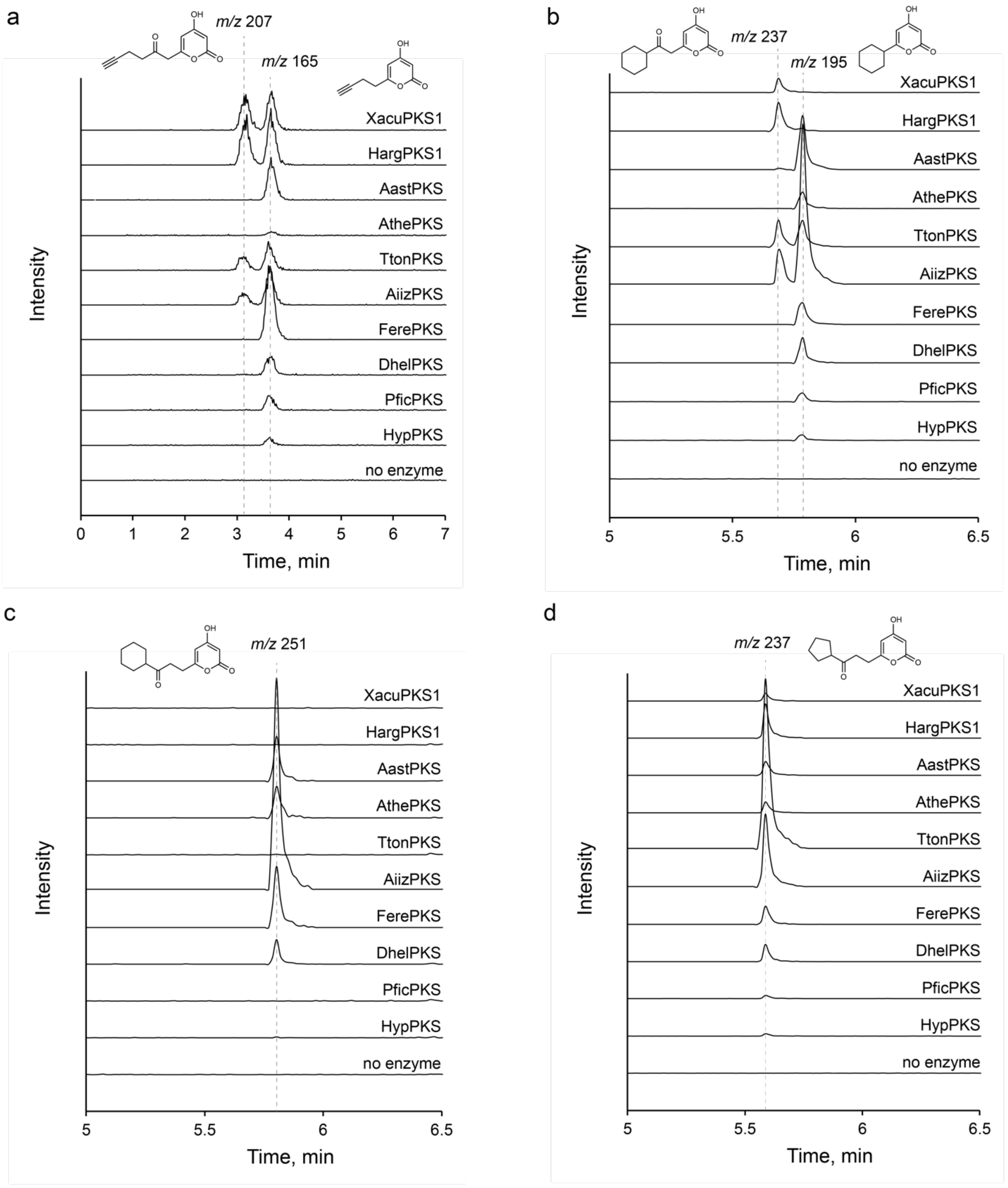
a
m/z 207
m/z 165
XacuPKS1
HargPKS1
AastPKS
AthePKS
TtonPKS
AiizPKS
FerePKS
DhelPKS
PficPKS
HypPKS
no enzyme
Intensity
0 1 2 3 4 5 6 7
Time, min
b
m/z 237
m/z 195
XacuPKS1
HargPKS1
AastPKS
AthePKS
TtonPKS
AiizPKS
FerePKS
DhelPKS
PficPKS
HypPKS
no enzyme
Intensity
5 5.5 6 6.5
Time, min
c
m/z 251
XacuPKS1
HargPKS1
AastPKS
AthePKS
TtonPKS
AiizPKS
FerePKS
DhelPKS
PficPKS
HypPKS
no enzyme
Intensity
5 5.5 6 6.5
Time, min
d
m/z 237
XacuPKS1
HargPKS1
AastPKS
AthePKS
TtonPKS
AiizPKS
FerePKS
DhelPKS
PficPKS
HypPKS
no enzyme
Intensity
5 5.5 6 6.5
Time, min

e

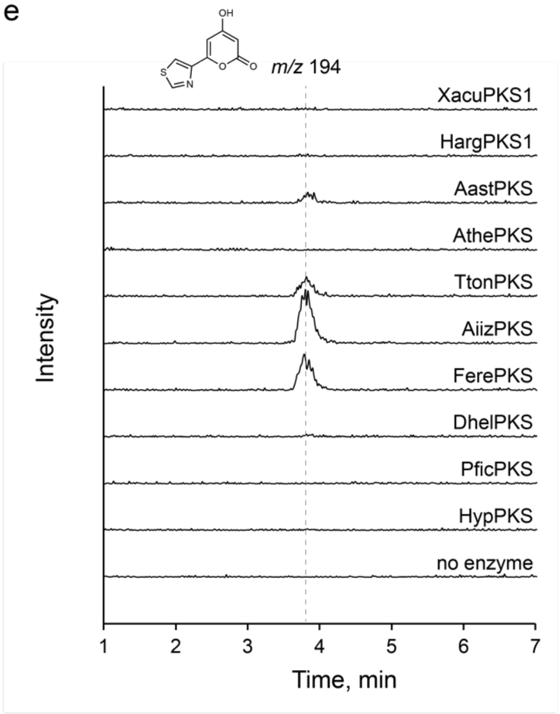
m/z 194
XacuPKS1
HargPKS1
AastPKS
AthePKS
TtonPKS
AiizPKS
FerePKS
DhelPKS
PficPKS
HypPKS
no enzyme
Intensity
1
2
3
4
5
6
7
Time, min

f

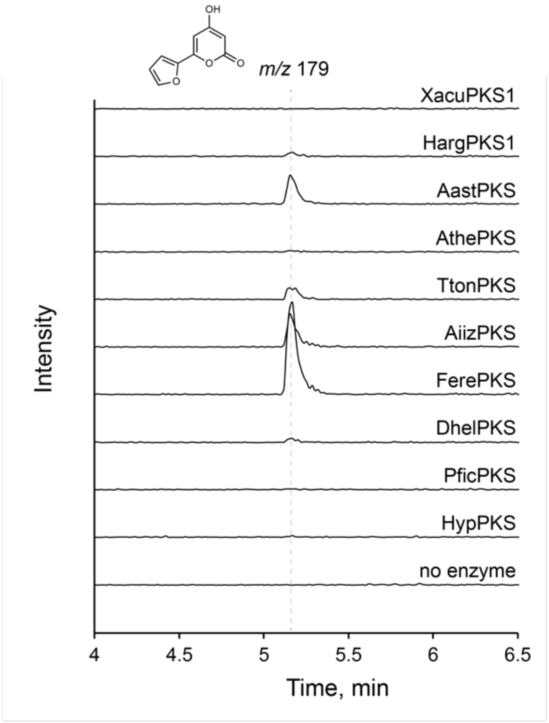
m/z 179
XacuPKS1
HargPKS1
AastPKS
AthePKS
TtonPKS
AiizPKS
FerePKS
DhelPKS
PficPKS
HypPKS
no enzyme
Intensity
4
4.5
5
5.5
6
6.5
Time, min

g

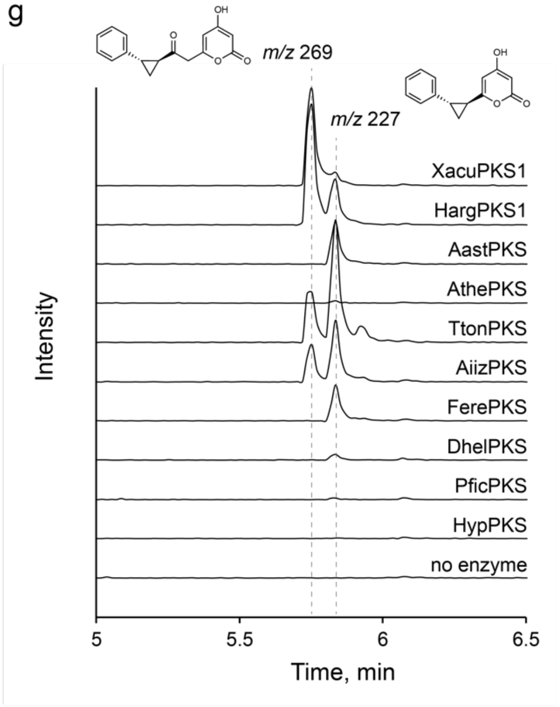
m/z 269
m/z 227
XacuPKS1
HargPKS1
AastPKS
AthePKS
TtonPKS
AiizPKS
FerePKS
DhelPKS
PficPKS
HypPKS
no enzyme
Intensity
5
5.5
6
6.5
Time, min

h

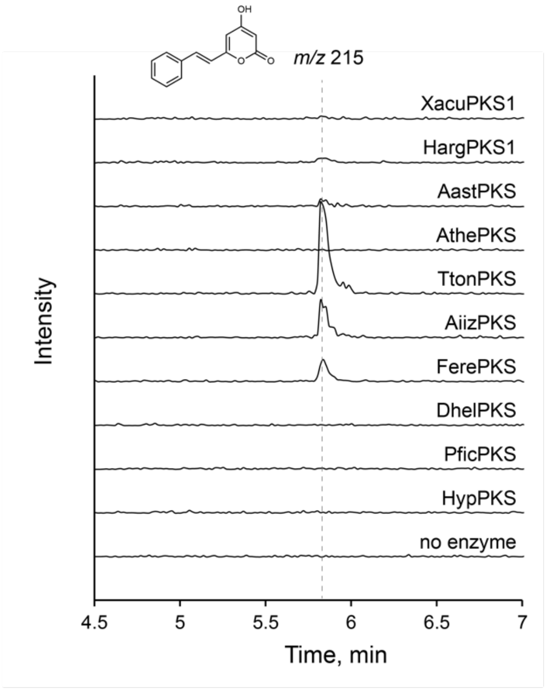
m/z 215
XacuPKS1
HargPKS1
AastPKS
AthePKS
TtonPKS
AiizPKS
FerePKS
DhelPKS
PficPKS
HypPKS
no enzyme
Intensity
4.5
5
5.5
6
6.5
7
Time, min

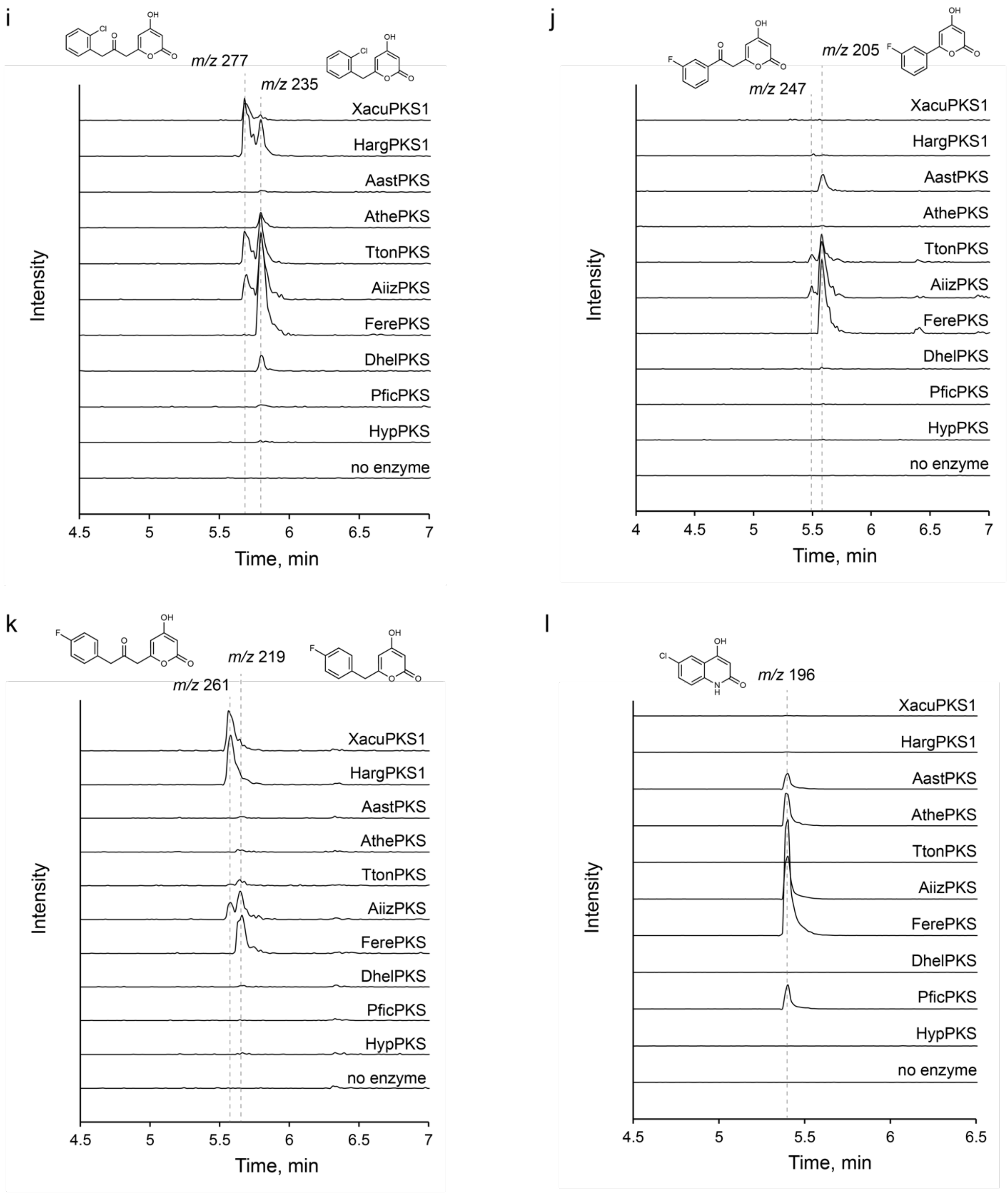

**Supplementary Figure 8.** Extracted ion chromatograms in low-resolution LC-MS of reaction products of the purified T3PKSs with substrates **13-24** compared to the "no enzyme" control: a) substrate **13**; b) substrate **14**; c) substrate **15**; d) substrate **16**; e) substrate **17**; f) substrate **18**; g) substrate **19**; h) substrate **20**; i) substrate **21**; j) substrate **22**; k) substrate **23**; l) substrate **24**.

The following mass-to-charge ratios (*m/z*) were detected in positive ion mode: putative triketide pyrone from **13**, detected *m/z* = 165 (theoretical *m/z* = 165.0546, calculated for $[C_9H_9O_3]^+$); putative tetraketide pyrone from **13**, detected *m/z* = 207 (theoretical *m/z* = 207.0652, calculated for $[C_{11}H_{11}O_4]^+$); putative triketide pyrone from **14**, detected *m/z* = 195 (theoretical *m/z* = 195.1016, calculated for $[C_{11}H_{15}O_3]^+$); putative tetraketide pyrone from **14**, detected *m/z* = 237 (theoretical *m/z* = 237.1121, calculated for $[C_{13}H_{17}O_4]^+$); putative triketide pyrone from **15**, detected *m/z* = 251 (theoretical *m/z* = 251.1278, calculated for $[C_{14}H_{19}O_4]^+$); putative triketide pyrone from **16**, detected *m/z* = 237 (theoretical *m/z* = 237.1121, calculated for $[C_{13}H_{17}O_4]^+$); putative triketide pyrone from **18**, detected *m/z* = 179 (theoretical *m/z* = 179.0339, calculated for $[C_9H_7O_4]^+$); putative triketide pyrone from **20**, detected *m/z* = 215 (theoretical *m/z* = 215.0703, calculated for $[C_{13}H_{11}O_3]^+$); putative diketide quinolone from **24**, detected *m/z* = 196 (theoretical *m/z* = 196.0160, calculated for $[C_9H_7ClNO_2]^+$).

The following mass-to-charge ratios (*m/z*) were detected in negative ion mode: putative triketide pyrone from **17**, detected *m/z* = 194 (theoretical *m/z* = 193.9917, calculated for $[C_8H_4NO_3S]^-$); putative triketide pyrone from **19**, detected *m/z* = 227 (theoretical *m/z* = 227.0714, calculated for $[C_{14}H_{11}O_3]^-$); putative tetraketide pyrone from **19**, detected *m/z* = 269 (theoretical *m/z* = 269.0819, calculated for $[C_{16}H_{13}O_4]^-$); putative triketide pyrone from **21**, detected *m/z* = 235 (theoretical *m/z* = 235.0167, calculated for $[C_{12}H_8ClO_3]^-$); putative tetraketide pyrone from **21**, detected *m/z* = 277 (theoretical *m/z* = 277.0273, calculated for $[C_{14}H_{10}ClO_4]^-$); putative triketide pyrone from **22**, detected *m/z* = 205 (theoretical *m/z* = 205.0306, calculated for $[C_{11}H_6FO_3]^-$); putative tetraketide pyrone from **22**, detected *m/z* = 247 (theoretical *m/z* = 247.0412, calculated for $[C_{13}H_8FO_4]^-$); putative triketide pyrone from **23**, detected *m/z* = 219 (theoretical *m/z* = 219.0463, calculated for $[C_{12}H_8FO_3]^-$); putative tetraketide pyrone from **23**, detected *m/z* = 261 (theoretical *m/z* = 261.0569, calculated for $[C_{14}H_{10}FO_4]^-$).

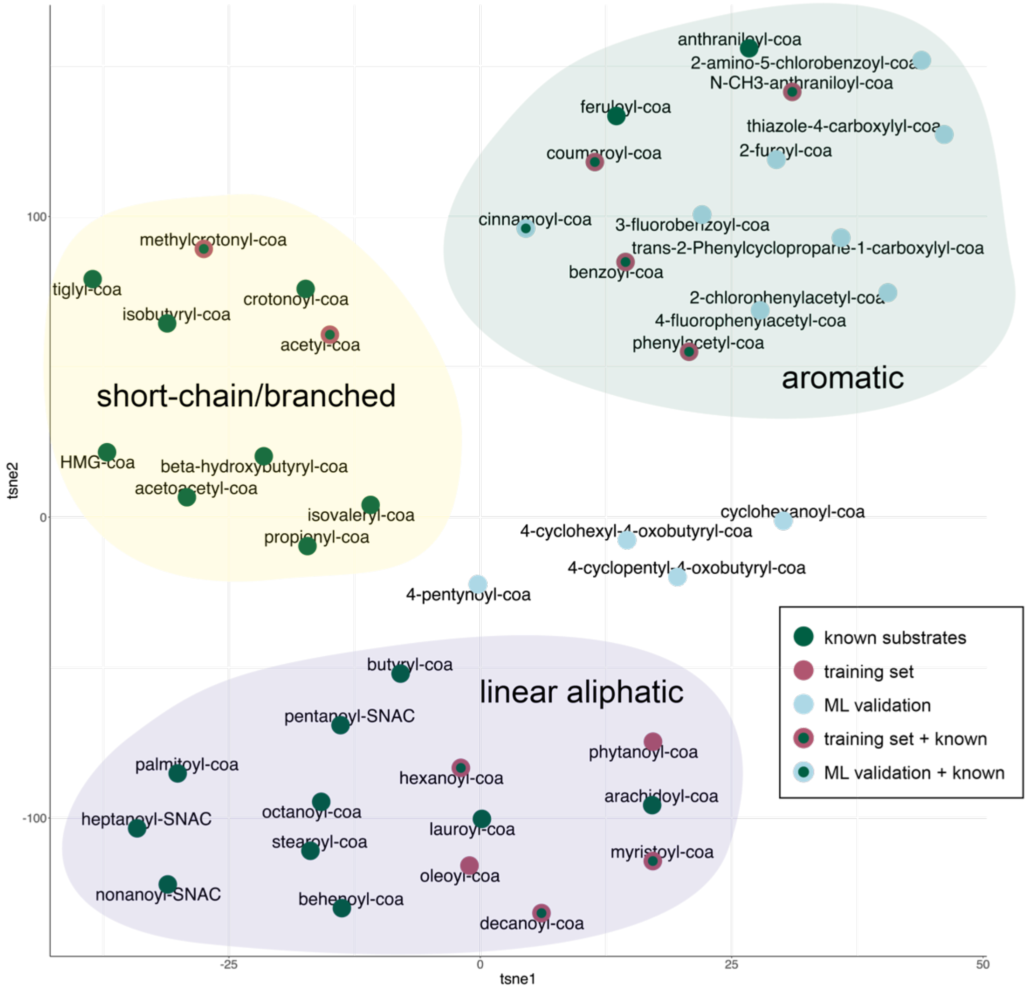

**Supplementary Figure 9.** Representation of the chemical space of T3PKS substrates obtained by t-SNE dimensionality reduction of a substrate similarity matrix. The matrix was obtained by calculating pairwise similarity of the substrates' MACCS Keys using the RDKit *DataStructs.DiceSimilarity* function. Dimensionality reduction was performed using scikit-learn *manifold.TSNE* with perplexity 10. Clusters containing linear, short chain/branched and aromatic CoA thioesters are highlighted. Nodes are coloured based on the source of the substrate.

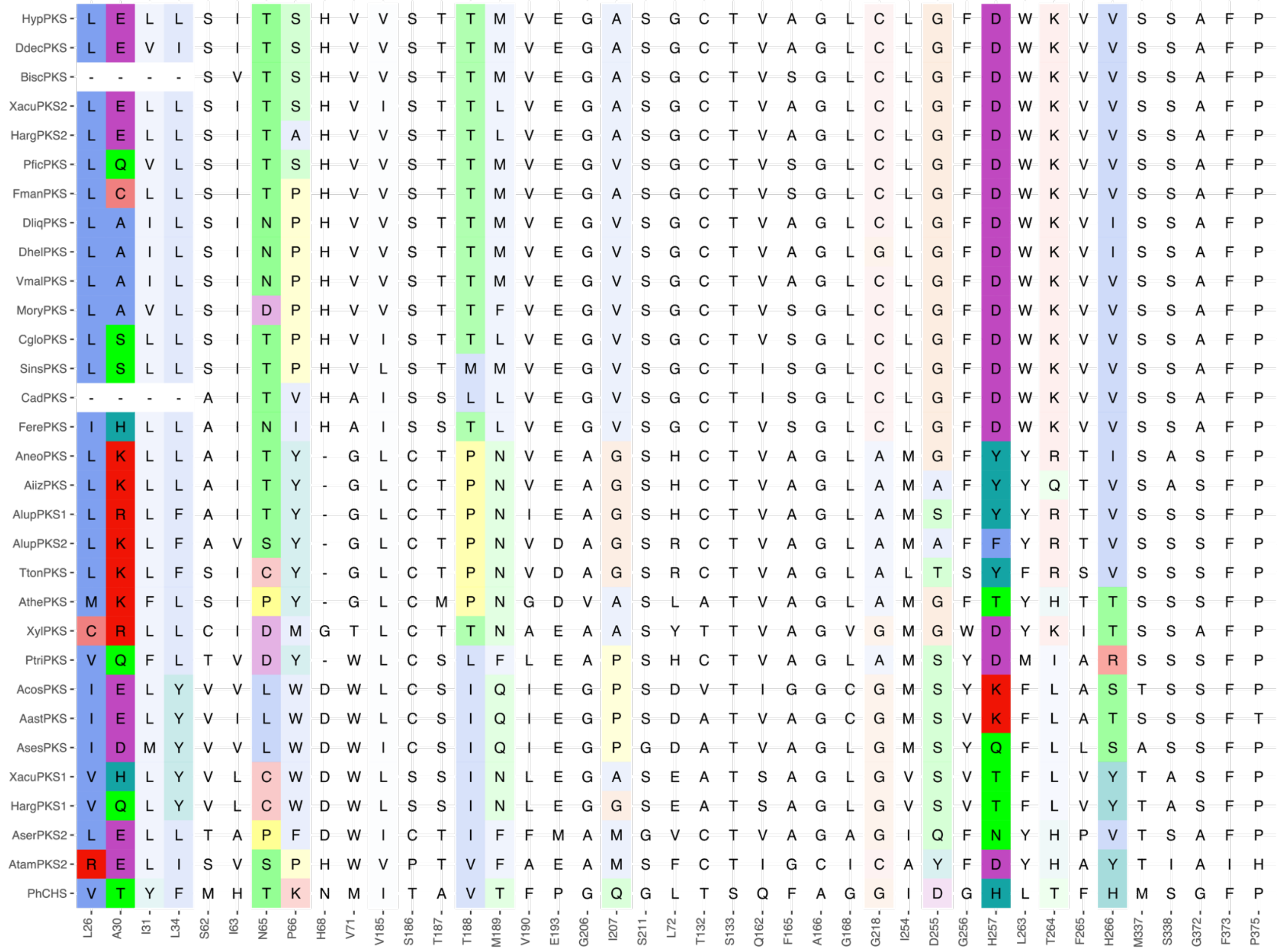

**Supplementary Figure 10.** Multiple sequence alignment of the amino acids lining active site and substrate-binding tunnel in the 31 expressed T3PKSs. The residues are highlighted using Clustal X Default Colouring; colour intensity is scaled according to residue importances as determined by the descriptive ML model.

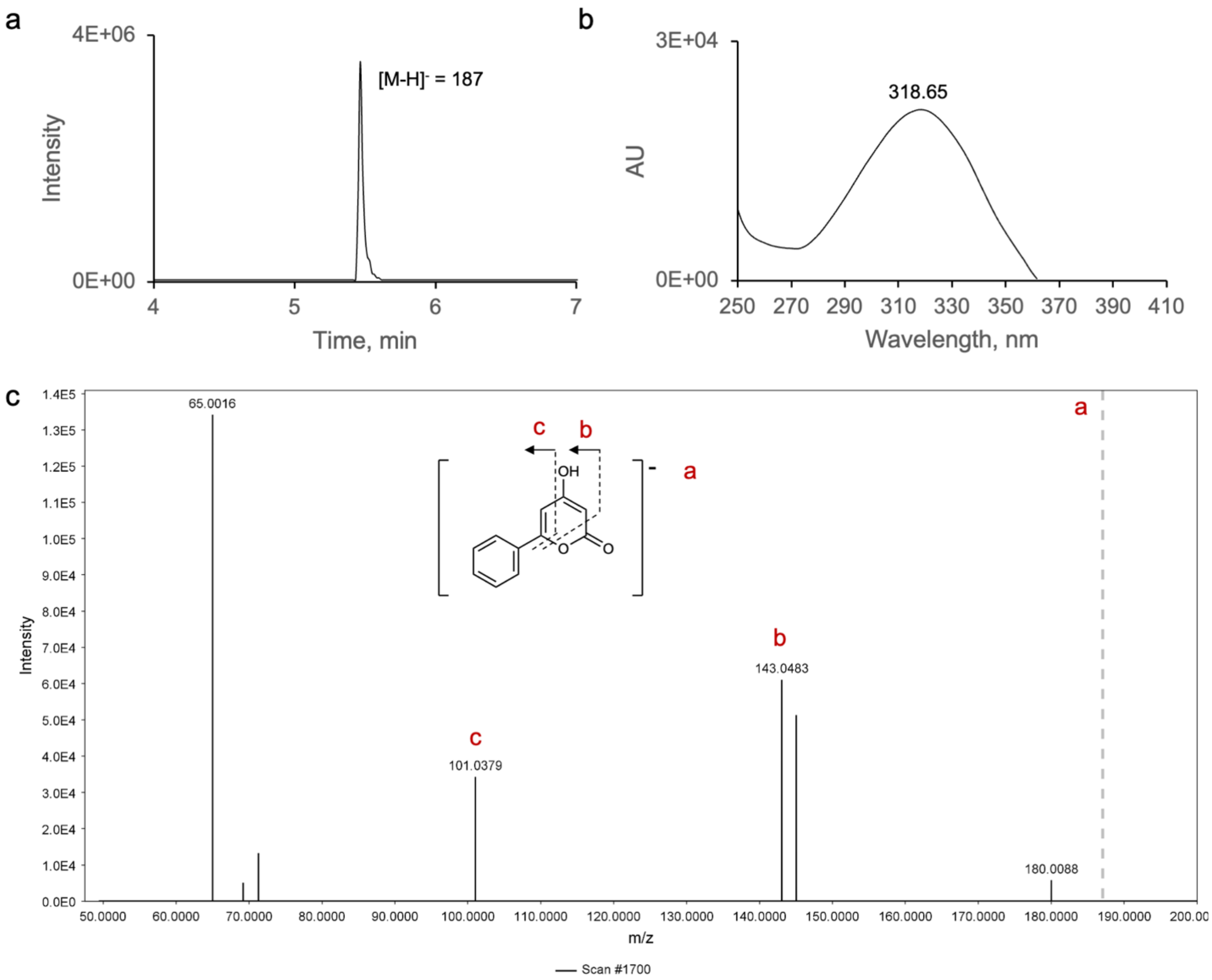

**Supplementary Figure 11.** Spectral data analysis of product **1a**. a) Low-resolution LC-MS analysis of the EtOAc extract of the enzymatic reaction of AiizPKS with **1**; extracted ion chromatogram of the predicted *m/z* of 187 (negative mode). Y-axis shows relative ion intensity. b) Corresponding UV absorption spectrum. c) ESI-HR-MS/MS (negative mode) with ions matching expected fragments of **1a**; observed *m/z* = 187.0391 (theoretical *m/z* = 187.0401, calculated for $[C_{11}H_7O_3]^-$). The precursor ion is indicated with a dashed grey line.

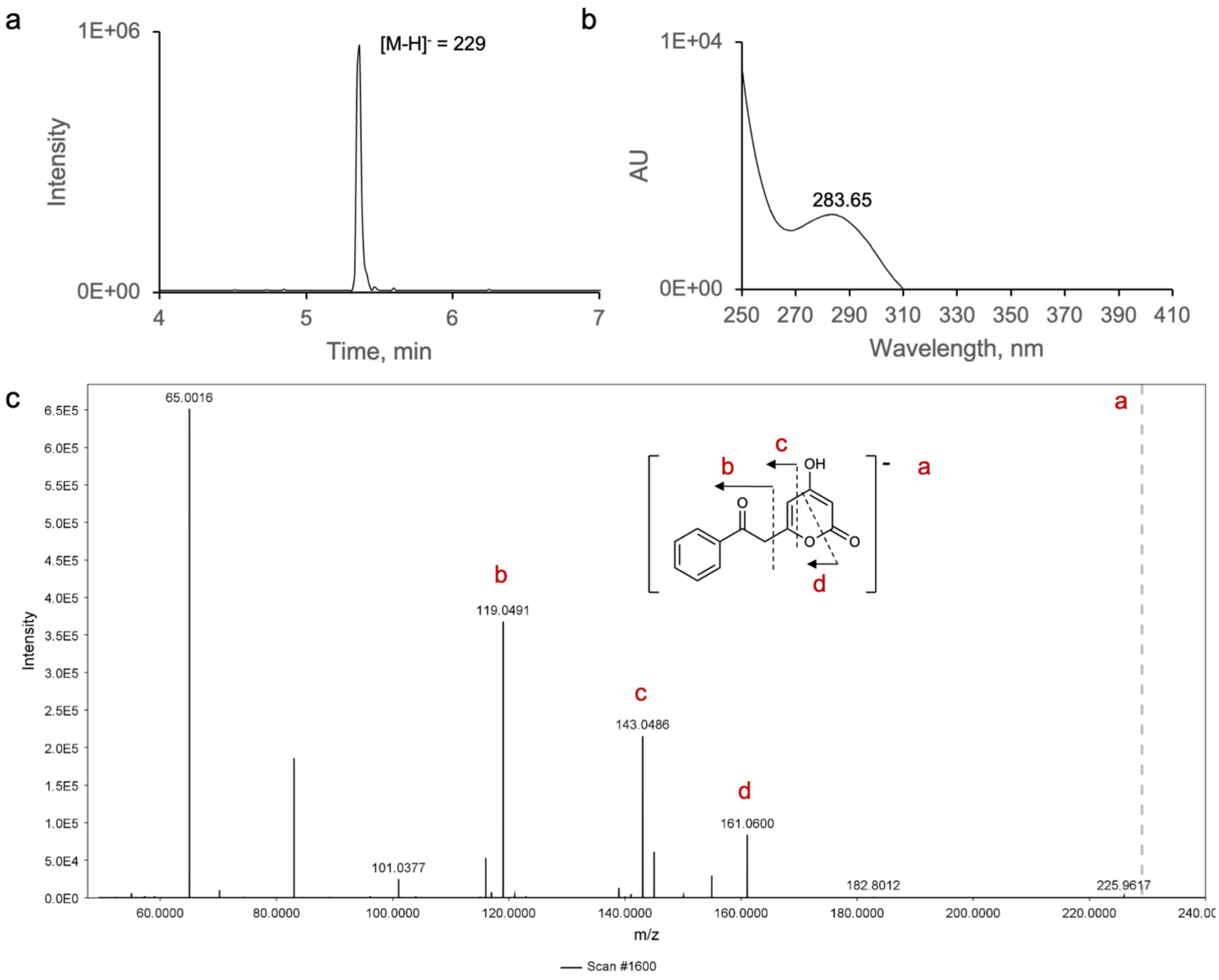

**Supplementary Figure 12.** Spectral data analysis of product **1b**. a) Low-resolution LC-MS analysis of the EtOAc extract of the enzymatic reaction of AiizPKS with **1**; extracted ion chromatogram of the predicted *m/z* of 229 (negative mode). Y-axis shows relative ion intensity. b) Corresponding UV absorption spectrum. c) ESI-HR-MS/MS (negative mode) with ions matching expected fragments of **1b**; observed *m/z* = 229.0503 (theoretical *m/z* = 229.0506, calculated for $[C_{13}H_9O_4]^-$). The precursor ion is indicated with a dashed grey line.

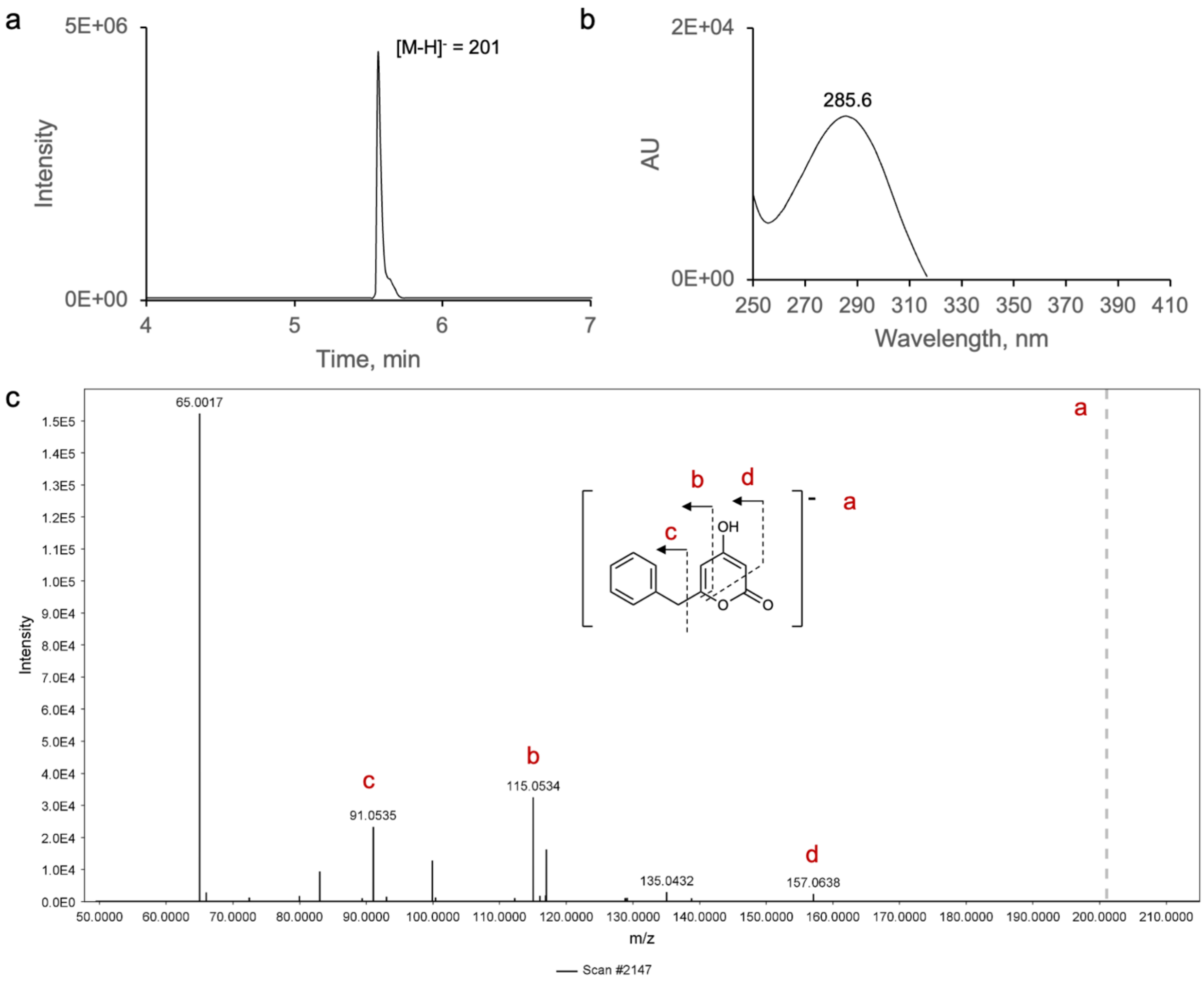

**Supplementary Figure 13.** Spectral data analysis of product **2a**. a) Low-resolution LC-MS analysis of the EtOAc extract of the enzymatic reaction of AlupPKS1 with **2**; extracted ion chromatogram of the predicted *m/z* of 201 (negative mode). Y-axis shows relative ion intensity. b) Corresponding UV absorption spectrum. c) ESI-HR-MS/MS (negative mode) with ions matching expected fragments of **2a**; observed *m/z* = 201.0547 (theoretical *m/z* = 201.0557, calculated for $[C_{12}H_9O_3]^-$). The precursor ion is indicated with a dashed grey line.

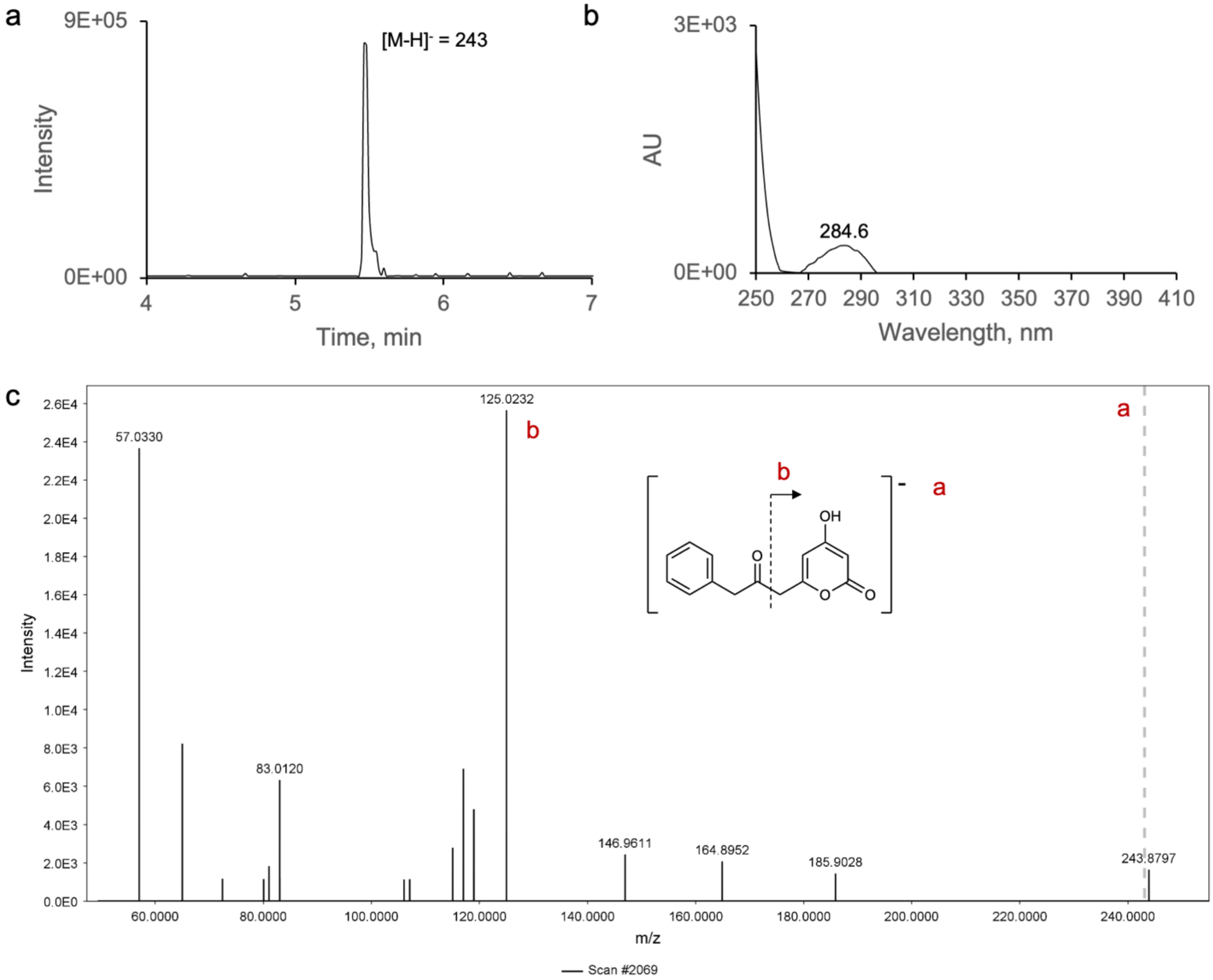

**Supplementary Figure 14.** Spectral data analysis of product **2b**. a) Low-resolution LC-MS analysis of the EtOAc extract of the enzymatic reaction of AlupPKS1 with **2**; extracted ion chromatogram of the predicted *m/z* of 243 (negative mode). Y-axis shows relative ion intensity. b) Corresponding UV absorption spectrum. c) ESI-HR-MS/MS (negative mode) with ions matching expected fragments of **2b**; observed *m/z* = 243.0657 (theoretical *m/z* = 243.0663, calculated for $[C_{14}H_{11}O_4]^-$). The precursor ion is indicated with a dashed grey line.

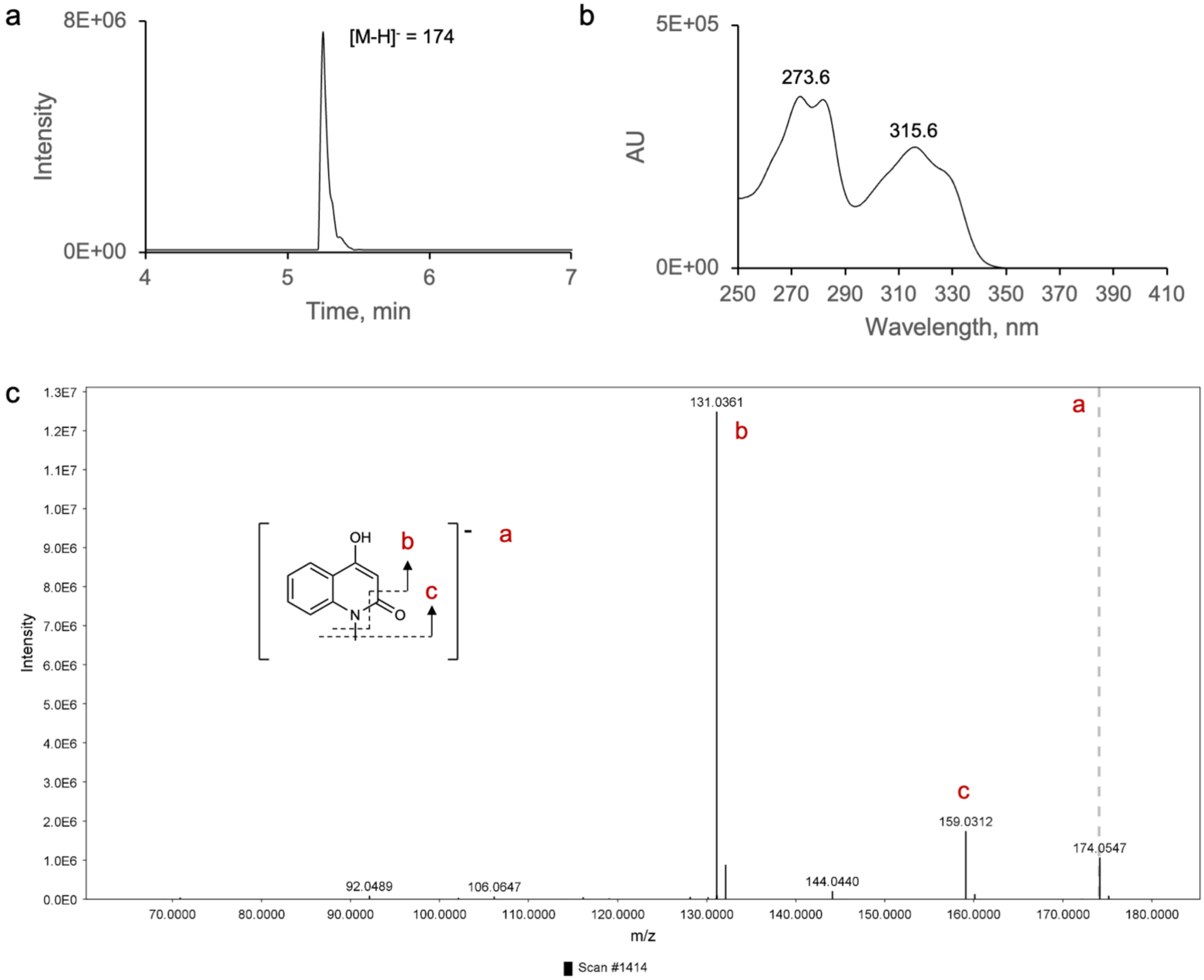

**Supplementary Figure 15.** Spectral data analysis of product **3a**. a) Low-resolution LC-MS analysis of the EtOAc extract of the enzymatic reaction of AthePKS with **3**; extracted ion chromatogram of the predicted *m/zm/z* of 174 (negative mode). Y-axis shows relative ion intensity. b) Corresponding UV absorption spectrum. c) ESI-HR-MS/MS (negative mode) with ions matching expected fragments of **3a**; observed *m/z* = 174.0546 (theoretical *m/z* = 174.0561, calculated for $[C_{10}H_8NO_2]^-$). The precursor ion is indicated with a dashed grey line.

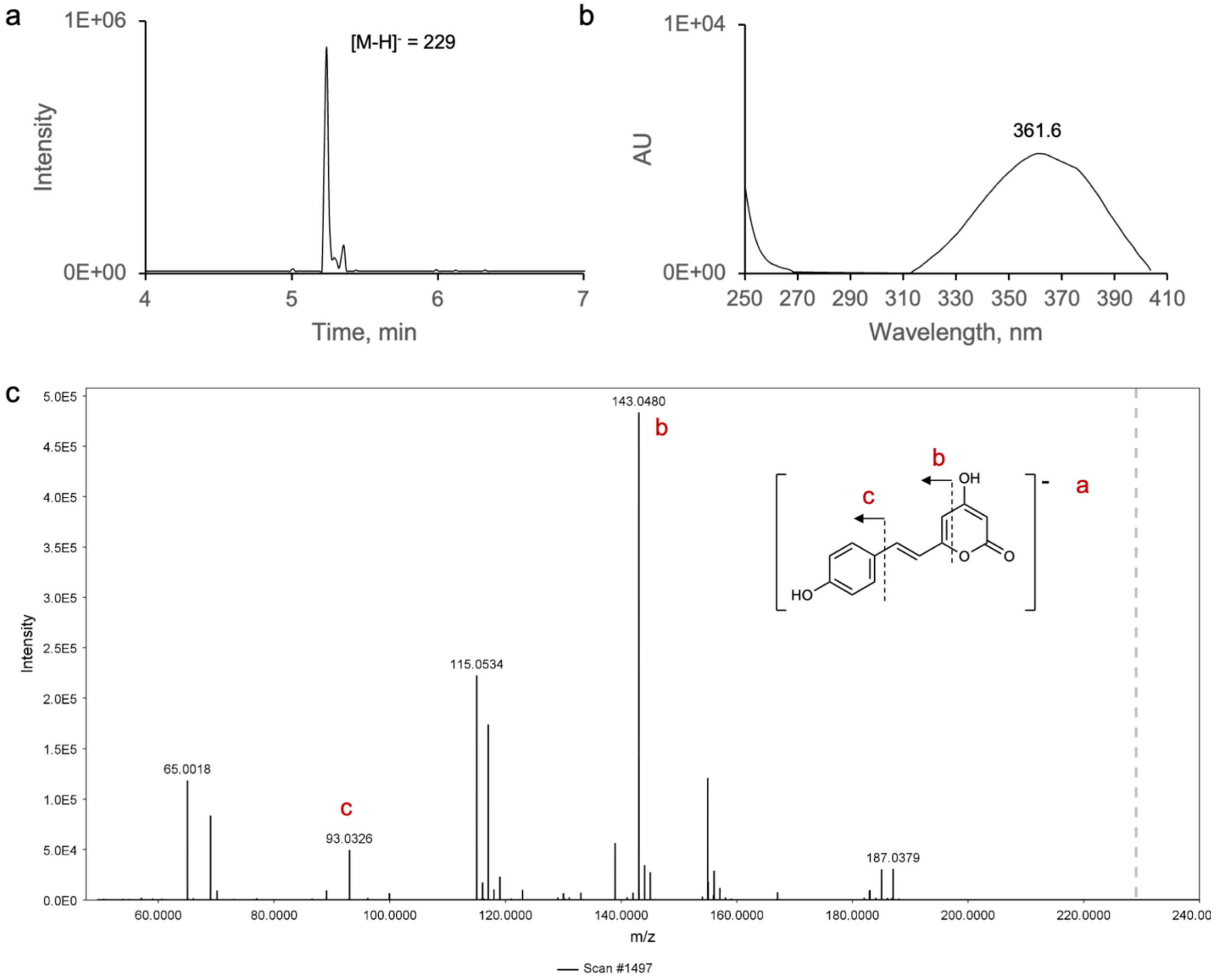

**Supplementary Figure 16.** Spectral data analysis of product **4a**. a) Low-resolution LC-MS analysis of the EtOAc extract of the enzymatic reaction of 4 with HargPKS1; extracted ion chromatogram of the predicted *m/z* of 229 (negative mode). Y-axis shows relative ion intensity. b) Corresponding UV absorption spectrum. c) ESI-HR-MS/MS (negative mode) with ions matching expected fragments of **4a**; observed *m/z* = 229.0500 (theoretical *m/z* = 229.0506, calculated for $[C_{13}H_9O_4]^-$). The precursor ion is indicated with a dashed grey line.

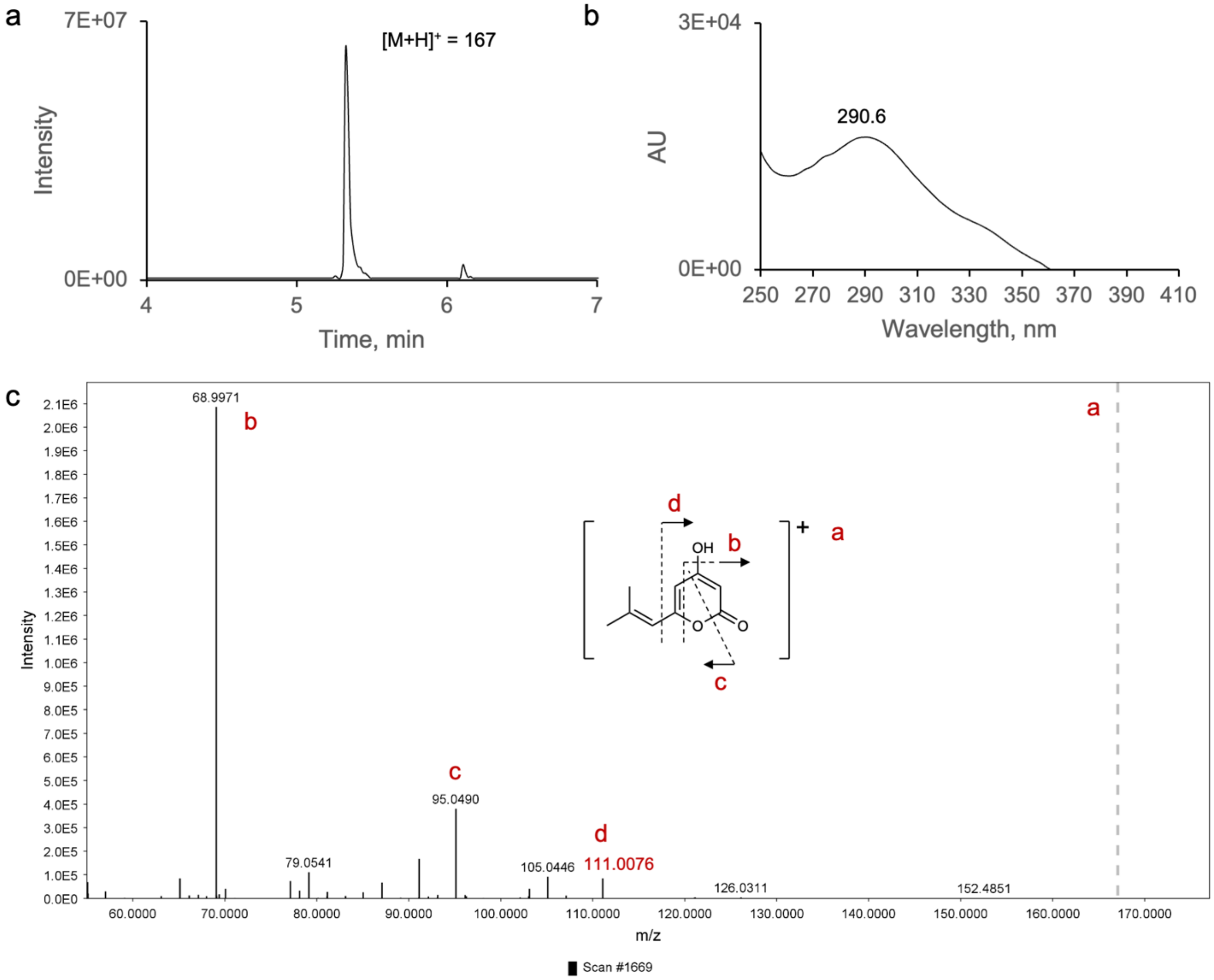

**Supplementary Figure 17.** Spectral data analysis of product **5a**. a) Low-resolution LC-MS analysis of the EtOAc extract of the enzymatic reaction of CgloPKS with **5**; extracted ion chromatogram of the predicted *m/z* of 167 (positive mode). Y-axis shows relative ion intensity. b) Corresponding UV absorption spectrum. c) ESI-HR-MS/MS (positive mode) with ions matching expected fragments of **5a**; observed *m/z* = 167.0703 (theoretical *m/z* = 167.0703, calculated for $[C_9H_{11}O_3]^+$). The precursor ion is indicated with a dashed grey line.

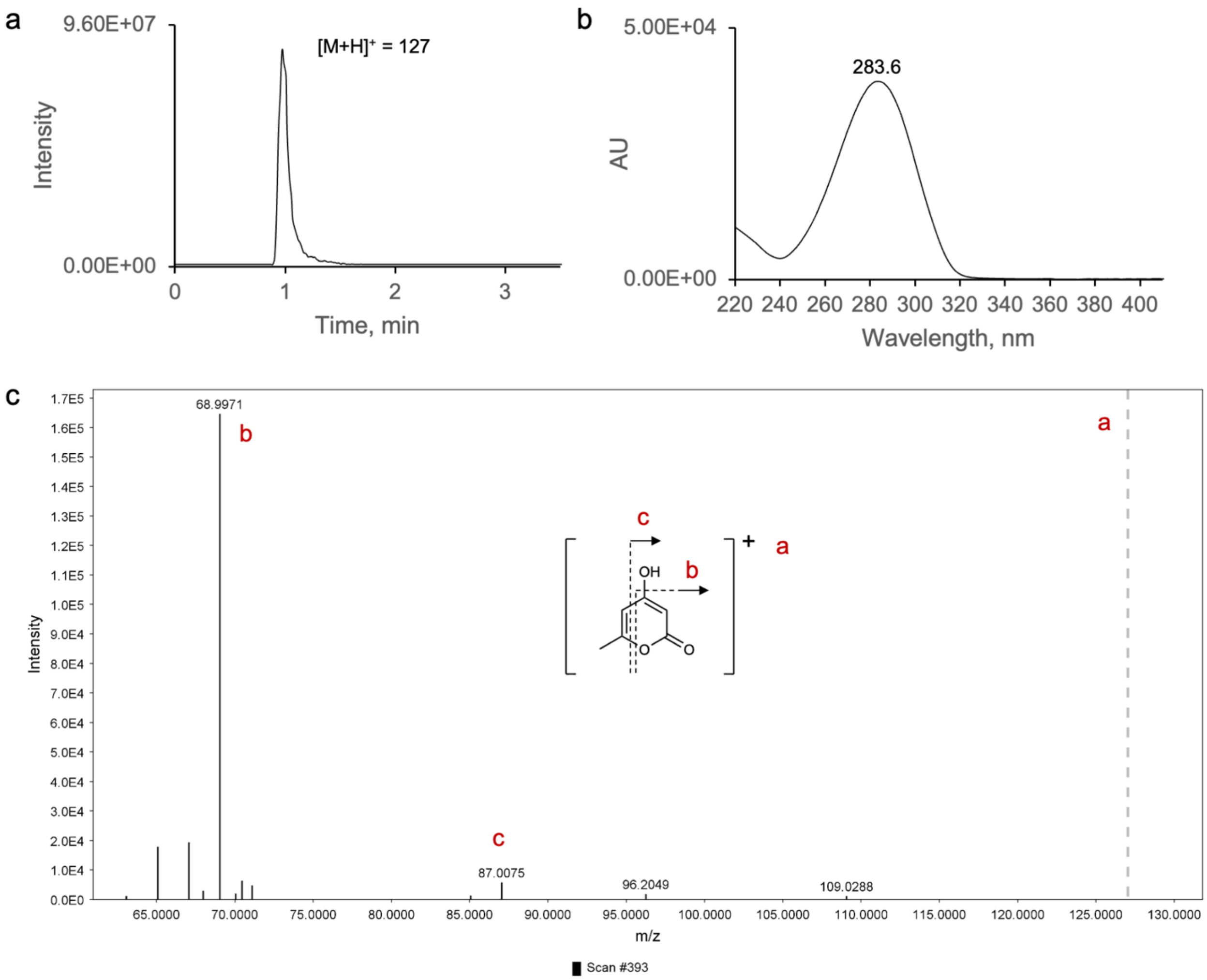

**Supplementary Figure 18.** Spectral data analysis of product **6a**. a) Low-resolution LC-MS analysis of the EtOAc extract of the enzymatic reaction of FerePKS with **6**; extracted ion chromatogram of the predicted *m/z* of 127 (positive mode). Y-axis shows relative ion intensity. b) Corresponding UV absorption spectrum. c) ESI-HR-MS/MS (positive mode) with ions matching expected fragments of **6a**; observed *m/z* = 127.0388 (theoretical *m/z* = 127.0390, calculated for $[C_6H_7O_3]^+$). The precursor ion is indicated with a dashed grey line.

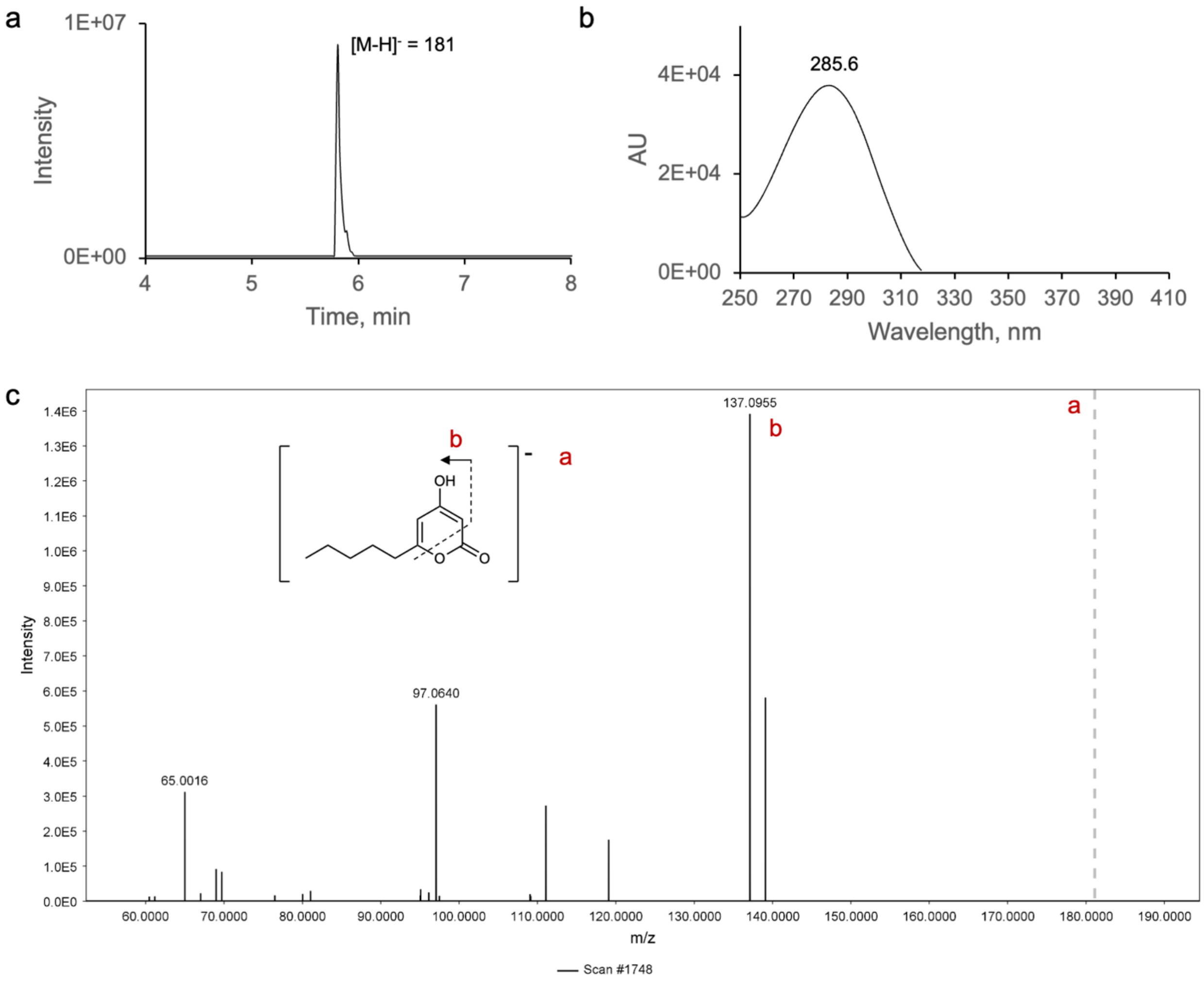

**Supplementary Figure 19.** Spectral data analysis of product **7a**. a) Low-resolution LC-MS analysis of the EtOAc extract of the enzymatic reaction of AserPKS2 with **7**; extracted ion chromatogram of the predicted *m/z* of 181 (negative mode). Y-axis shows relative ion intensity. b) Corresponding UV absorption spectrum. c) ESI-HR-MS/MS (negative mode) with ions matching expected fragments of **7a**; observed *m/z* = 181.0861 (theoretical *m/z* = 181.0870, calculated for $[C_{10}H_{13}O_3]^-$). The precursor ion is indicated with a dashed grey line.

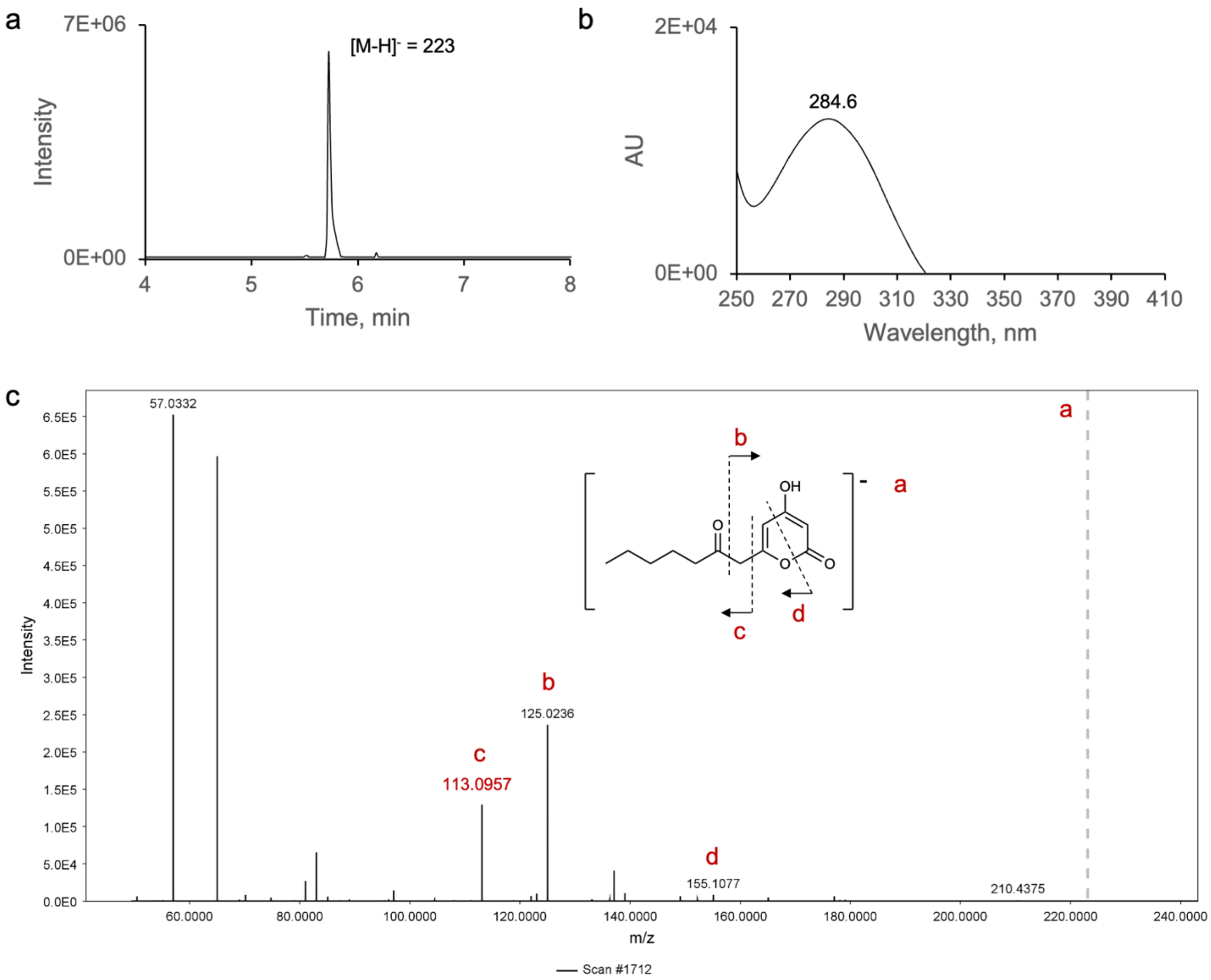

**Supplementary Figure 20.** Spectral data analysis of product **7b**. a) Low-resolution LC-MS analysis of the EtOAc extract of the enzymatic reaction of AserPKS2 with **7**; extracted ion chromatogram of the predicted *m/z* of 223 (negative mode). Y-axis shows relative ion intensity. b) Corresponding UV absorption spectrum. c) ESI-HR-MS/MS (xxx mode) with ions matching expected fragments of **7b**; observed *m/z* = 223.0973 (theoretical *m/z* = 223.0976, calculated for $[C_{12}H_{15}O_4]^-$). The precursor ion is indicated with a dashed grey line.

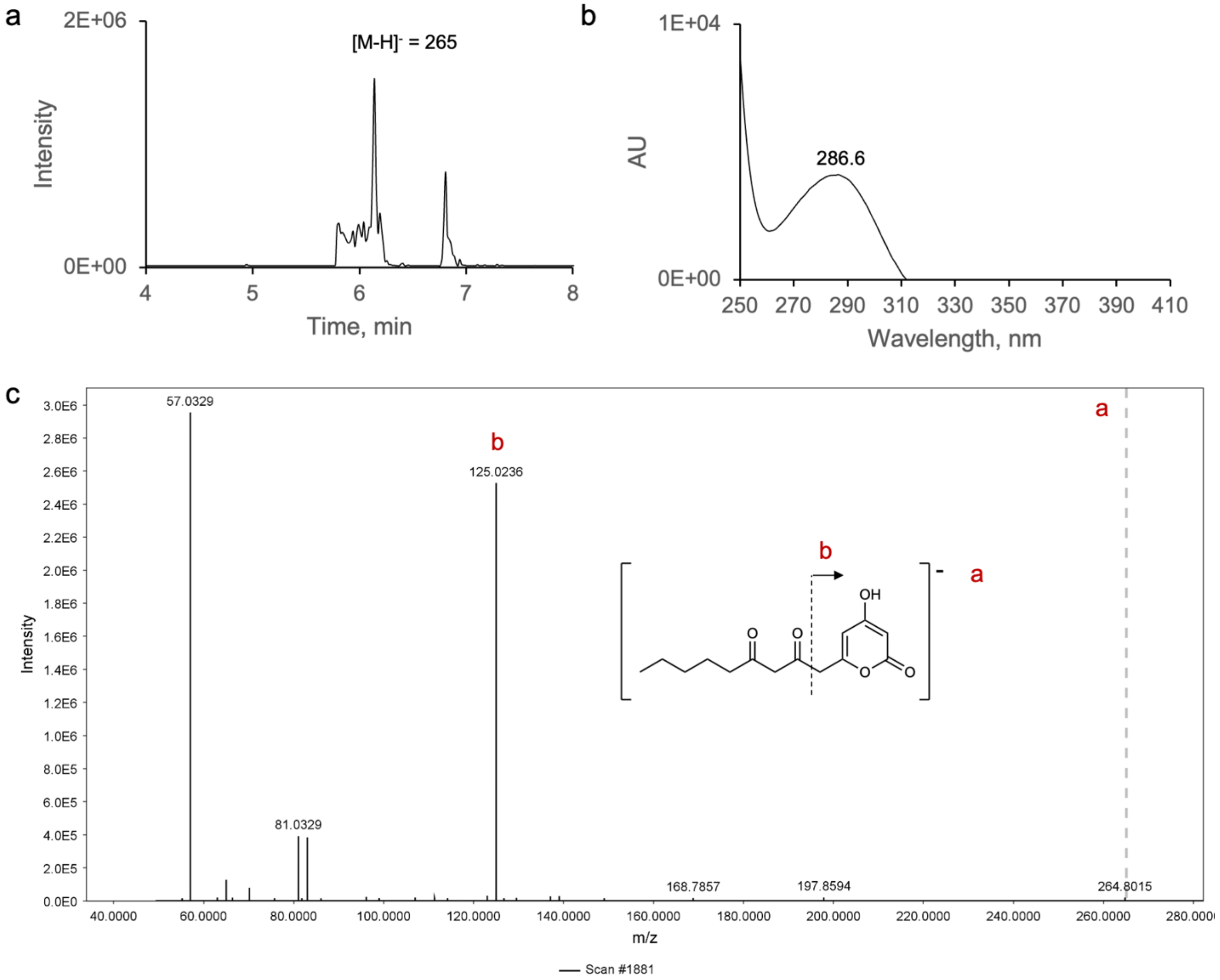

**Supplementary Figure 21.** Spectral data analysis of product **7c**. a) Low-resolution LC-MS analysis of the EtOAc extract of the enzymatic reaction of HargPKS1 with **7**; extracted ion chromatogram of the predicted *m/z* of 265 (negative mode). Y-axis shows relative ion intensity. b) Corresponding UV absorption spectrum. c) ESI-HR-MS/MS (negative mode) with ions matching expected fragments of **7c**; observed *m/z* = 265.1083 (theoretical *m/z* = 265.1081, calculated for $[C_{14}H_{17}O_5]^-$). The precursor ion is indicated with a dashed grey line.

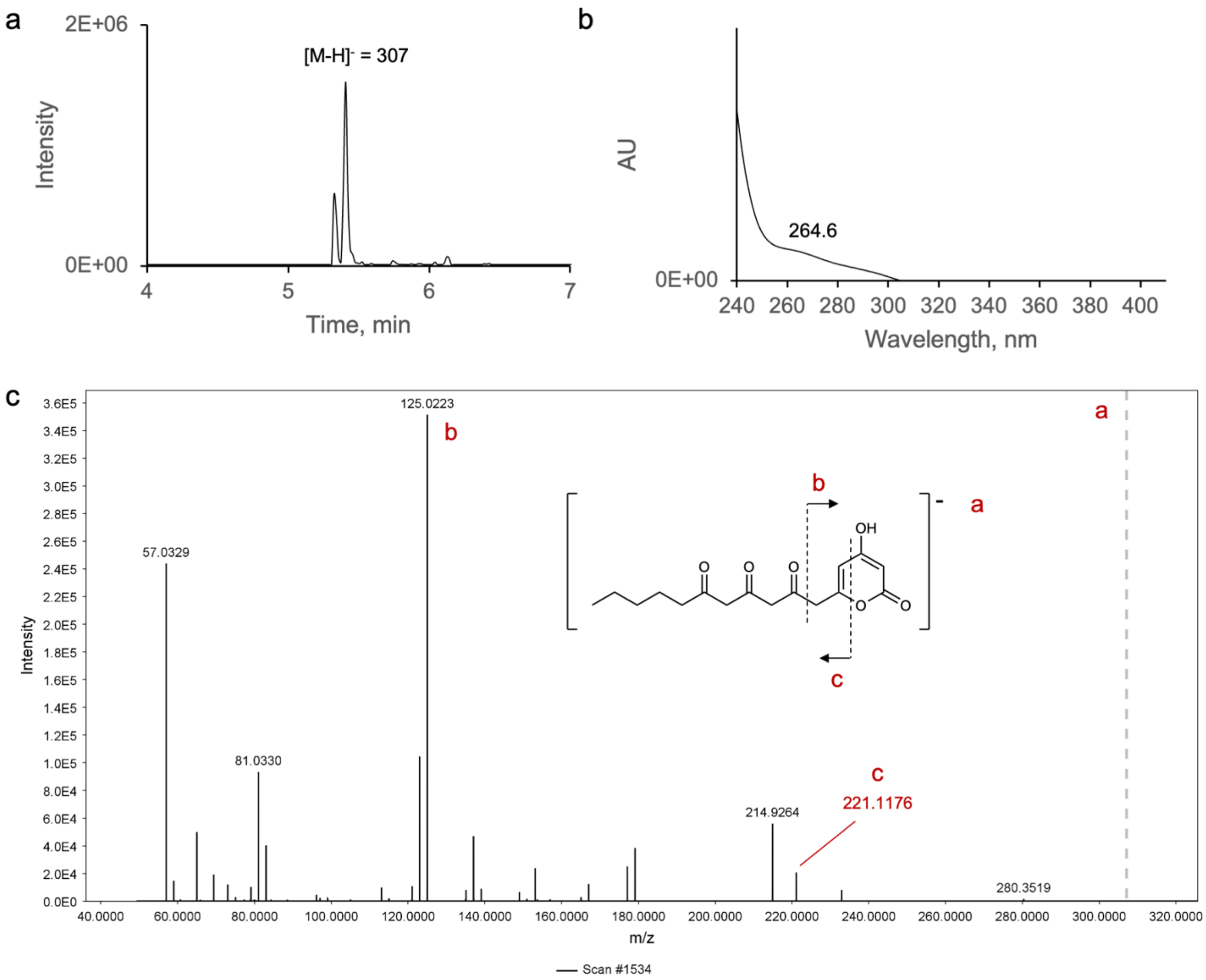

**Supplementary Figure 22.** Spectral data analysis of product **7d**. a) Low-resolution LC-MS analysis of the EtOAc extract of the enzymatic reaction of HargPKS1 with **7**; extracted ion chromatogram of the predicted *m/z* of 307 (negative mode). Y-axis shows relative ion intensity. b) Corresponding UV absorption spectrum. c) ESI-HR-MS/MS (negative mode) with ions matching expected fragments of **7d**; observed *m/z* = 307.1190 (theoretical *m/z* = 307.1187, calculated for $[C_{16}H_{19}O_6]^-$). The precursor ion is indicated with a dashed grey line.

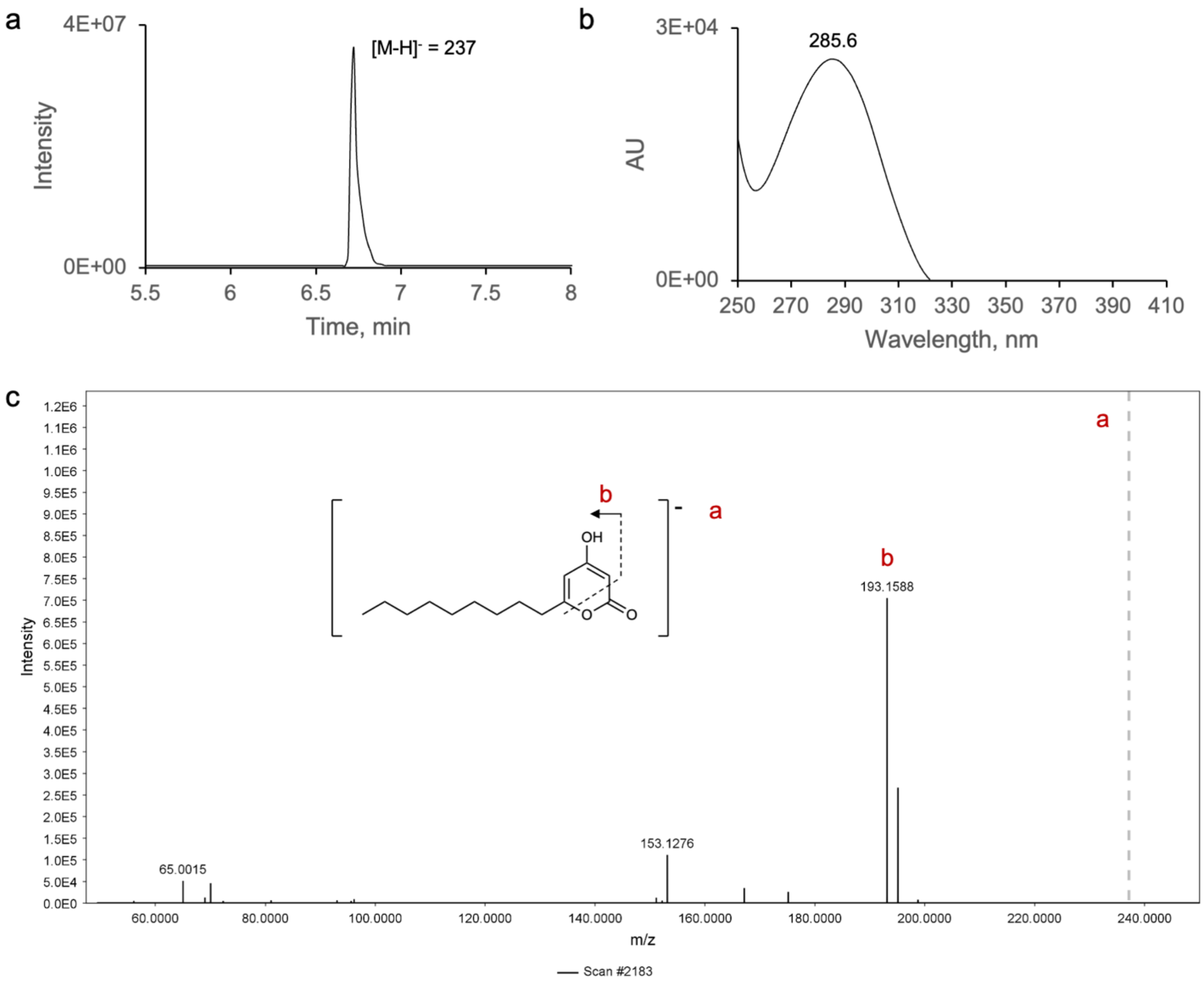

**Supplementary Figure 23.** Spectral data analysis of product **8a**. a) Low-resolution LC-MS analysis of the EtOAc extract of the enzymatic reaction of HargPKS1 with **8**; extracted ion chromatogram of the predicted *m/z* of 237 (negative mode). Y-axis shows relative ion intensity. b) Corresponding UV absorption spectrum. c) ESI-HR-MS/MS (negative mode) with ions matching expected fragments of **8a**; observed *m/z* = 237.1495 (theoretical *m/z* = 237.1496, calculated for $[C_{14}H_{21}O_3]^-$). The precursor ion is indicated with a dashed grey line.

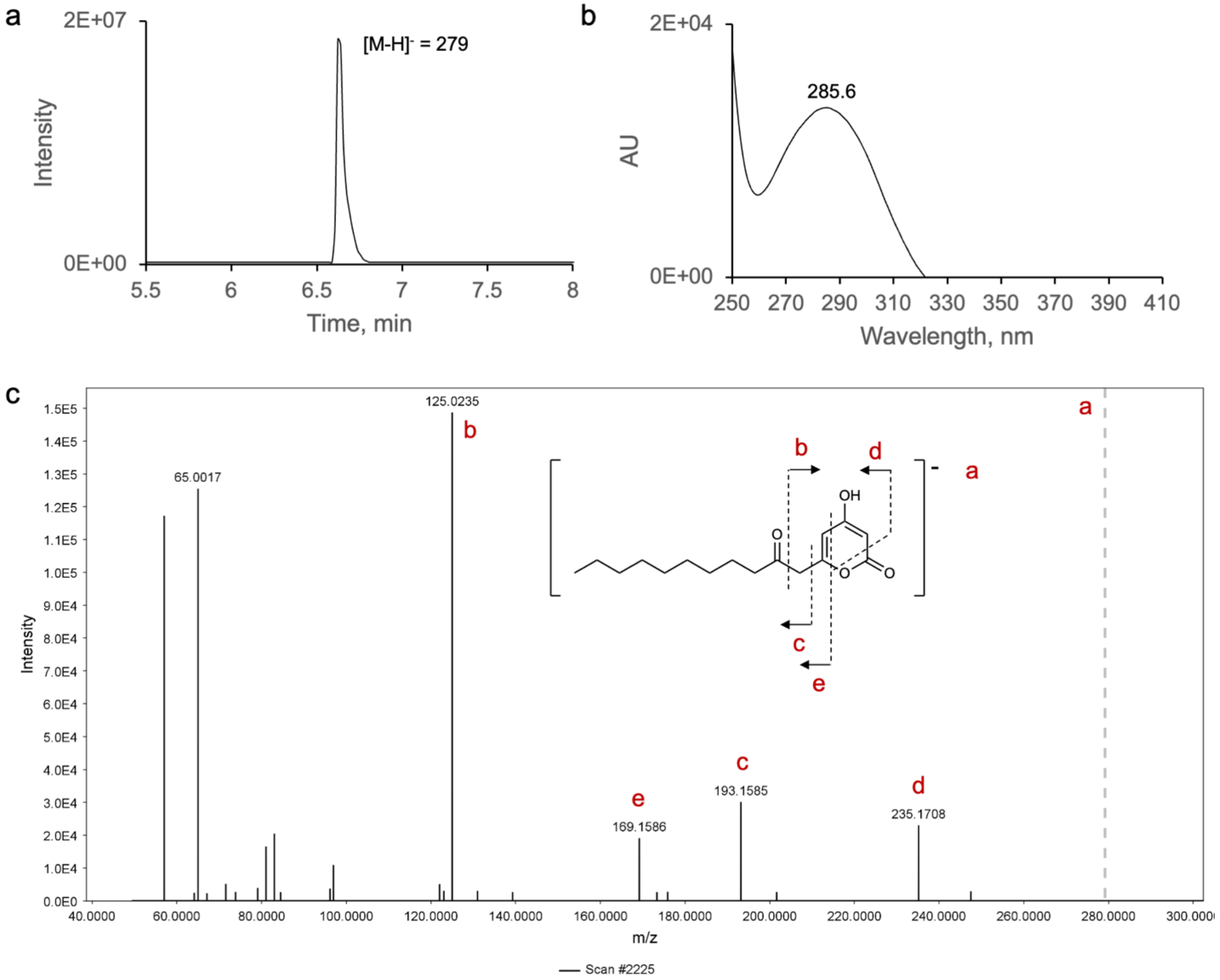

**Supplementary Figure 24.** Spectral data analysis of product **8b**. a) Low-resolution LC-MS analysis of the EtOAc extract of the enzymatic reaction of DhelPKS with **8**; extracted ion chromatogram of the predicted *m/z* of 279 (negative mode). Y-axis shows relative ion intensity. b) Corresponding UV absorption spectrum. c) ESI-HR-MS/MS (negative mode) with ions matching expected fragments of **8b**; observed *m/z* = 279.1603 (theoretical *m/z* = 279.1602, calculated for $[C_{16}H_{23}O_4]^-$). The precursor ion is indicated with a dashed grey line.

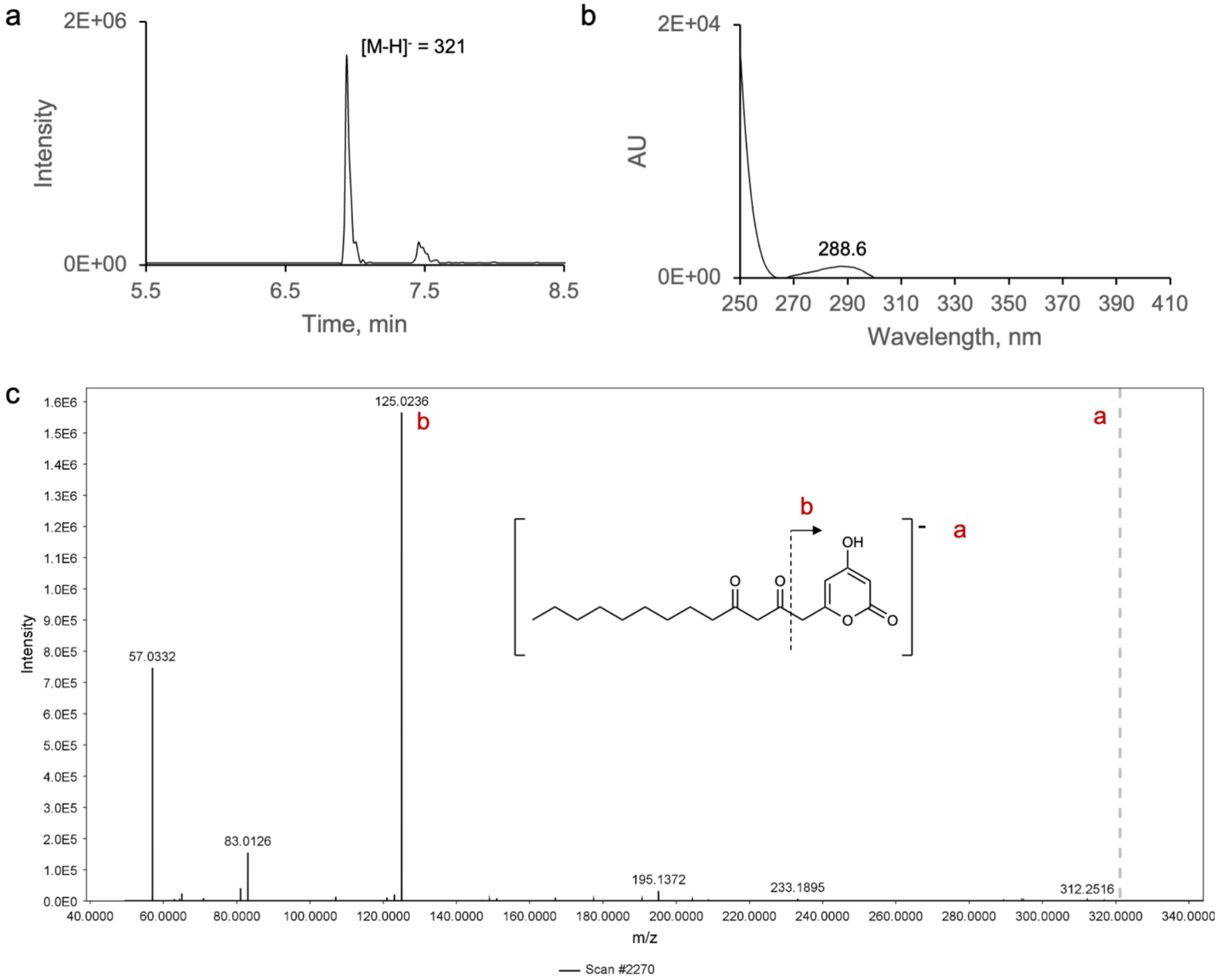

**Supplementary Figure 25.** Spectral data analysis of product **8c**. a) Low-resolution LC-MS analysis of the EtOAc extract of the enzymatic reaction of AsesPKS with **8**; extracted ion chromatogram of the predicted *m/z* of 321 (negative mode). Y-axis shows relative ion intensity. b) Corresponding UV absorption spectrum. c) ESI-HR-MS/MS (negative mode) with ions matching expected fragments of **8c**; observed *m/z* = 321.1711 (theoretical *m/z* = 321.1707, calculated for $[C_{18}H_{25}O_5]^-$). The precursor ion is indicated with a dashed grey line.

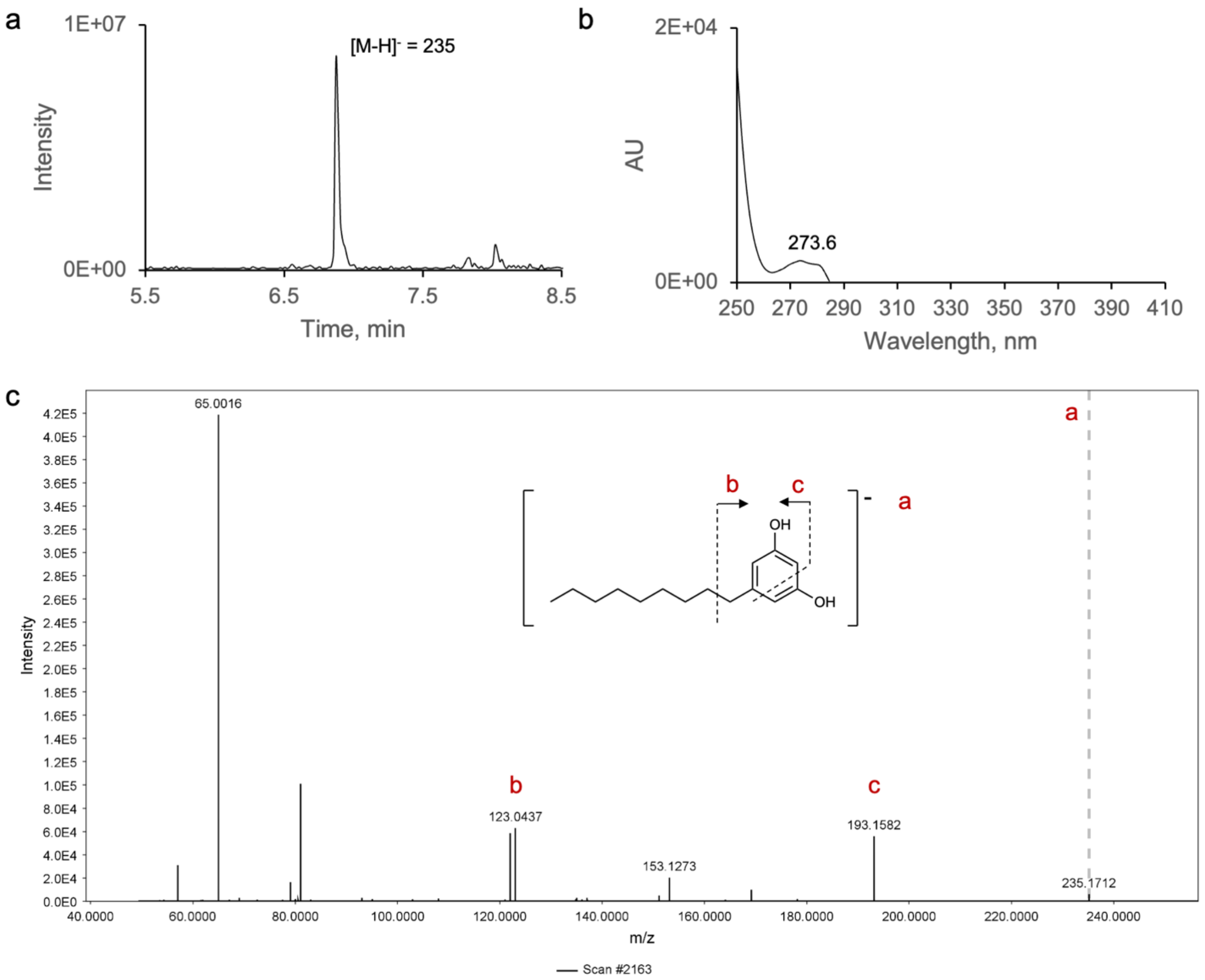

**Supplementary Figure 26.** Spectral data analysis of product **8d**. a) Low-resolution LC-MS analysis of the EtOAc extract of the enzymatic reaction of DhelPKS with **8**; extracted ion chromatogram of the predicted *m/z* of 235 (negative mode). Y-axis shows relative ion intensity. b) Corresponding UV absorption spectrum. c) ESI-HR-MS/MS (negative mode) with ions matching expected fragments of **8d**; observed *m/z* = 235.1701 (theoretical *m/z* = 235.1704, calculated for $[C_{15}H_{23}O_2]^-$). The precursor ion is indicated with a dashed grey line.

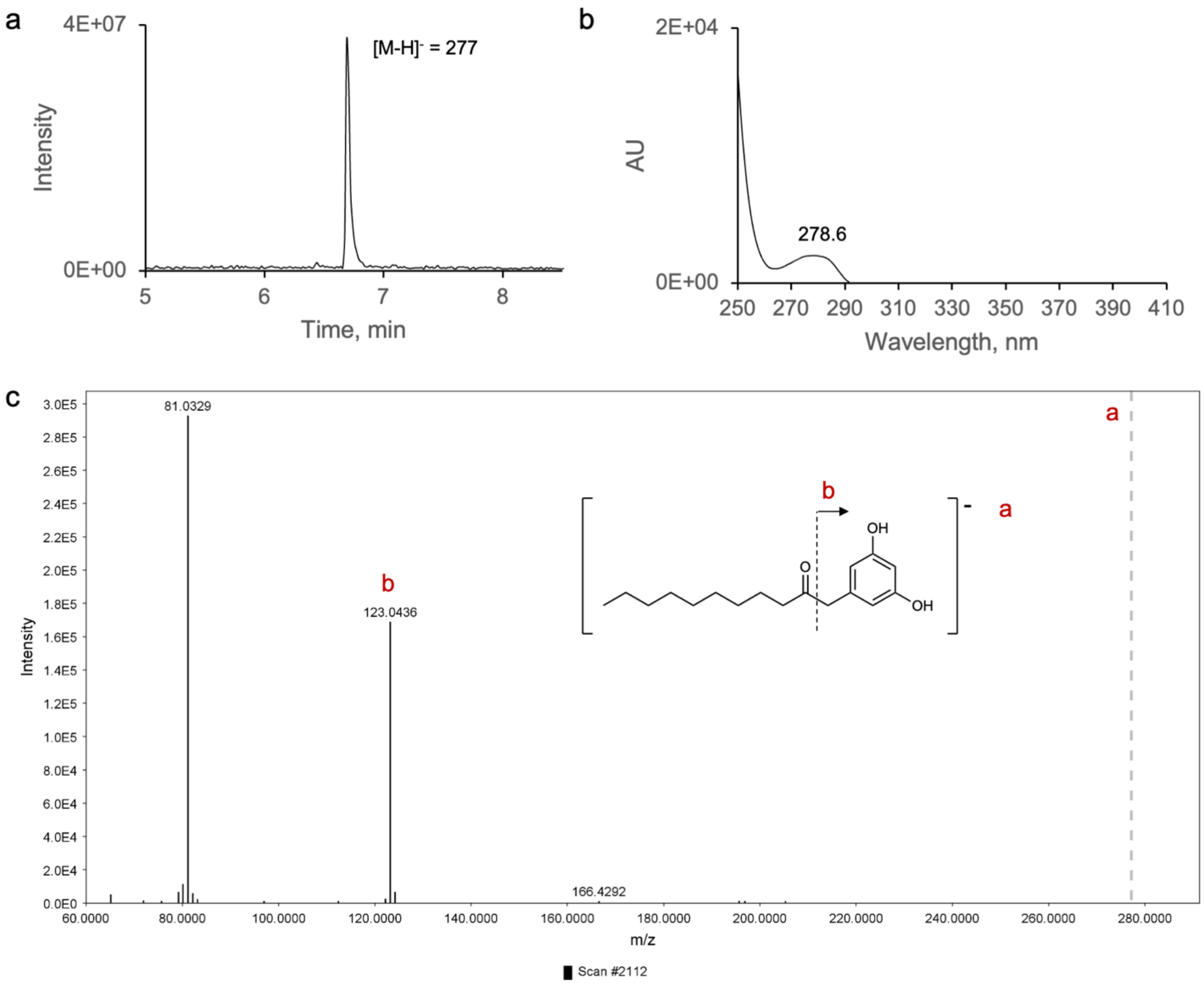

**Supplementary Figure 27.** Spectral data analysis of product **8e**. a) Low-resolution LC-MS analysis of the EtOAc extract of the enzymatic reaction of AcosPKS with **8**; extracted ion chromatogram of the predicted *m/z* of 277 (negative mode). Y-axis shows relative ion intensity. b) Corresponding UV absorption spectrum. c) ESI-HR-MS/MS (negative mode) with ions matching expected fragments of **8e**; observed *m/z* = 277.1805 (theoretical *m/z* = 277.1809, calculated for $[C_{17}H_{25}O_3]^-$). The precursor ion is indicated with a dashed grey line.

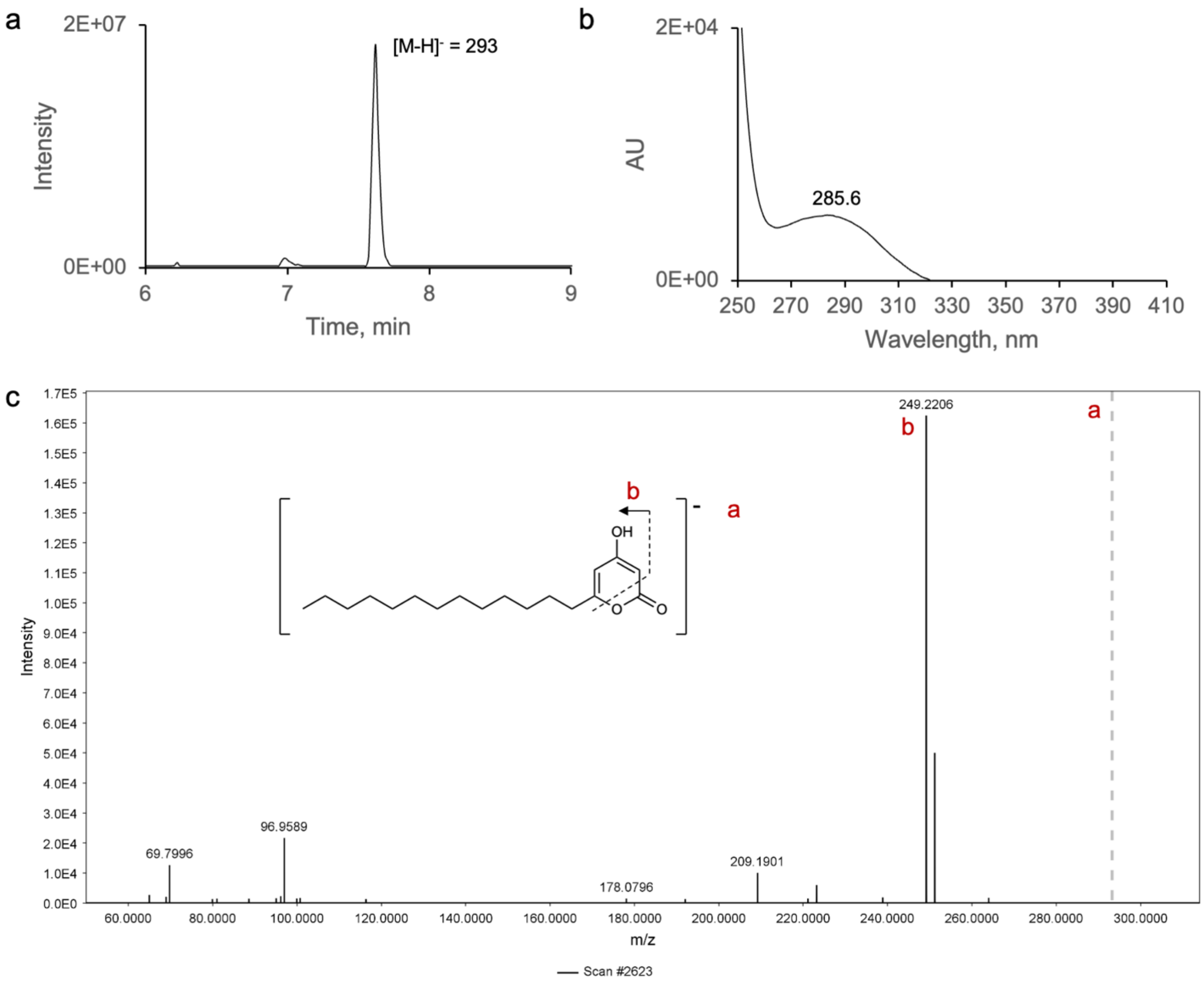

**Supplementary Figure 28.** Spectral data analysis of product **9a**. a) Low-resolution LC-MS analysis of the EtOAc extract of the enzymatic reaction of DhelPKS with **9**; extracted ion chromatogram of the predicted *m/z* of 293 (negative mode). Y-axis shows relative ion intensity. b) Corresponding UV absorption spectrum. c) ESI-HR-MS/MS (negative mode) with ions matching expected fragments of **9a**; observed *m/z* = 293.2127 (theoretical *m/z* = 293.2122, calculated for $[C_{18}H_{29}O_3]^-$). The precursor ion is indicated with a dashed grey line.

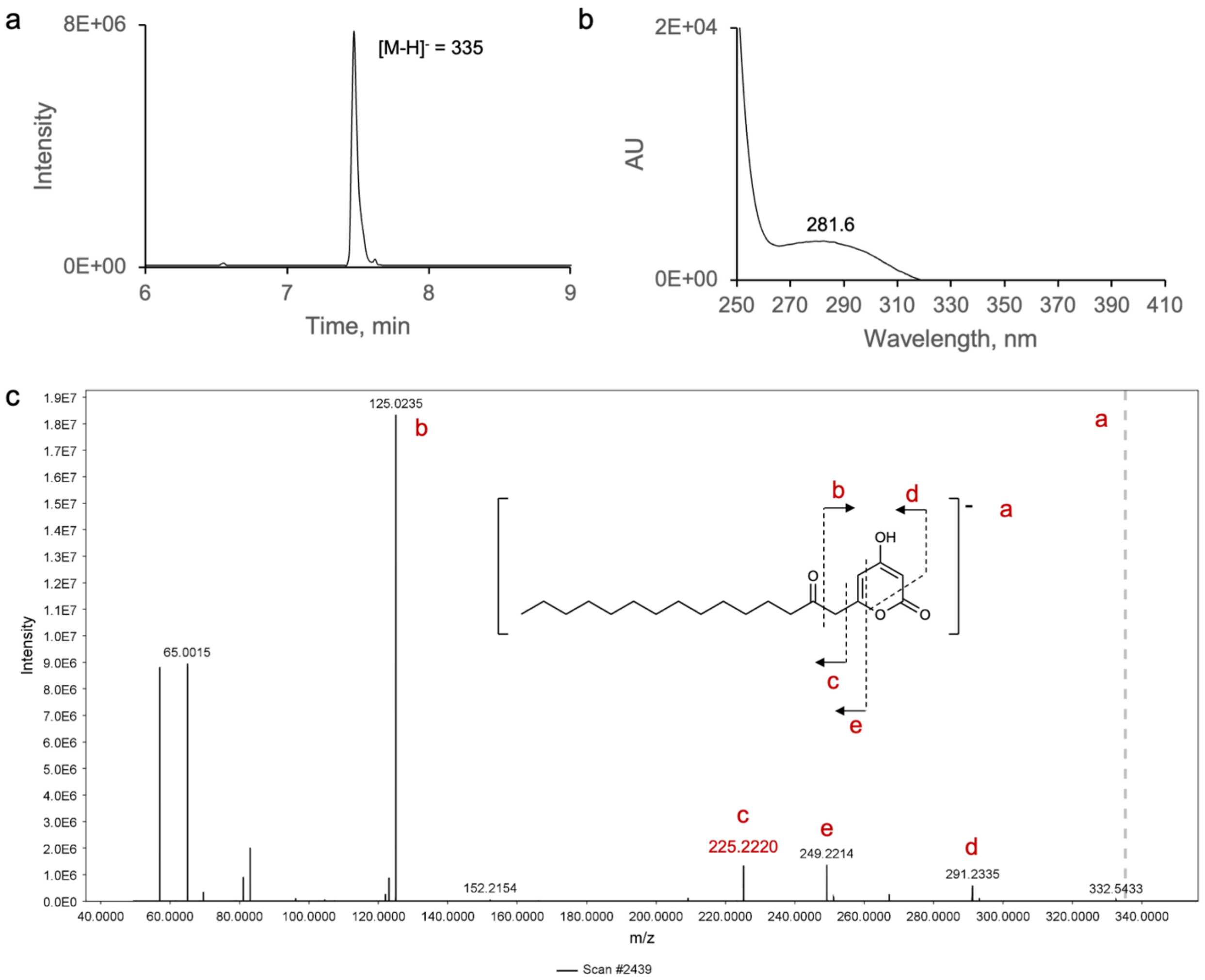

**Supplementary Figure 29.** Spectral data analysis of product **9b**. a) Low-resolution LC-MS analysis of the EtOAc extract of the enzymatic reaction of DhelPKS with **9**; extracted ion chromatogram of the predicted *m/z* of 335 (negative mode). Y-axis shows relative ion intensity. b) Corresponding UV absorption spectrum. c) ESI-HR-MS/MS (negative mode) with ions matching expected fragments of **9b**; observed *m/z* = 335.2231 (theoretical *m/z* = 335.2228, calculated for $[C_{20}H_{31}O_4]^-$). The precursor ion is indicated with a dashed grey line.

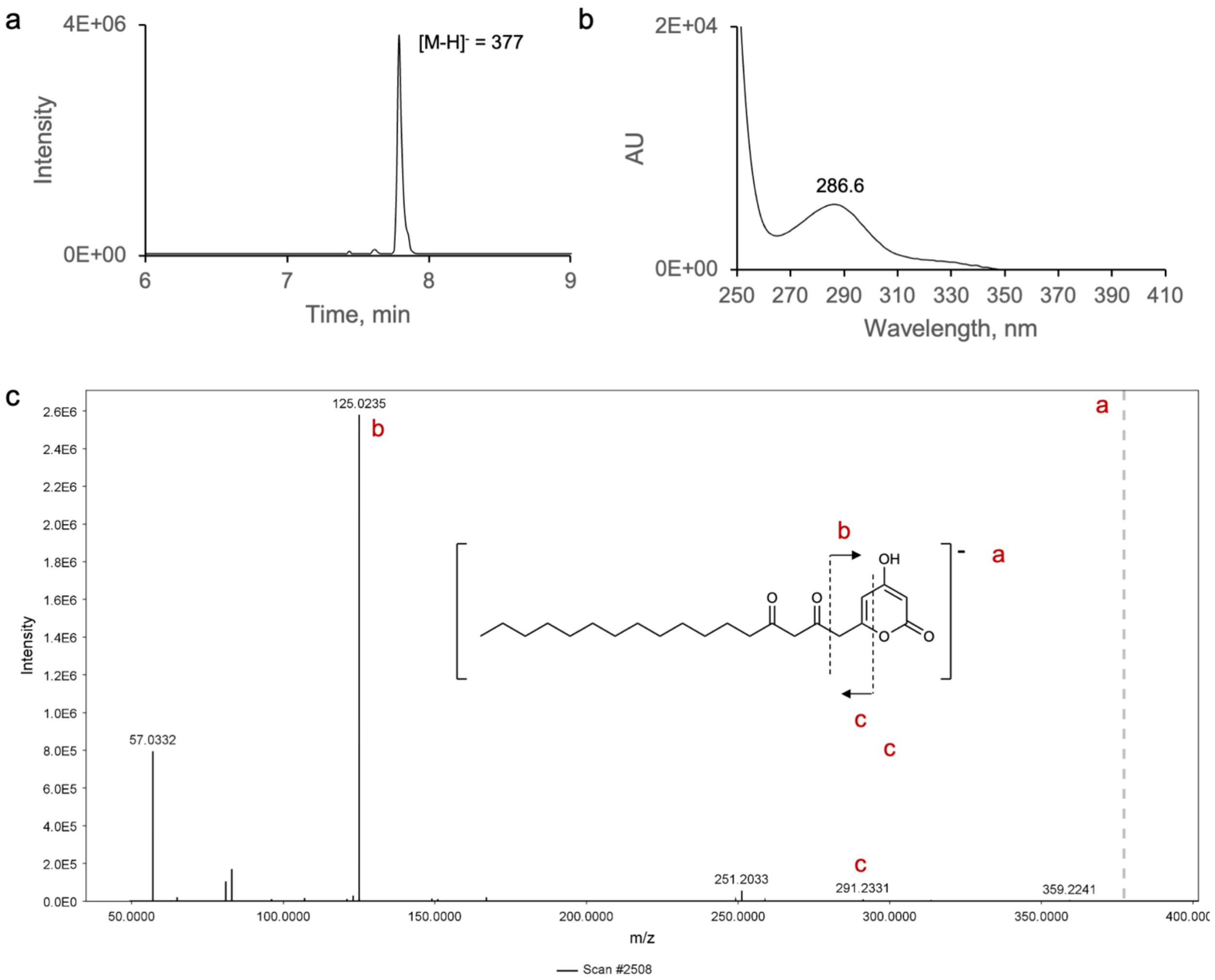

**Supplementary Figure 30.** Spectral data analysis of product **9c**. a) Low-resolution LC-MS analysis of the EtOAc extract of the enzymatic reaction of HargPKS1 with **9**; extracted ion chromatogram of the predicted *m/z* of 377 (negative mode). Y-axis shows relative ion intensity. b) Corresponding UV absorption spectrum. c) ESI-HR-MS/MS (negative mode) with ions matching expected fragments of **9c**; observed *m/z* = 377.2338 (theoretical *m/z* = 377.2333, calculated for $[C_{22}H_{33}O_5]^-$). The precursor ion is indicated with a dashed grey line.

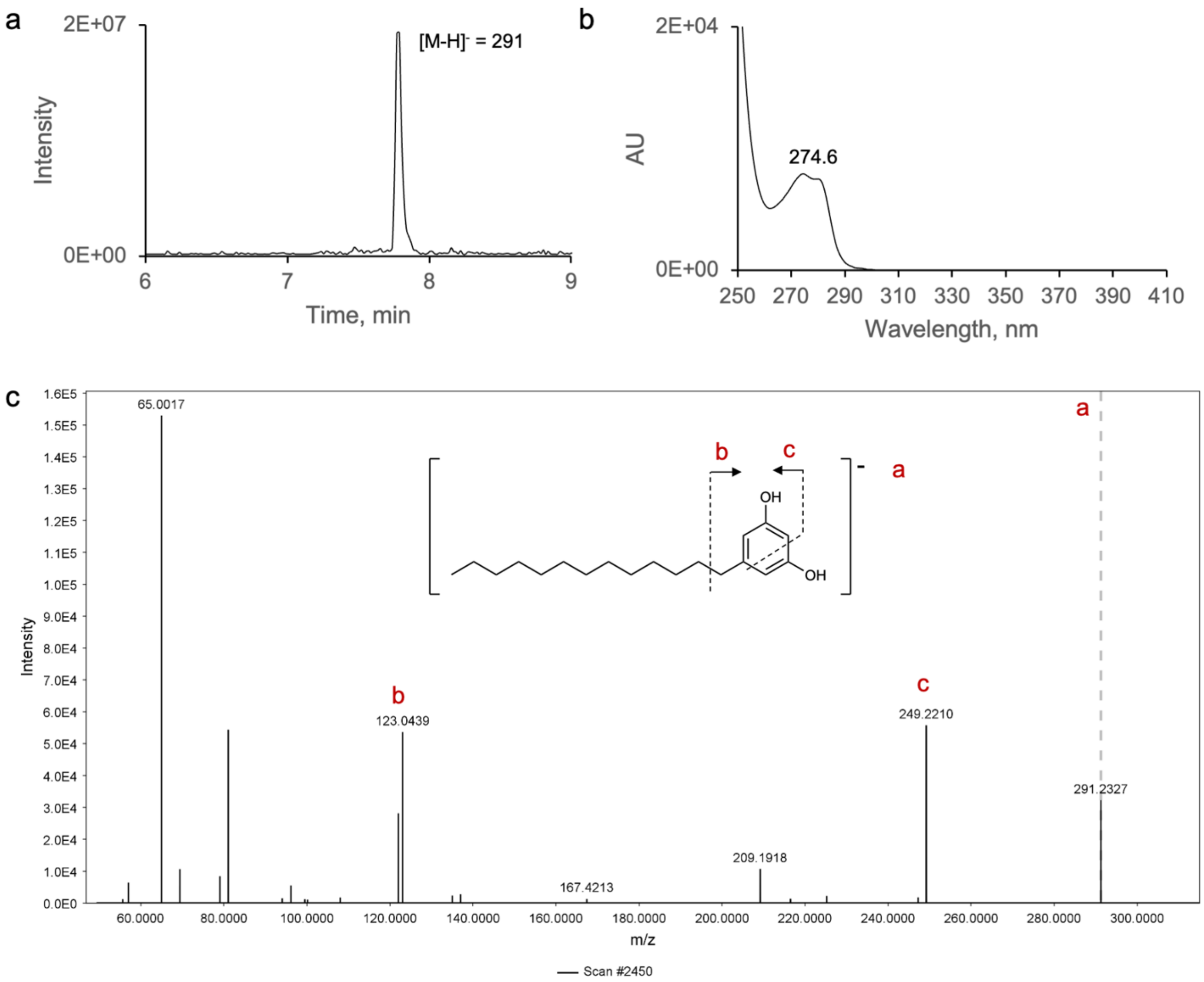

**Supplementary Figure 31.** Spectral data analysis of product **9d**. a) Low-resolution LC-MS analysis of the EtOAc extract of the enzymatic reaction of DhelPKS with **9**; extracted ion chromatogram of the predicted *m/z* of 291 (negative mode). Y-axis shows relative ion intensity. b) Corresponding UV absorption spectrum. c) ESI-HR-MS/MS (negative mode) with ions matching expected fragments of **9d**; observed *m/z* = 291.2330 (theoretical *m/z* = 291.2330, calculated for $[C_{19}H_{31}O_2]^-$). The precursor ion is indicated with a dashed grey line.

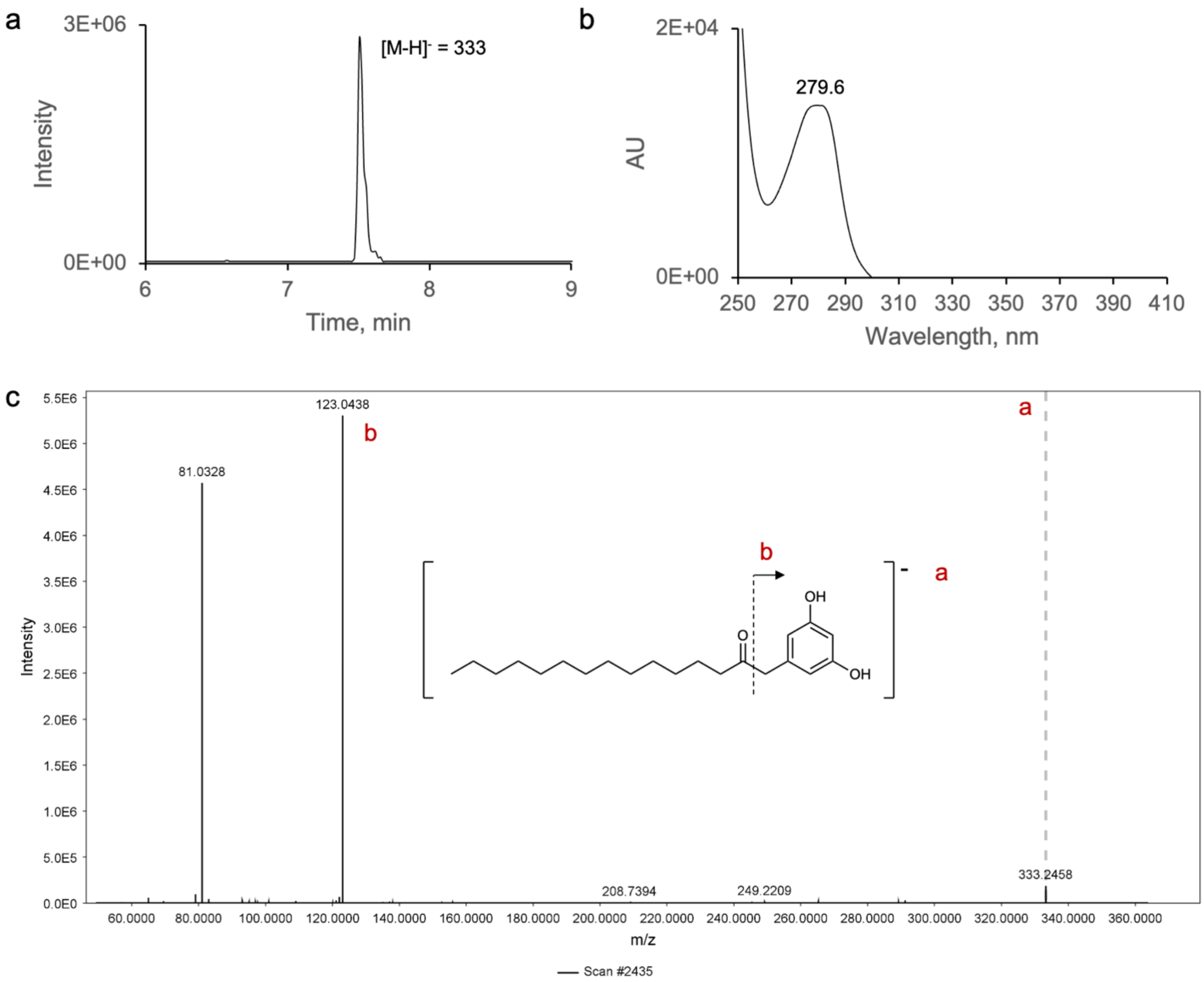

**Supplementary Figure 32.** Spectral data analysis of product **9e**. a) Low-resolution LC-MS analysis of the EtOAc extract of the enzymatic reaction of AserPKS2 with **9**; extracted ion chromatogram of the predicted *m/z* of 333 (negative mode). Y-axis shows relative ion intensity. b) Corresponding UV absorption spectrum. c) ESI-HR-MS/MS (negative mode) with ions matching expected fragments of **9e**; observed *m/z* = 333.2438 (theoretical *m/z* = 333.2435, calculated for $[C_{21}H_{33}O_3]^-$). The precursor ion is indicated with a dashed grey line.

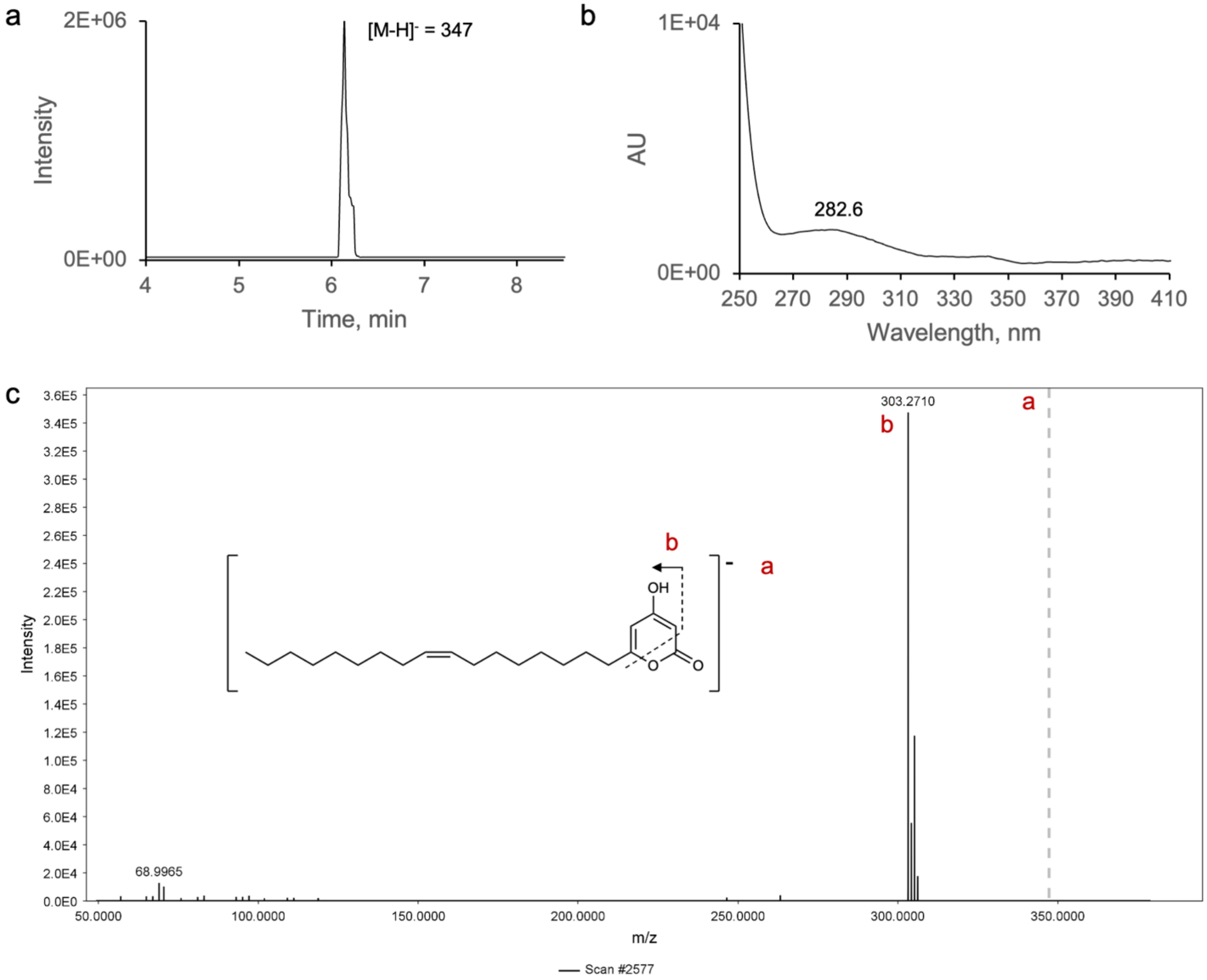

**Supplementary Figure 33.** Spectral data analysis of product **10a**. a) Low-resolution LC-MS analysis of the EtOAc extract of the enzymatic reaction of VmalPKS with **10**; extracted ion chromatogram of the predicted *m/z* of 347 (negative mode). Y-axis shows relative ion intensity. b) Corresponding UV absorption spectrum. c) ESI-HR-MS/MS (negative mode) with ions matching expected fragments of **10a**; observed *m/z* = 347.2590 (theoretical *m/z* = 347.2592, calculated for $[C_{22}H_{35}O_3]^-$). The precursor ion is indicated with a dashed grey line.

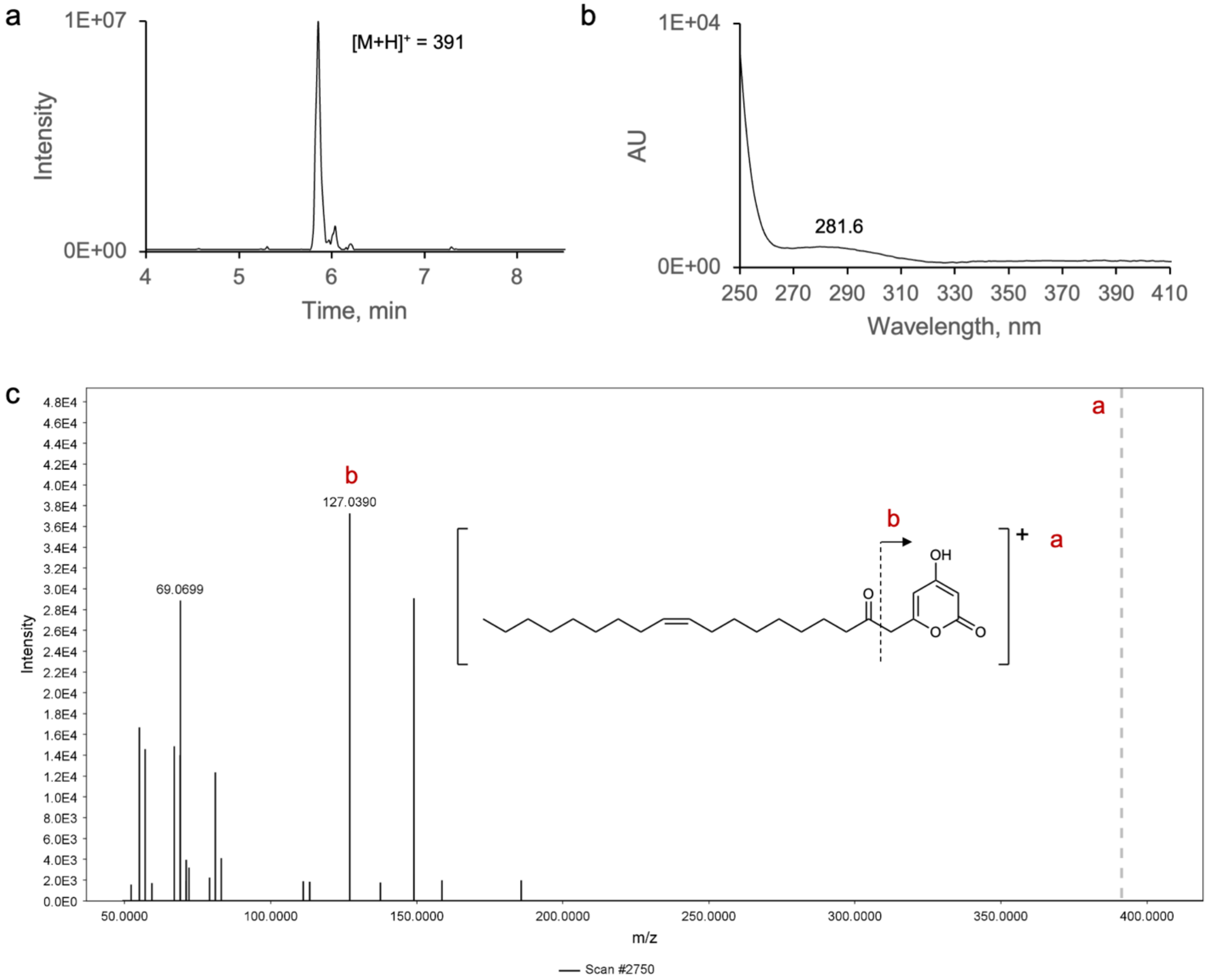

**Supplementary Figure 34.** Spectral data analysis of product **10b**. a) Low-resolution LC-MS analysis of the EtOAc extract of the enzymatic reaction of AastPKS with **10**; extracted ion chromatogram of the predicted *m/z* of 391 (positive mode). Y-axis shows relative ion intensity. b) Corresponding UV absorption spectrum. c) ESI-HR-MS/MS (positive mode) with ions matching expected fragments of **10b**; observed *m/z* = 391.2847 (theoretical *m/z* = 391.2843, calculated for $[C_{24}H_{39}O_4]^+$). The precursor ion is indicated with a dashed grey line.

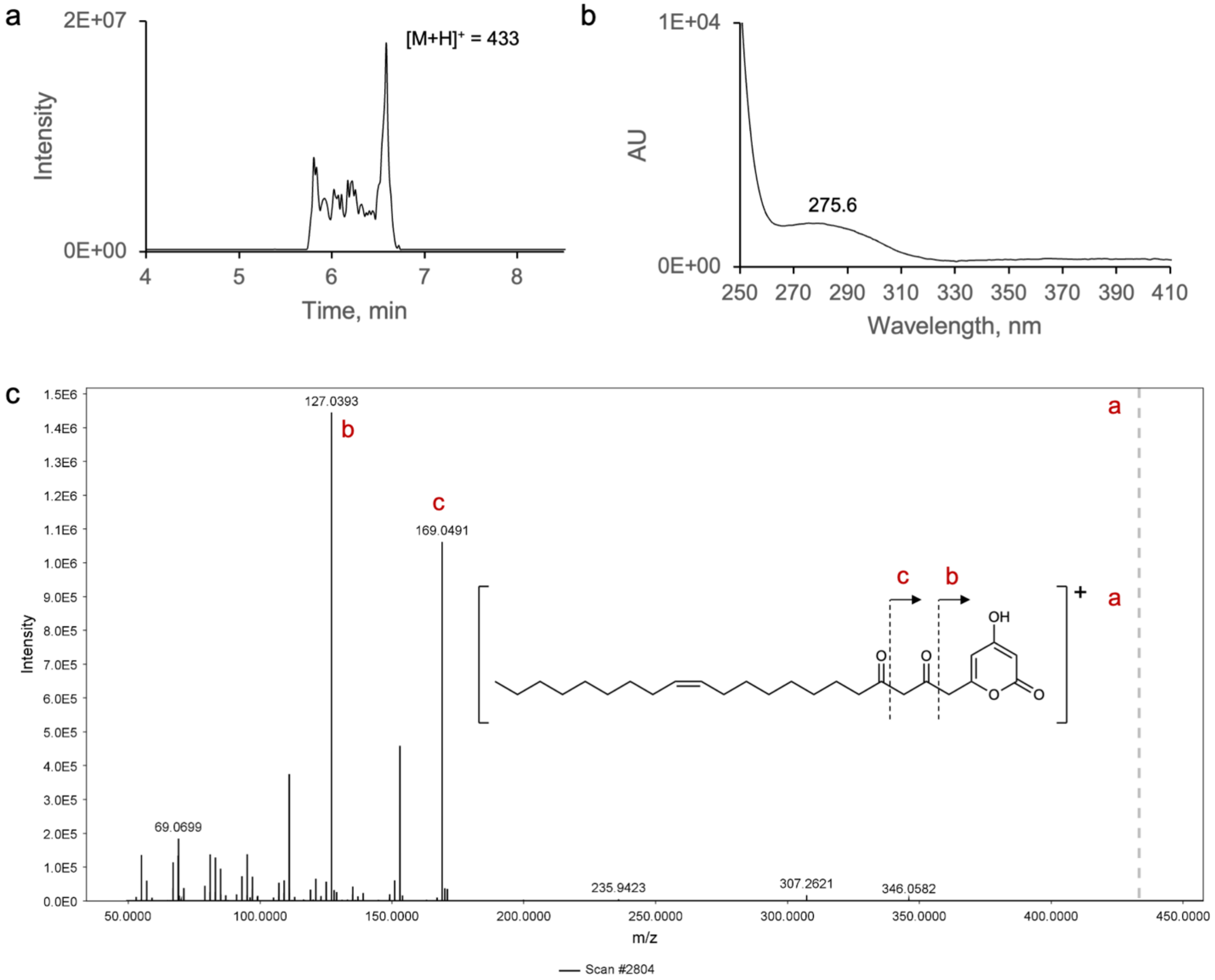

**Supplementary Figure 35.** Spectral data analysis of product **10c**. a) Low-resolution LC-MS analysis of the EtOAc extract of the enzymatic reaction of AastPKS with **10**; extracted ion chromatogram of the predicted *m/z* of 433 (positive mode). Y-axis shows relative ion intensity. b) Corresponding UV absorption spectrum. c) ESI-HR-MS/MS (positive mode) with ions matching expected fragments of **10c**; observed *m/z* = 433.2955 (theoretical *m/z* = 433.2949, calculated for $[C_{26}H_{41}O_5]^+$). The precursor ion is indicated with a dashed grey line.

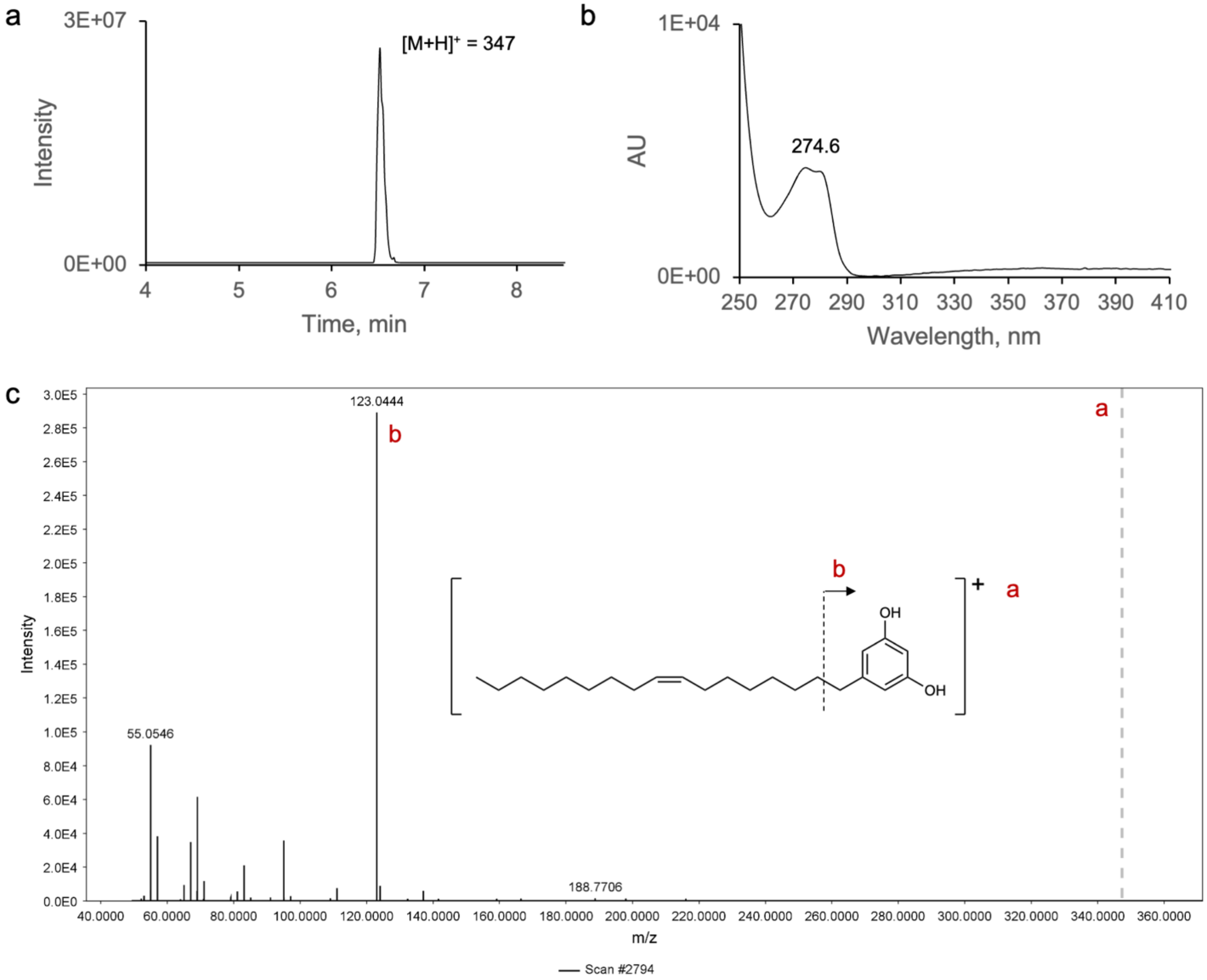

**Supplementary Figure 36.** Spectral data analysis of product **10d**. a) Low-resolution LC-MS analysis of the EtOAc extract of the enzymatic reaction of AserPKS2 with **10**; extracted ion chromatogram of the predicted *m/z* of 347 (positive mode). Y-axis shows relative ion intensity. b) Corresponding UV absorption spectrum. c) ESI-HR-MS/MS (positive mode) with ions matching expected fragments of **10d**; observed *m/z* = 347.2948 (theoretical *m/z* = 347.2945, calculated for $[C_{23}H_{39}O_2]^+$). The precursor ion is indicated with a dashed grey line.

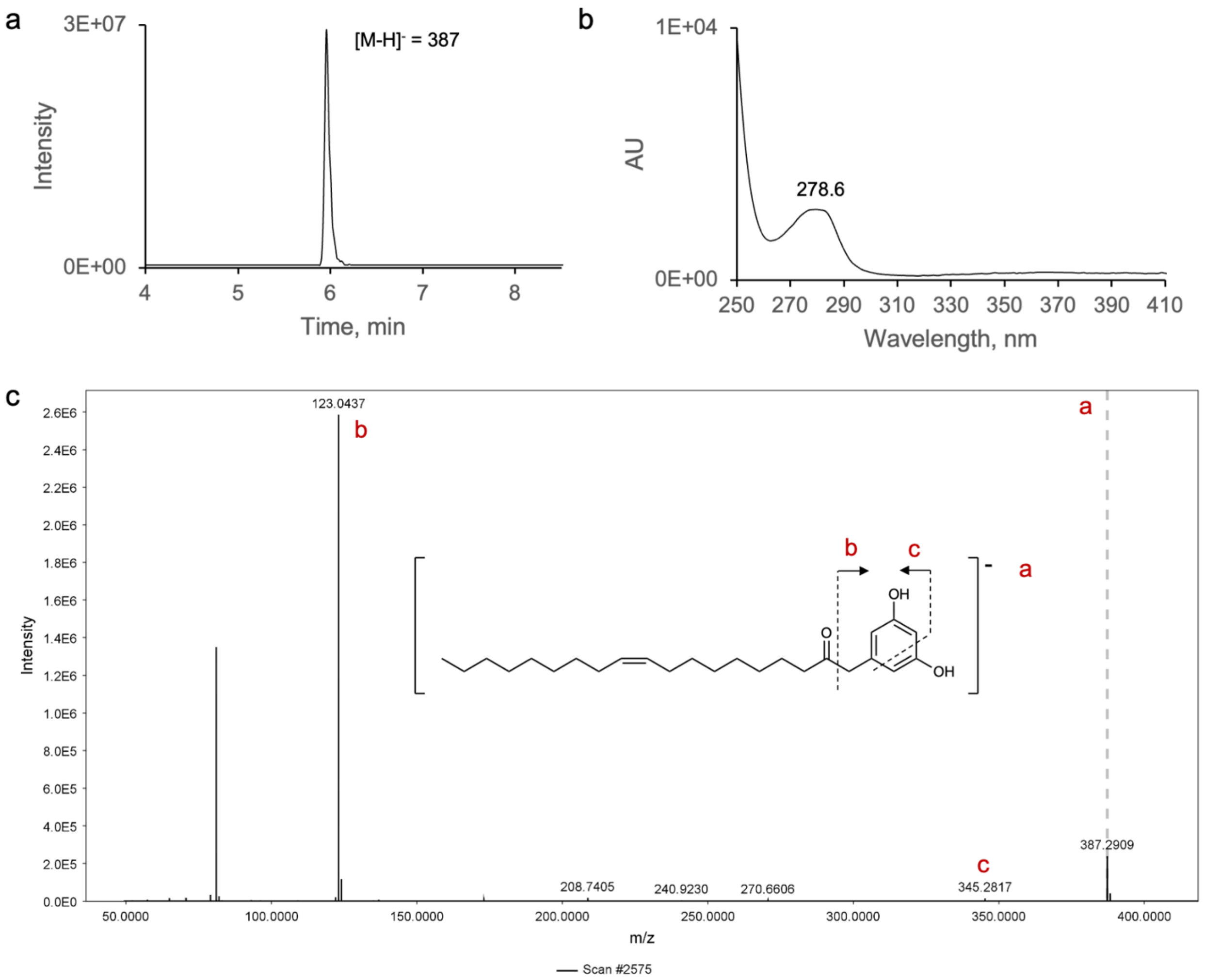

**Supplementary Figure 37.** Spectral data analysis of product **10e**. a) Low-resolution LC-MS analysis of the EtOAc extract of the enzymatic reaction of VmalPKS with **10**; extracted ion chromatogram of the predicted *m/z* of 387 (negative mode). Y-axis shows relative ion intensity. b) Corresponding UV absorption spectrum. c) ESI-HR-MS/MS (negative mode) with ions matching expected fragments of **10e**; observed *m/z* = 387.2904 (theoretical *m/z* = 387.2905, calculated for C25H39O3-). The precursor ion is indicated with a dashed grey line.

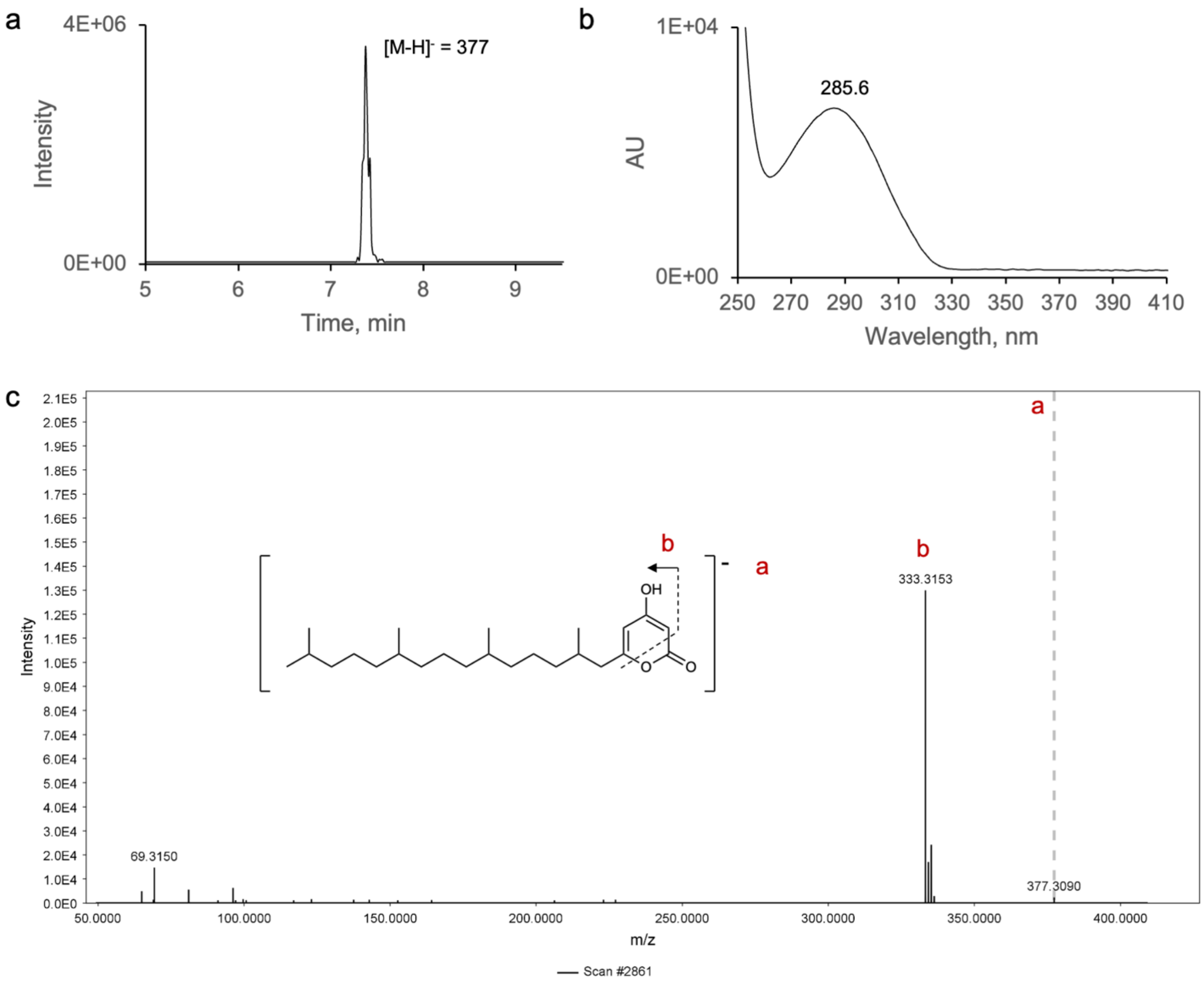

**Supplementary Figure 38.** Spectral data analysis of product **11a**. a) Low-resolution LC-MS analysis of the EtOAc extract of the enzymatic reaction of AserPKS2 with **11**; extracted ion chromatogram of the predicted *m/z* of 377 (negative mode). Y-axis shows relative ion intensity. b) Corresponding UV absorption spectrum. c) ESI-HR-MS/MS (negative mode) with ions matching expected fragments of **11a**; observed *m/z* = 377.3059 (theoretical *m/z* = 377.3061, calculated for $[C_{24}H_{41}O_3]^-$). The precursor ion is indicated with a dashed grey line.

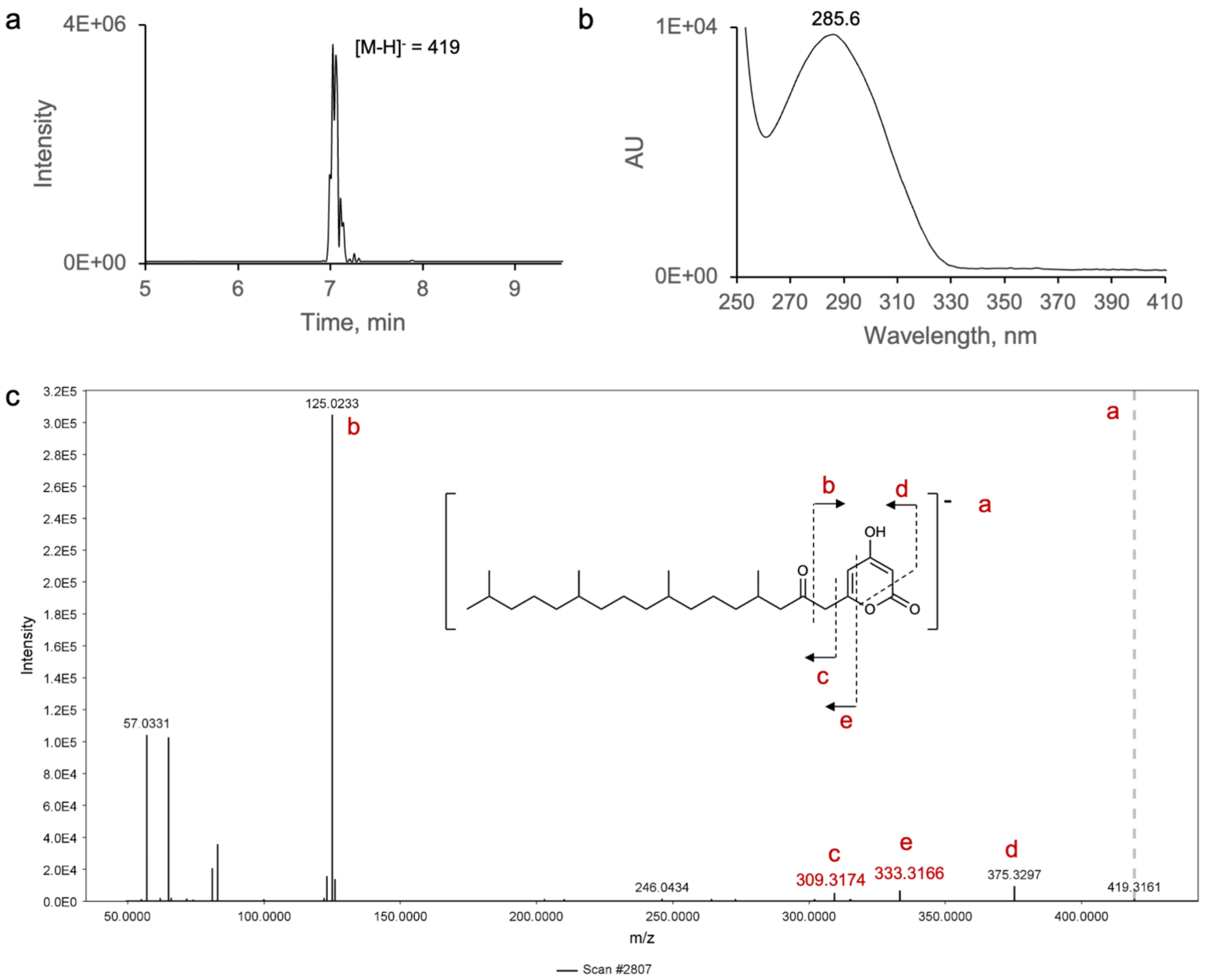

**Supplementary Figure 39.** Spectral data analysis of product **11b**. a) Low-resolution LC-MS analysis of the EtOAc extract of the enzymatic reaction of AserPKS2 with **11**; extracted ion chromatogram of the predicted *m/z* of 419 (negative mode). Y-axis shows relative ion intensity. b) Corresponding UV absorption spectrum. c) ESI-HR-MS/MS (negative mode) with ions matching expected fragments of **11b**; observed *m/z* = 419.3165 (theoretical *m/z* = 419.3167, calculated for $[C_{26}H_{43}O_4]^-$). The precursor ion is indicated with a dashed grey line.

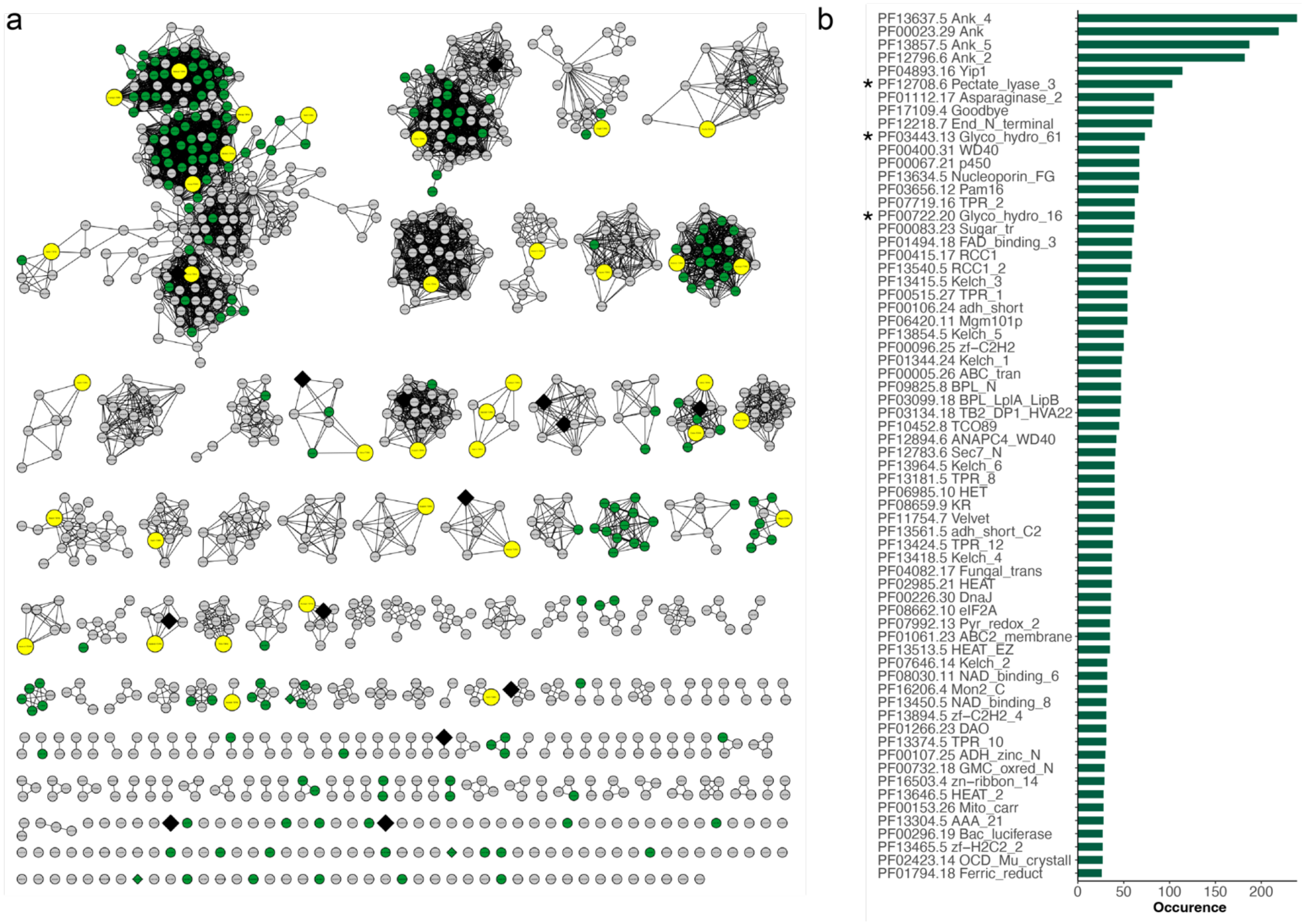

**Supplementary Figure 40.** T3PKSs originating from endophytic or phytopathogenic fungi. a) Sequence similarity network of the fungal T3PKSs with 127 sequences from endophytic or phytopathogenic fungi represented as green nodes. The sequences selected for characterisation in this study are represented as yellow nodes. b) The most abundant PFAM domains in the neighbourhood of these PKSs; putative carbohydrate-active enzymes (CAZymes) are marked with an asterisk.
