## Supplementary Data Files for "Unravelling the functional diversity of type III polyketide synthases in fungi": Supplementary_Data_7.pdf

Cluster 1–1

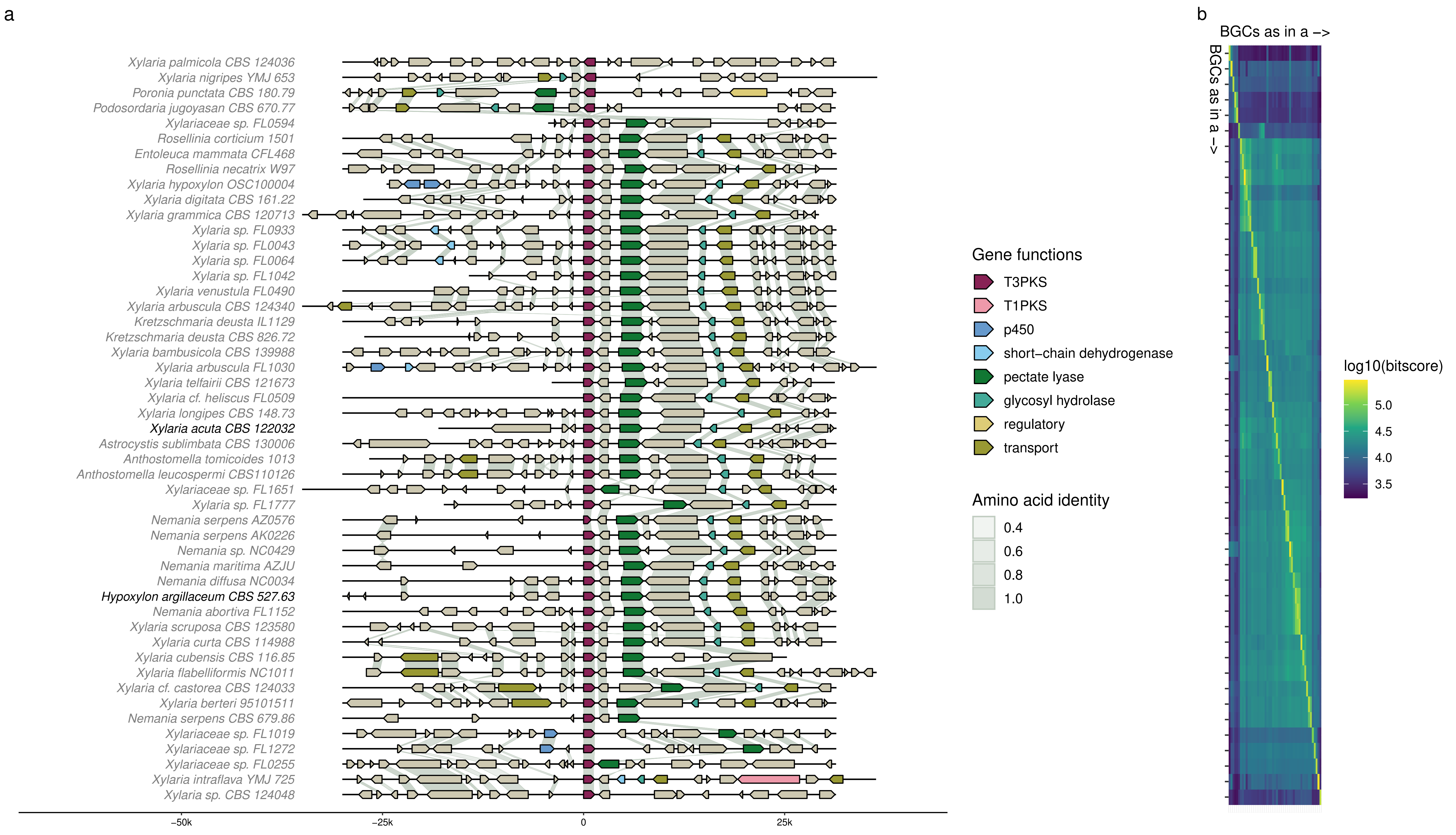
a
b
BGCs as in a ->
BGCs as in a ->
Xylaria palmicola CBS 124036
Xylaria nigripes YMJ 653
Poronia punctata CBS 180.79
Podosordaria jugoyasan CBS 670.77
Xylariaceae sp. FL0594
Rosellinia corticium 1501
Entoleuca mammata CFL468
Rosellinia necatrix W97
Xylaria hypoxylon OSC100004
Xylaria digitata CBS 161.22
Xylaria grammica CBS 120713
Xylaria sp. FL0933
Xylaria sp. FL0043
Xylaria sp. FL0064
Xylaria sp. FL1042
Xylaria venustula FL0490
Xylaria arbuscula CBS 124340
Kretzschmaria deusta IL1129
Kretzschmaria deusta CBS 826.72
Xylaria bambusicola CBS 139988
Xylaria arbuscula FL1030
Xylaria telfairii CBS 121673
Xylaria cf. heliscus FL0509
Xylaria longipes CBS 148.73
Xylaria acuta CBS 122032
Astrocystis sublimbata CBS 130006
Anthostomella tomicoides 1013
Anthostomella leucospermi CBS110126
Xylariaceae sp. FL1651
Xylaria sp. FL1777
Nemania serpens AZ0576
Nemania serpens AK0226
Nemania sp. NC0429
Nemania maritima AZJU
Nemania diffusa NC0034
Hypoxylon argillaceum CBS 527.63
Nemania abortiva FL1152
Xylaria scruposa CBS 123580
Xylaria curta CBS 114988
Xylaria cubensis CBS 116.85
Xylaria flabelliformis NC1011
Xylaria cf. castorea CBS 124033
Xylaria berteri 95101511
Nemania serpens CBS 679.86
Xylariaceae sp. FL1019
Xylariaceae sp. FL1272
Xylariaceae sp. FL0255
Xylaria intraflava YMJ 725
Xylaria sp. CBS 124048
-50k
-25k
0
25k
Gene functions
T3PKS
T1PKS
p450
short-chain dehydrogenase
pectate lyase
glycosyl hydrolase
regulatory
transport
Amino acid identity
0.4
0.6
0.8
1.0
log10(bitscore)
5.0
4.5
4.0
3.5

Cluster 1–2

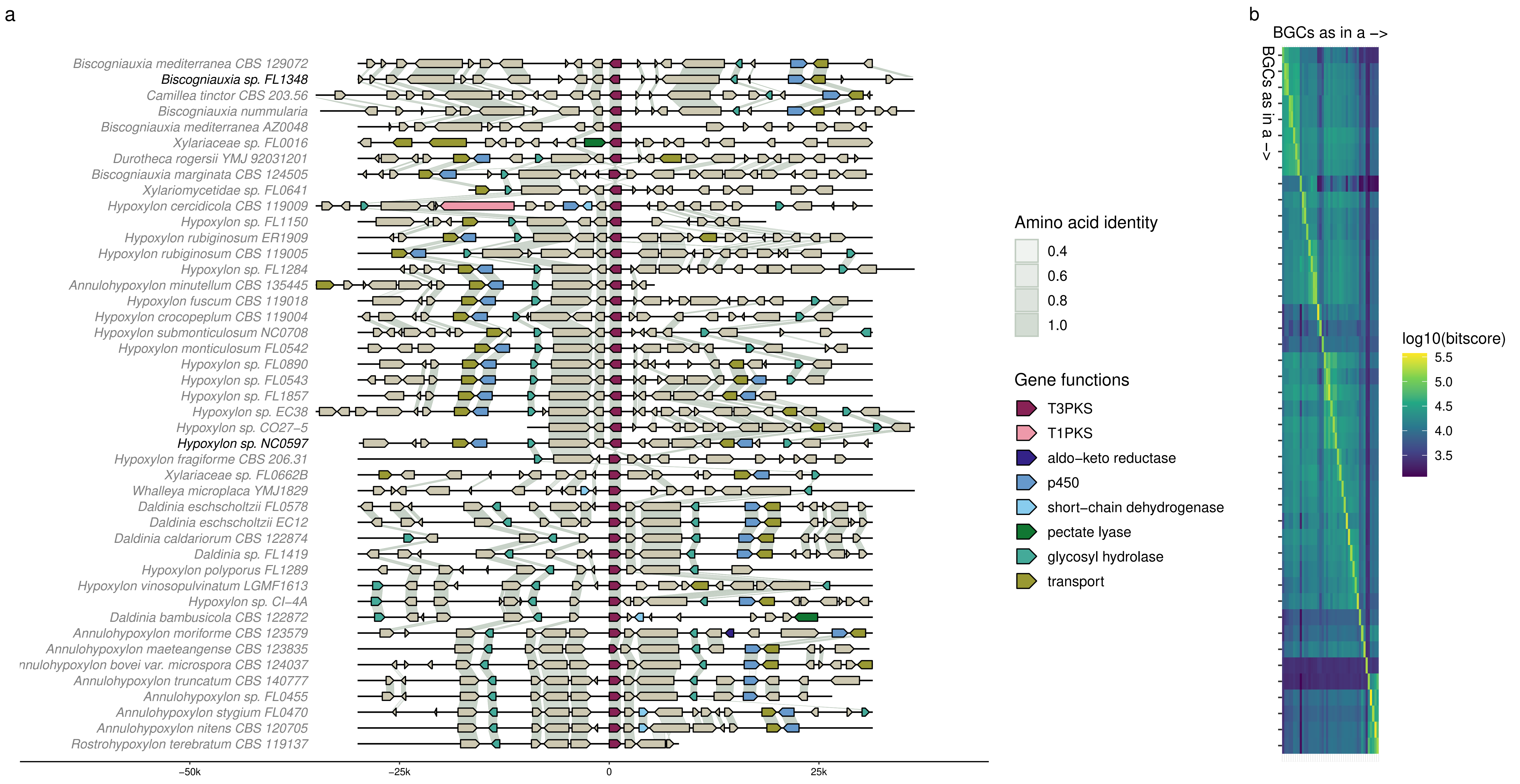
a
Biscogniauxia mediterranea CBS 129072
Biscogniauxia sp. FL1348
Camillea tinctor CBS 203.56
Biscogniauxia nummularia
Biscogniauxia mediterranea AZ0048
Xylariaceae sp. FL0016
Durotheca rogersii YMJ 92031201
Biscogniauxia marginata CBS 124505
Xylariomycetidae sp. FL0641
Hypoxylon cercidicola CBS 119009
Hypoxylon sp. FL1150
Hypoxylon rubiginosum ER1909
Hypoxylon rubiginosum CBS 119005
Hypoxylon sp. FL1284
Annulohypoxylon minutellum CBS 135445
Hypoxylon fuscum CBS 119018
Hypoxylon crocopeplum CBS 119004
Hypoxylon submonticulosum NC0708
Hypoxylon monticulosum FL0542
Hypoxylon sp. FL0890
Hypoxylon sp. FL0543
Hypoxylon sp. FL1857
Hypoxylon sp. EC38
Hypoxylon sp. CO27–5
Hypoxylon sp. NC0597
Hypoxylon fragiforme CBS 206.31
Xylariaceae sp. FL0662B
Whalleya microplaca YMJ1829
Daldinia eschscholtzii FL0578
Daldinia eschscholtzii EC12
Daldinia caldariorum CBS 122874
Daldinia sp. FL1419
Hypoxylon polyporus FL1289
Hypoxylon vinosopulvinatum LGMF1613
Hypoxylon sp. CI–4A
Daldinia bambusicola CBS 122872
Annulohypoxylon moriforme CBS 123579
Annulohypoxylon maeteangense CBS 123835
nnulohypoxylon bovei var. microspora CBS 124037
Annulohypoxylon truncatum CBS 140777
Annulohypoxylon sp. FL0455
Annulohypoxylon stygium FL0470
Annulohypoxylon nitens CBS 120705
Rostrohypoxylon terebratum CBS 119137
–50k
–25k
0
25k
Amino acid identity
0.4
0.6
0.8
1.0
Gene functions
T3PKS
T1PKS
aldo–keto reductase
p450
short–chain dehydrogenase
pectate lyase
glycosyl hydrolase
transport
b
BGCs as in a –>
BGCs as in a –>
log10(bitscore)
5.5
5.0
4.5
4.0
3.5

Cluster 1–3

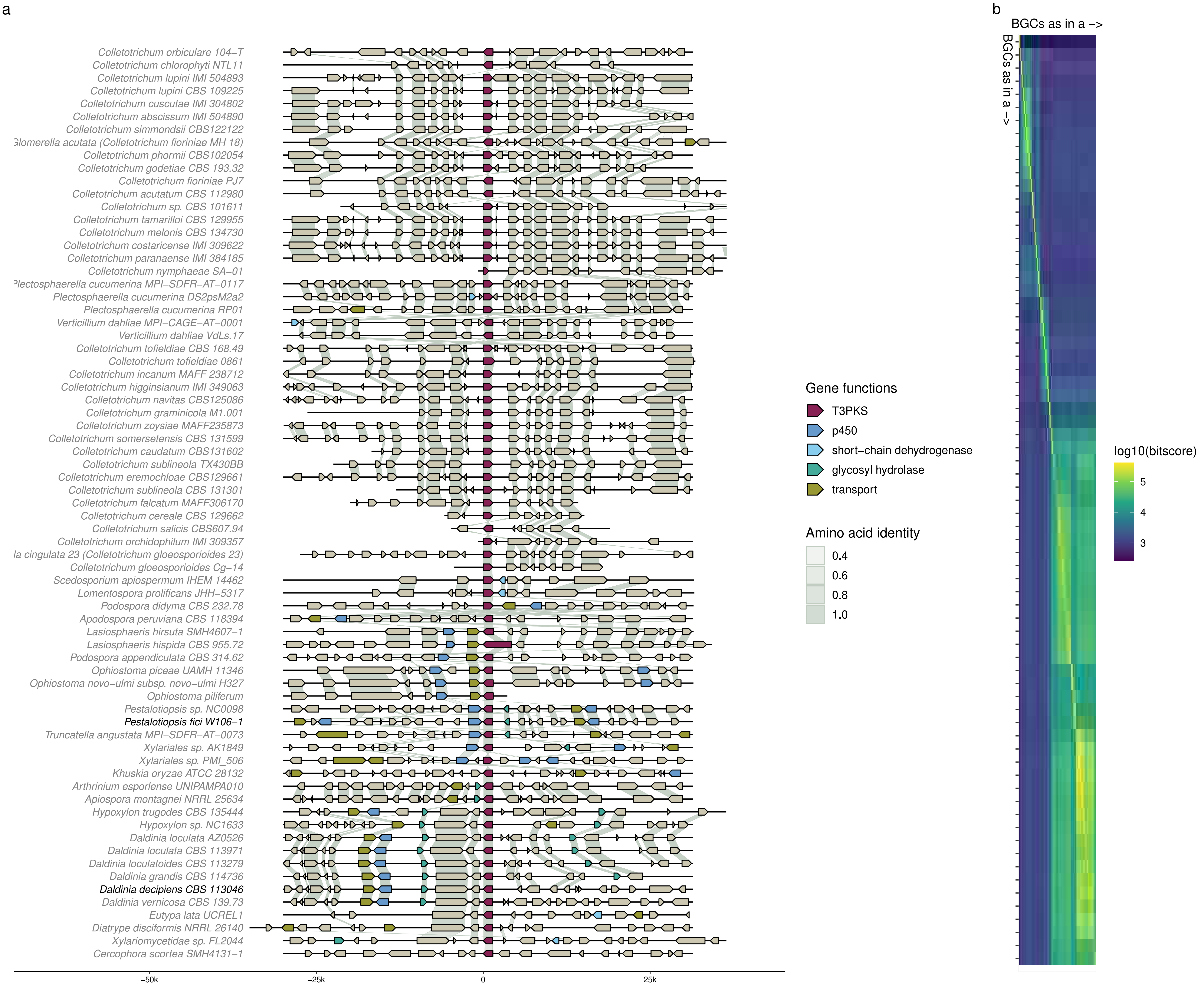
a
b
BGCs as in a –>
BGCs as in a –>
Colletotrichum orbiculare 104–T
Colletotrichum chlorophyti NTL11
Colletotrichum lupini IMI 504893
Colletotrichum lupini CBS 109225
Colletotrichum cuscutae IMI 304802
Colletotrichum abscissum IMI 504890
Colletotrichum simmondsii CBS122122
lomerella acutata (Colletotrichum fioriniae MH 18)
Colletotrichum phormii CBS102054
Colletotrichum godetiae CBS 193.32
Colletotrichum fioriniae PJ7
Colletotrichum acutatum CBS 112980
Colletotrichum sp. CBS 101611
Colletotrichum tamarilloi CBS 129955
Colletotrichum melonis CBS 134730
Colletotrichum costaricense IMI 309622
Colletotrichum paranaense IMI 384185
Colletotrichum nymphaeae SA–01
lectosphaerella cucumerina MPI–SDFR–AT–0117
Plectosphaerella cucumerina DS2psM2a2
Plectosphaerella cucumerina RP01
Verticillium dahliae MPI–CAGE–AT–0001
Verticillium dahliae VdLs.17
Colletotrichum tofieldiae CBS 168.49
Colletotrichum tofieldiae 0861
Colletotrichum incanum MAFF 238712
Colletotrichum higginsianum IMI 349063
Colletotrichum navitas CBS125086
Colletotrichum graminicola M1.001
Colletotrichum zoysiae MAFF235873
Colletotrichum somersetensis CBS 131599
Colletotrichum caudatum CBS131602
Colletotrichum sublineola TX430BB
Colletotrichum eremochloae CBS129661
Colletotrichum sublineola CBS 131301
Colletotrichum falcatum MAFF306170
Colletotrichum cereale CBS 129662
Colletotrichum salicis CBS607.94
Colletotrichum orchidophilum IMI 309357
la cingulata 23 (Colletotrichum gloeosporioides 23)
Colletotrichum gloeosporioides Cg–14
Scedosporium apiospermum IHEM 14462
Lomentospora prolificans JHH–5317
Podospora didyma CBS 232.78
Apodospora peruviana CBS 118394
Lasiosphaeris hirsuta SMH4607–1
Lasiosphaeris hispida CBS 955.72
Podospora appendiculata CBS 314.62
Ophiostoma piceae UAMH 11346
Ophiostoma novo–ulmi subsp. novo–ulmi H327
Ophiostoma piliferum
Pestalotiopsis sp. NC0098
Pestalotiopsis fici W106–1
Truncatella angustata MPI–SDFR–AT–0073
Xylariales sp. AK1849
Xylariales sp. PMI_506
Khuskia oryzae ATCC 28132
Arthrinium esporlense UNIPAMPA010
Apiospora montagnei NRRL 25634
Hypoxylon trugodes CBS 135444
Hypoxylon sp. NC1633
Daldinia loculata AZ0526
Daldinia loculata CBS 113971
Daldinia loculatoides CBS 113279
Daldinia grandis CBS 114736
Daldinia decipiens CBS 113046
Daldinia vernicosa CBS 139.73
Eutypa lata UCREL1
Diatrype disciformis NRRL 26140
Xylariomycetidae sp. FL2044
Cercophora scortea SMH4131–1
–50k
–25k
0
25k
Gene functions
T3PKS
p450
short–chain dehydrogenase
glycosyl hydrolase
transport
Amino acid identity
0.4
0.6
0.8
1.0
log10(bitscore)
5
4
3

Cluster 1–4

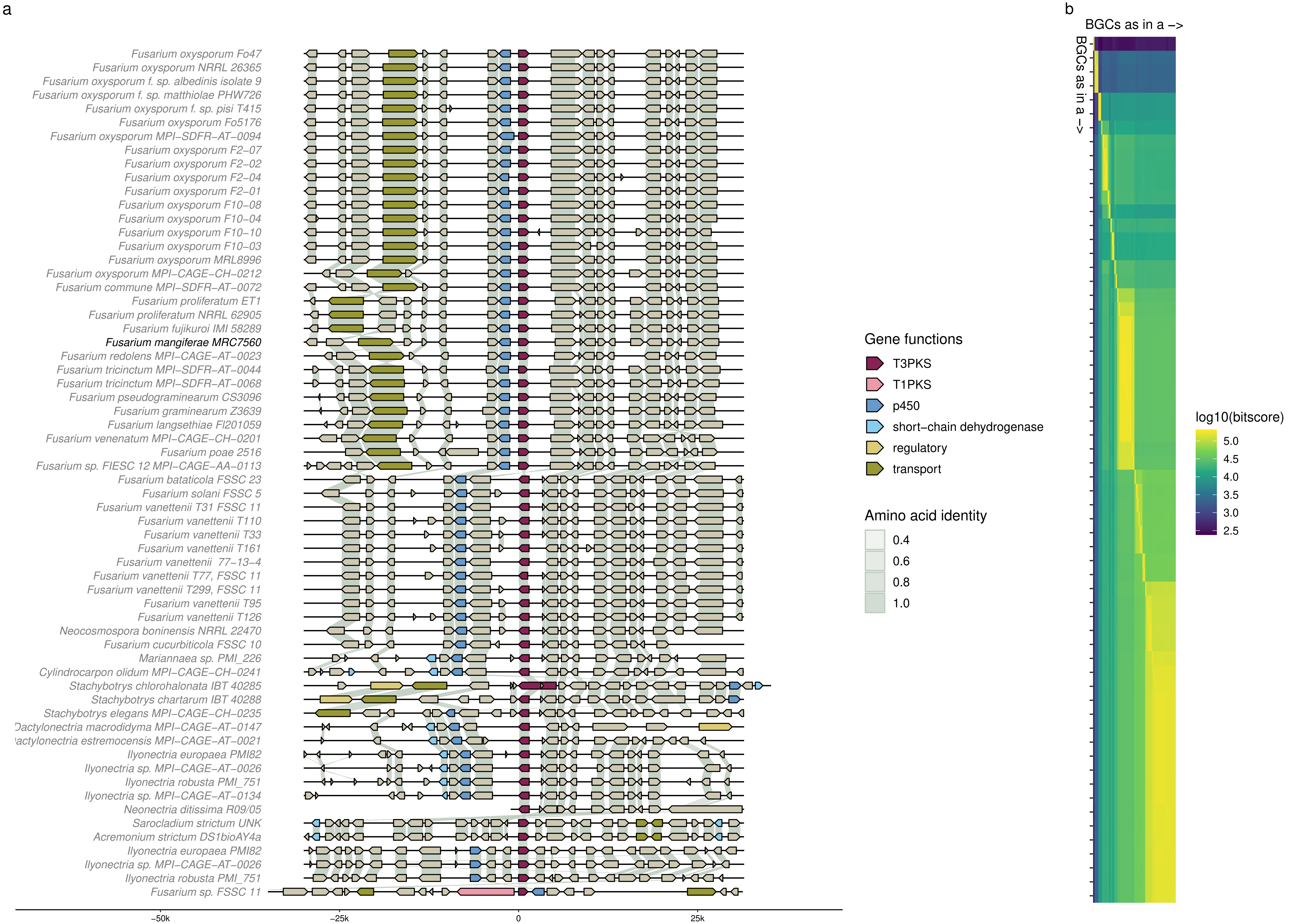
a
b
Fusarium oxysporum Fo47
Fusarium oxysporum NRRL 26365
Fusarium oxysporum f. sp. albedinis isolate 9
Fusarium oxysporum f. sp. matthiolae PHW726
Fusarium oxysporum f. sp. pisi T415
Fusarium oxysporum Fo5176
Fusarium oxysporum MPI–SDFR–AT–0094
Fusarium oxysporum F2–07
Fusarium oxysporum F2–02
Fusarium oxysporum F2–04
Fusarium oxysporum F2–01
Fusarium oxysporum F10–08
Fusarium oxysporum F10–04
Fusarium oxysporum F10–10
Fusarium oxysporum F10–03
Fusarium oxysporum MRL8996
Fusarium oxysporum MPI–CAGE–CH–0212
Fusarium commune MPI–SDFR–AT–0072
Fusarium proliferatum ET1
Fusarium proliferatum NRRL 62905
Fusarium fujikuroi IMI 58289
Fusarium mangiferae MRC7560
Fusarium redolens MPI–CAGE–AT–0023
Fusarium tricinctum MPI–SDFR–AT–0044
Fusarium tricinctum MPI–SDFR–AT–0068
Fusarium pseudograminearum CS3096
Fusarium graminearum Z3639
Fusarium langsethiae Fl201059
Fusarium venenatum MPI–CAGE–CH–0201
Fusarium poae 2516
Fusarium sp. FIESC 12 MPI–CAGE–AA–0113
Fusarium batalicola FSSC 23
Fusarium solani FSSC 5
Fusarium vanettenii T31 FSSC 11
Fusarium vanettenii T110
Fusarium vanettenii T33
Fusarium vanettenii T161
Fusarium vanettenii 77–13–4
Fusarium vanettenii T77, FSSC 11
Fusarium vanettenii T299, FSSC 11
Fusarium vanettenii T95
Fusarium vanettenii T126
Neocosmospora boninensis NRRL 22470
Fusarium cucurbiticola FSSC 10
Mariannaea sp. PMI_226
Cylindrocarpon olidum MPI–CAGE–CH–0241
Stachybotrys chlorohalonata IBT 40285
Stachybotrys chartarum IBT 40288
Stachybotrys elegans MPI–CAGE–CH–0235
Dactylonectria macrodidyma MPI–CAGE–AT–0147
actylonectria estremocensis MPI–CAGE–AT–0021
Ilyonectria europaea PMI82
Ilyonectria sp. MPI–CAGE–AT–0026
Ilyonectria robusta PMI_751
Ilyonectria sp. MPI–CAGE–AT–0134
Neonectria ditissima R09/05
Sarocladium strictum UNK
Acremonium strictum DS1bioAY4a
Ilyonectria europaea PMI82
Ilyonectria sp. MPI–CAGE–AT–0026
Ilyonectria robusta PMI_751
Fusarium sp. FSSC 11
−50k
−25k
0
25k
Gene functions
T3PKS
T1PKS
p450
short−chain dehydrogenase
regulatory
transport
Amino acid identity
0.4
0.6
0.8
1.0
BGCs as in a −>
BGCs as in a −>
log10(bitscore)
5.0
4.5
4.0
3.5
3.0
2.5

Cluster 1–5

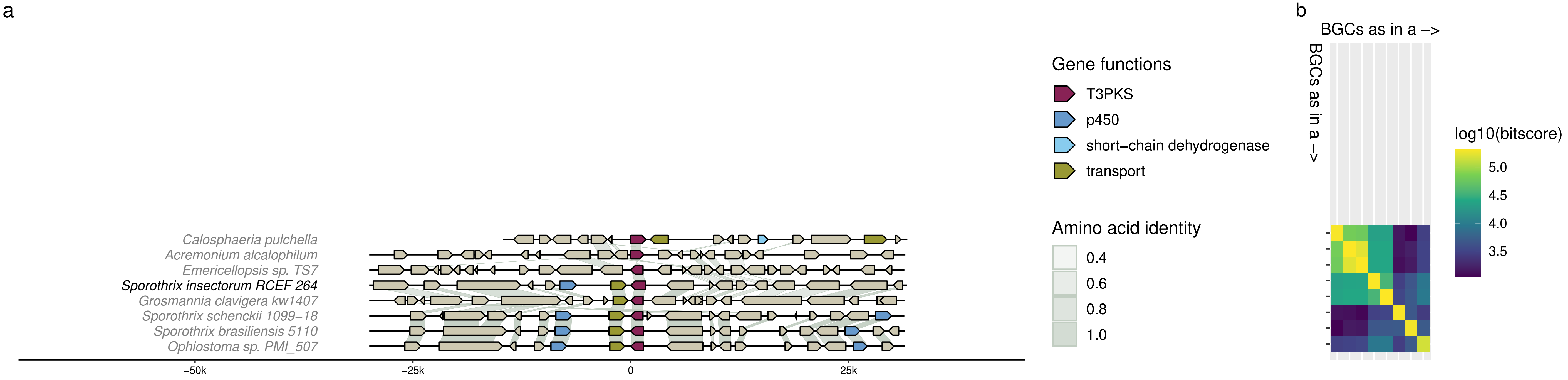
a
Calosphaeria pulchella
Acremonium alcalophilum
Emericellopsis sp. TS7
Sporothrix insectorum RCEF 264
Grosmannia clavigera kw1407
Sporothrix schenckii 1099–18
Sporothrix brasiliensis 5110
Ophiostoma sp. PMI_507
−50k
−25k
0
25k
Gene functions
T3PKS
p450
short–chain dehydrogenase
transport
Amino acid identity
0.4
0.6
0.8
1.0
b
BGCs as in a –>
BGCs as in a –>
log10(bitscore)
5.0
4.5
4.0
3.5

Cluster 2

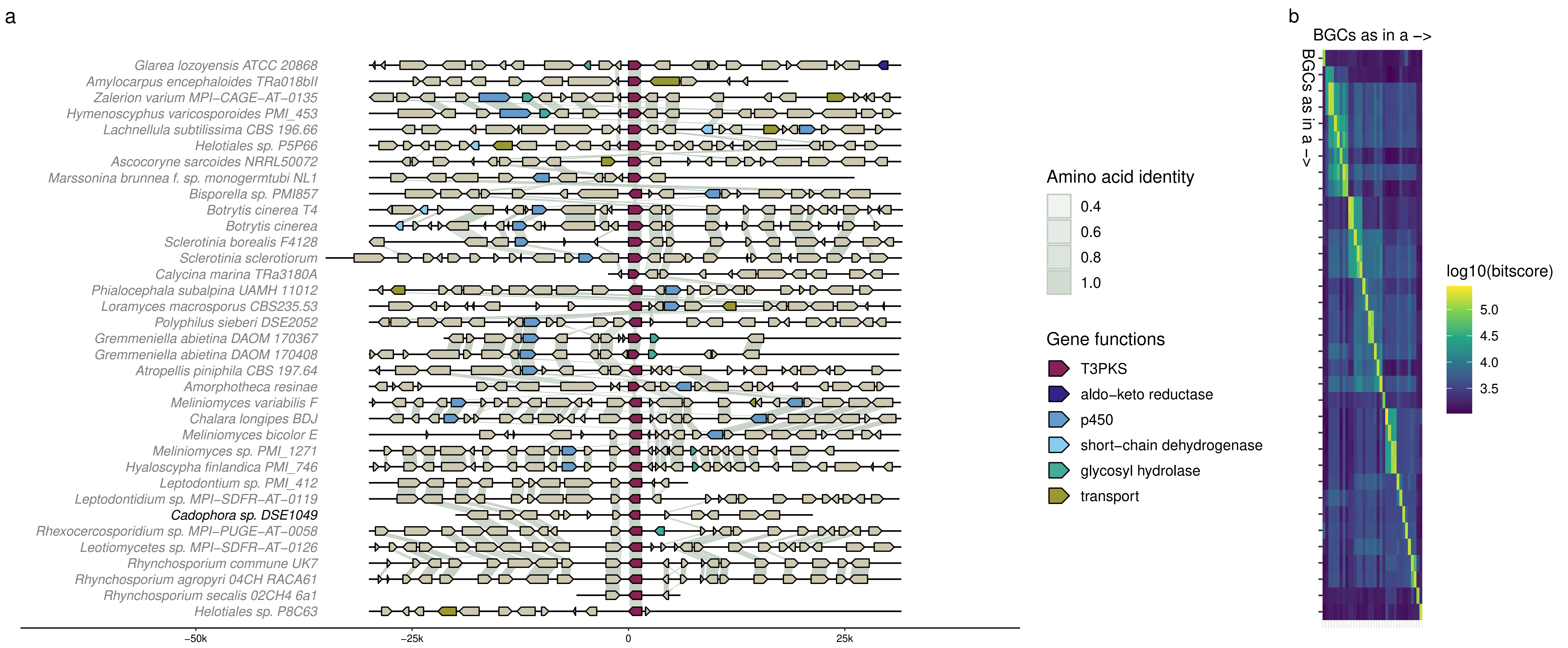

a
Glarea lozoyensis ATCC 20868
Amylocarpus encephaloides TRa018bII
Zalerion varium MPI–CAGE–AT–0135
Hymenoscyphus varicosporoides PMI_453
Lachnellula subtilissima CBS 196.66
Helotiales sp. P5P66
Ascocoryne sarcoides NRRL50072
Marssonina brunnea f. sp. monogermtubi NL1
Bisporella sp. PMI857
Botrytis cinerea T4
Botrytis cinerea
Sclerotinia borealis F4128
Sclerotinia sclerotiorum
Calycina marina TRa3180A
Phialocephala subalpina UAMH 11012
Loramyces macrosporus CBS235.53
Polyphilus sieberi DSE2052
Gremmeniella abietina DAOM 170367
Gremmeniella abietina DAOM 170408
Atropellis piniphila CBS 197.64
Amorphotheca resinae
Meliniomyces variabilis F
Chalara longipes BDJ
Meliniomyces bicolor E
Meliniomyces sp. PMI_1271
Hyaloscypha finlandica PMI_746
Leptodontium sp. PMI_412
Leptodontidium sp. MPI–SDFR–AT–0119
Cadophora sp. DSE1049
Rhexocercosporidium sp. MPI–PUGE–AT–0058
Leotiomycetes sp. MPI–SDFR–AT–0126
Rhynchosporium commune UK7
Rhynchosporium agropyri 04CH RACA61
Rhynchosporium secalis 02CH4 6a1
Helotiales sp. P8C63
–50k
–25k
0
25k
Amino acid identity
0.4
0.6
0.8
1.0
Gene functions
T3PKS
aldo–keto reductase
p450
short–chain dehydrogenase
glycosyl hydrolase
transport
b
BGCs as in a –>
BGCs as in a –>
log10(bitscore)
5.0
4.5
4.0
3.5

Cluster 3

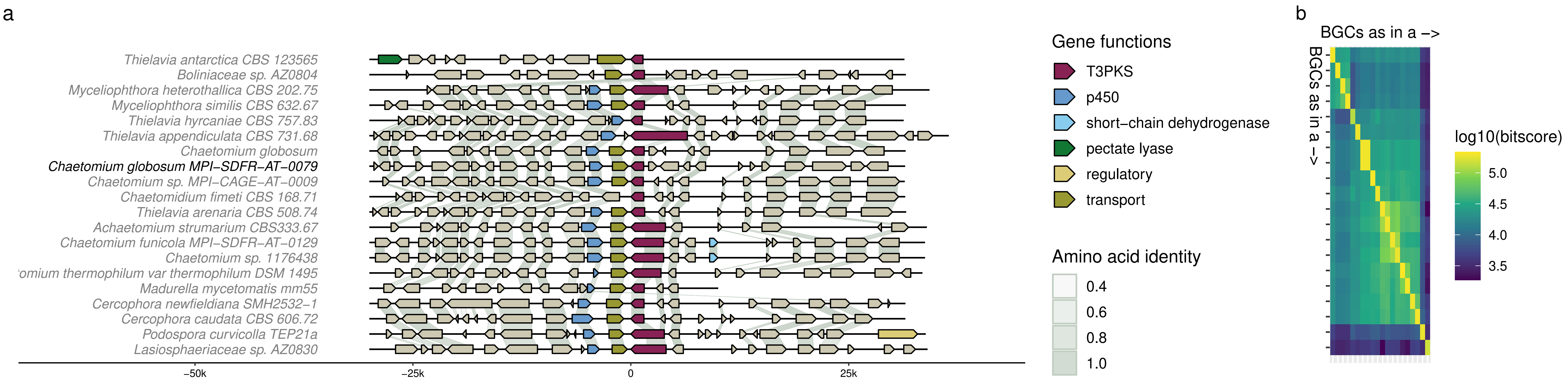
a
Thielavia antarctica CBS 123565
Boliniaceae sp. AZ0804
Myceliophthora heterothallica CBS 202.75
Myceliophthora similis CBS 632.67
Thielavia hyrcaniae CBS 757.83
Thielavia appendiculata CBS 731.68
Chaetomium globosum
Chaetomium globosum MPI–SDFR–AT–0079
Chaetomium sp. MPI–CAGE–AT–0009
Chaetomidium fimeti CBS 168.71
Thielavia arenaria CBS 508.74
Achaetomium strumarium CBS333.67
Chaetomium funicola MPI–SDFR–AT–0129
Chaetomium sp. 1176438
omium thermophilum var thermophilum DSM 1495
Madurella mycetomatis mm55
Cercophora newfieldiana SMH2532–1
Cercophora caudata CBS 606.72
Podospora curvicolla TEP21a
Lasiosphaeriaceae sp. AZ0830
–50k
–25k
0
25k
Gene functions
T3PKS
p450
short–chain dehydrogenase
pectate lyase
regulatory
transport
Amino acid identity
0.4
0.6
0.8
1.0
b
BGCs as in a –>
BGCs as in a –>
log10(bitscore)
5.0
4.5
4.0
3.5

Cluster 4

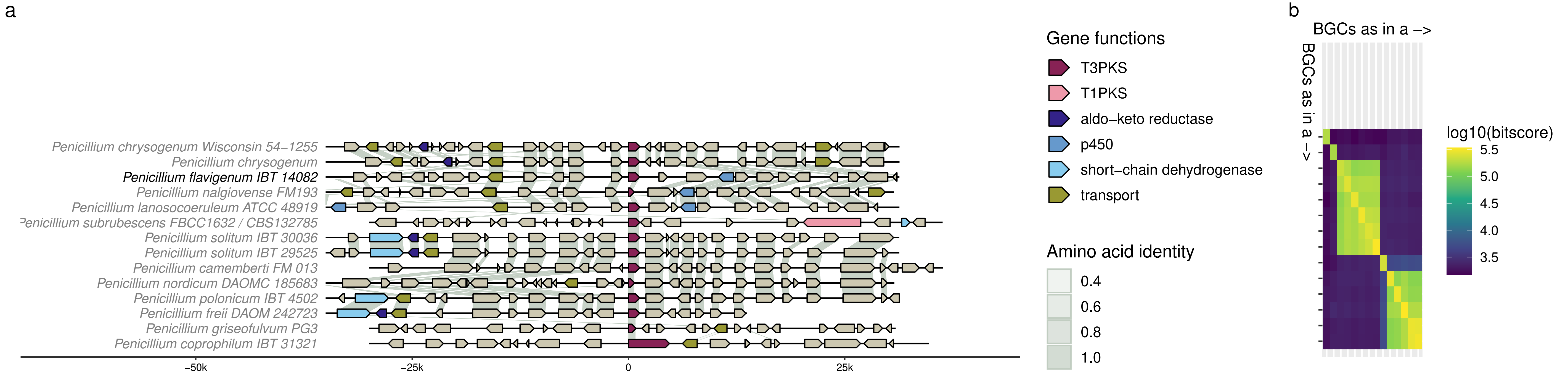
a
Penicillium chrysogenum Wisconsin 54–1255
Penicillium chrysogenum
Penicillium flavigenum IBT 14082
Penicillium nalgiovense FM193
Penicillium lanosocoeruleum ATCC 48919
Penicillium subrubescens FBCC1632 / CBS132785
Penicillium solitum IBT 30036
Penicillium solitum IBT 29525
Penicillium camemberti FM 013
Penicillium nordicum DAOMC 185683
Penicillium polonicum IBT 4502
Penicillium freii DAOM 242723
Penicillium griseofulvum PG3
Penicillium coprophilum IBT 31321
–50k
–25k
0
25k
Gene functions
T3PKS
T1PKS
aldo–keto reductase
p450
short–chain dehydrogenase
transport
Amino acid identity
0.4
0.6
0.8
1.0
b
BGCs as in a –>
BGCs as in a –>
log10(bitscore)
5.5
5.0
4.5
4.0
3.5

Cluster 5

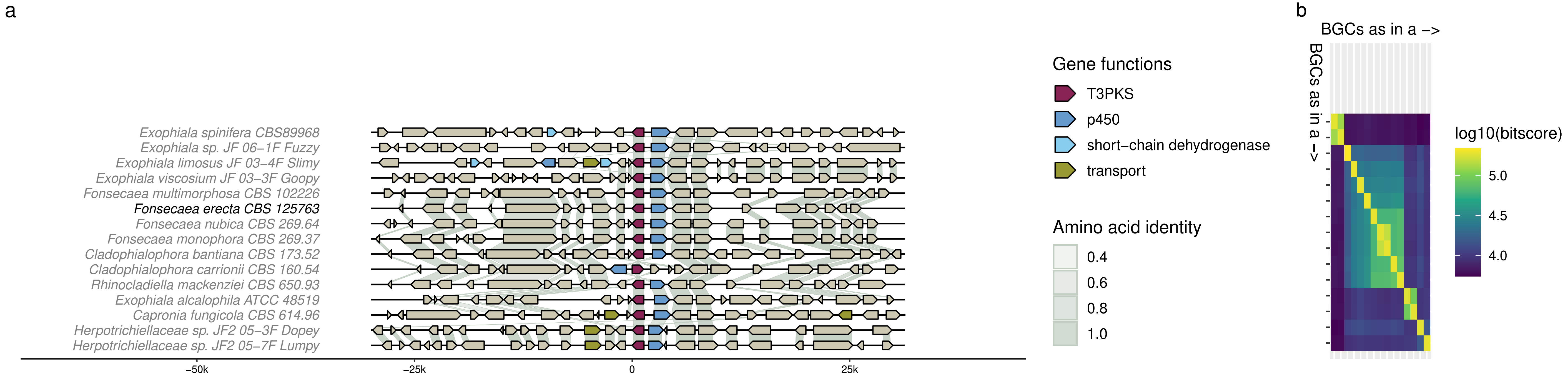

a
Exophiala spinifera CBS89968
Exophiala sp. JF 06–1F Fuzzy
Exophiala limosus JF 03–4F Slimy
Exophiala viscosium JF 03–3F Goopy
Fonsecaea multimorphosa CBS 102226
Fonsecaea erecta CBS 125763
Fonsecaea nubica CBS 269.64
Fonsecaea monophora CBS 269.37
Cladophialophora bantiana CBS 173.52
Cladophialophora carrionii CBS 160.54
Rhinocladiella mackenziei CBS 650.93
Exophiala alcalophila ATCC 48519
Capronia fungicola CBS 614.96
Herpotrichiellaceae sp. JF2 05–3F Dopey
Herpotrichiellaceae sp. JF2 05–7F Lumpy
–50k
–25k
0
25k
Gene functions
T3PKS
p450
short–chain dehydrogenase
transport
Amino acid identity
0.4
0.6
0.8
1.0
b
BGCs as in a –>
BGCs as in a –>
log10(bitscore)
5.0
4.5
4.0

Cluster 6

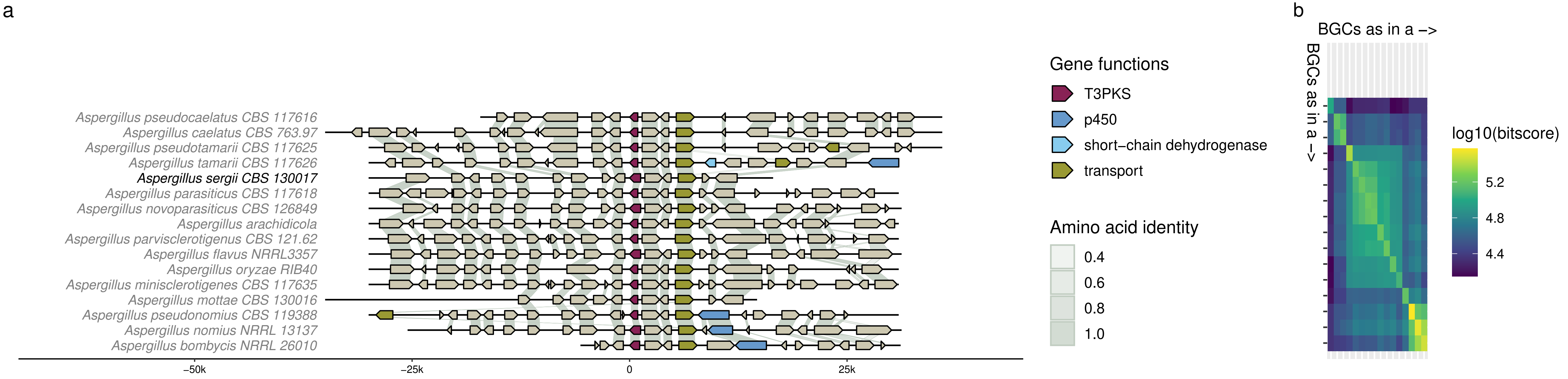
a
Aspergillus pseudocaelatus CBS 117616
Aspergillus caelatus CBS 763.97
Aspergillus pseudotamarii CBS 117625
Aspergillus tamarii CBS 117626
Aspergillus sergii CBS 130017
Aspergillus parasiticus CBS 117618
Aspergillus novoparasiticus CBS 126849
Aspergillus arachidicola
Aspergillus parvisclerotigenus CBS 121.62
Aspergillus flavus NRRL3357
Aspergillus oryzae RIB40
Aspergillus minisclerotigenes CBS 117635
Aspergillus mottae CBS 130016
Aspergillus pseudonomius CBS 119388
Aspergillus nomius NRRL 13137
Aspergillus bombycis NRRL 26010
−50k
−25k
0
25k
Gene functions
T3PKS
p450
short−chain dehydrogenase
transport
Amino acid identity
0.4
0.6
0.8
1.0
b
BGCs as in a −>
BGCs as in a −>
log10(bitscore)
5.2
4.8
4.4

Cluster 7

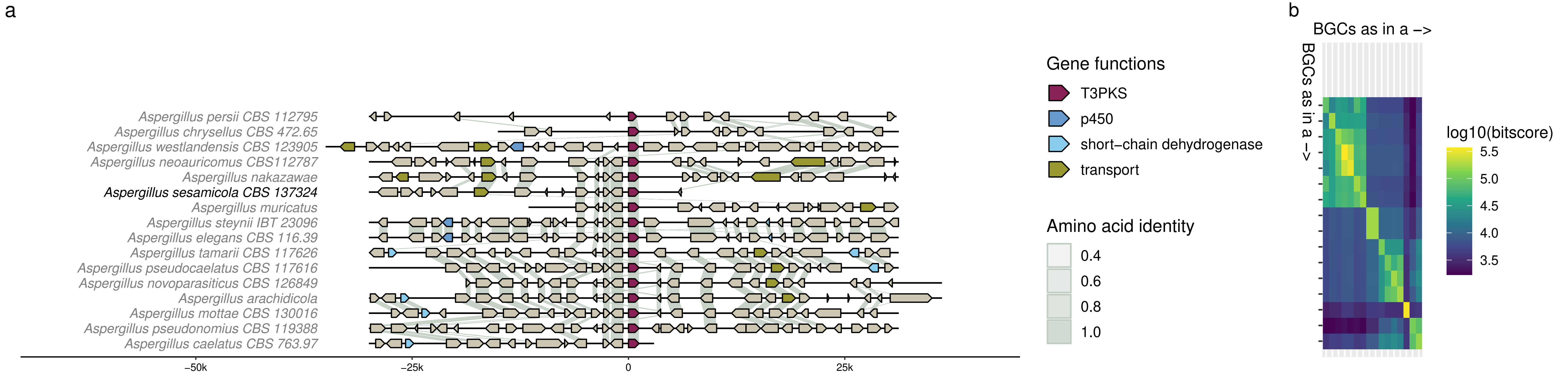

a
Aspergillus persii CBS 112795
Aspergillus chrysellus CBS 472.65
Aspergillus westlandensis CBS 123905
Aspergillus neoauricomus CBS112787
Aspergillus nakazawae
Aspergillus sesamicola CBS 137324
Aspergillus muricatus
Aspergillus steynii IBT 23096
Aspergillus elegans CBS 116.39
Aspergillus tamarii CBS 117626
Aspergillus pseudocaelatus CBS 117616
Aspergillus novoparasiticus CBS 126849
Aspergillus arachidicola
Aspergillus mottae CBS 130016
Aspergillus pseudonomius CBS 119388
Aspergillus caelatus CBS 763.97
−50k
−25k
0
25k
Gene functions
T3PKS
p450
short−chain dehydrogenase
transport
Amino acid identity
0.4
0.6
0.8
1.0
b
BGCs as in a −>
BGCs as in a −>
log10(bitscore)
5.5
5.0
4.5
4.0
3.5

Cluster 8

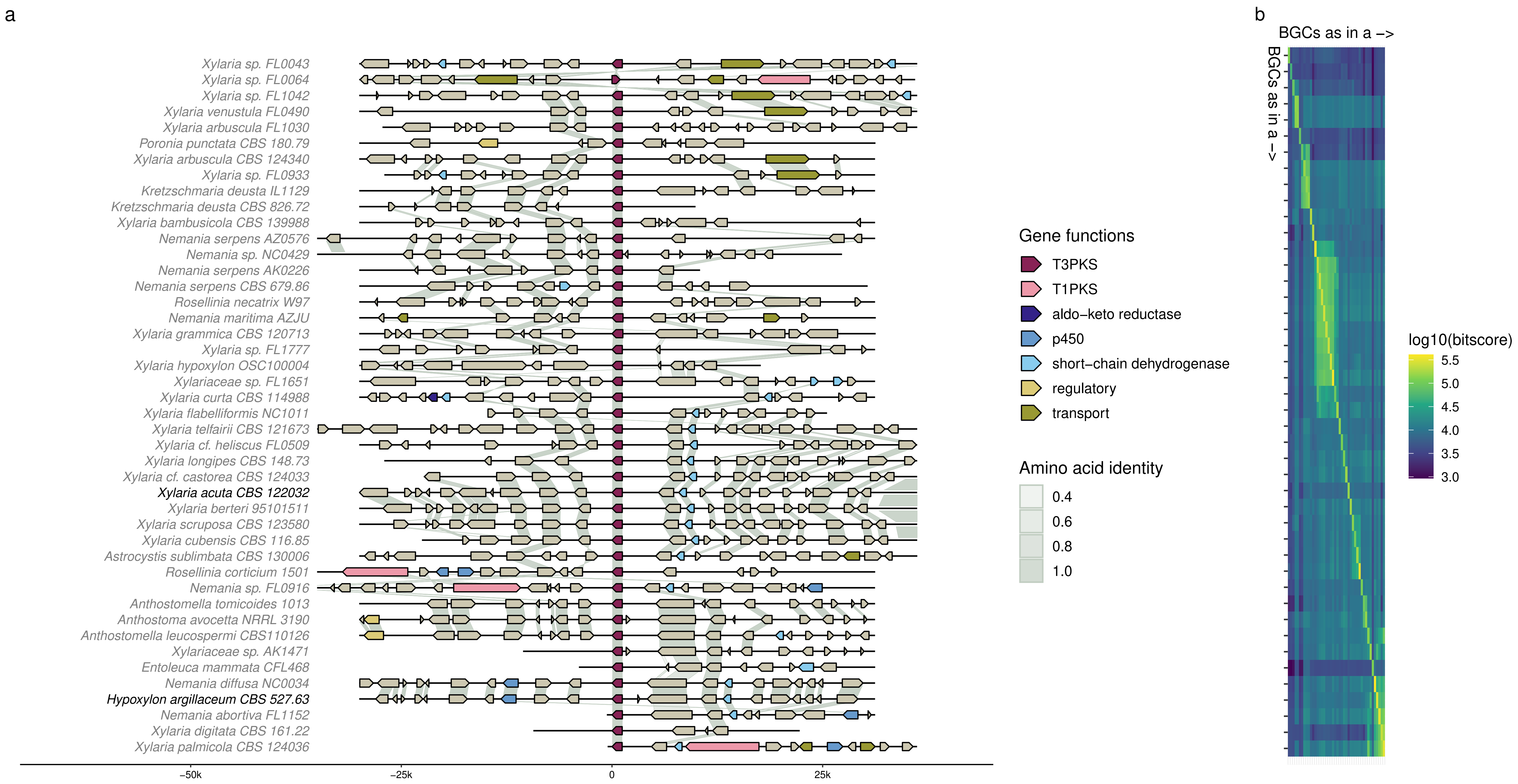
a
Xylaria sp. FL0043
Xylaria sp. FL0064
Xylaria sp. FL1042
Xylaria venustula FL0490
Xylaria arbuscula FL1030
Poronia punctata CBS 180.79
Xylaria arbuscula CBS 124340
Xylaria sp. FL0933
Kretzschmaria deusta IL1129
Kretzschmaria deusta CBS 826.72
Xylaria bambusicola CBS 139988
Nemania serpens AZ0576
Nemania sp. NC0429
Nemania serpens AK0226
Nemania serpens CBS 679.86
Rosellinia necatrix W97
Nemania maritima AZJU
Xylaria grammica CBS 120713
Xylaria sp. FL1777
Xylaria hypoxylon OSC100004
Xylariaceae sp. FL1651
Xylaria curta CBS 114988
Xylaria flabelliformis NC1011
Xylaria telfairii CBS 121673
Xylaria cf. heliscus FL0509
Xylaria longipes CBS 148.73
Xylaria cf. castorea CBS 124033
Xylaria acuta CBS 122032
Xylaria berteri 95101511
Xylaria scruposa CBS 123580
Xylaria cubensis CBS 116.85
Astrocystis sublimbata CBS 130006
Rosellinia corticium 1501
Nemania sp. FL0916
Anthostomella tomicoides 1013
Anthostoma avocetta NRRL 3190
Anthostomella leucospermi CBS110126
Xylariaceae sp. AK1471
Entoleuca mammata CFL468
Nemania diffusa NC0034
Hypoxylon argillaceum CBS 527.63
Nemania abortiva FL1152
Xylaria digitata CBS 161.22
Xylaria palmicola CBS 124036
−50k
−25k
0
25k
Gene functions
T3PKS
T1PKS
aldo−keto reductase
p450
short−chain dehydrogenase
regulatory
transport
Amino acid identity
0.4
0.6
0.8
1.0
b
BGCs as in a −>
BGCs as in a −>
log10(bitscore)
5.5
5.0
4.5
4.0
3.5
3.0

Cluster 9

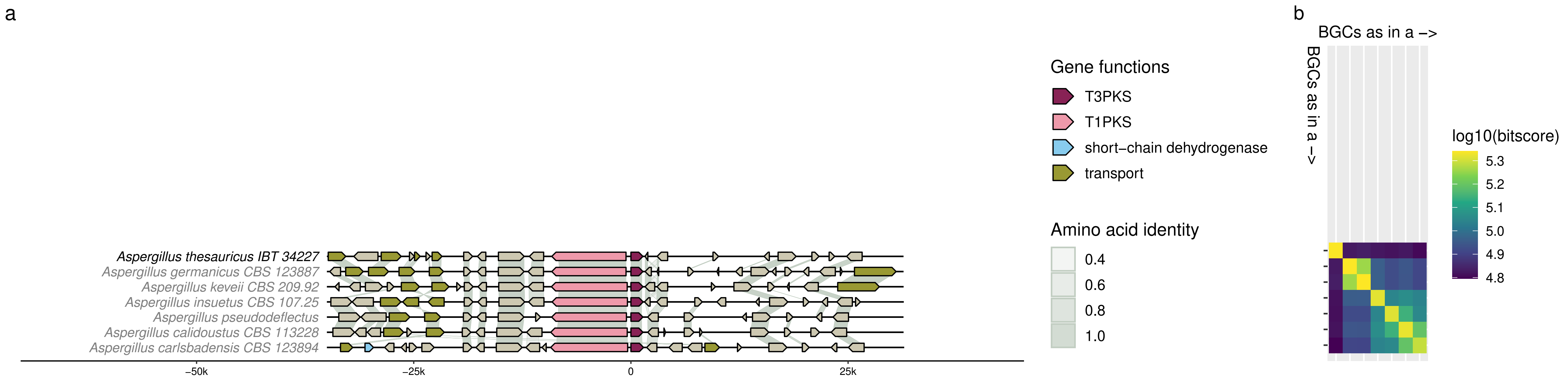
a
Aspergillus thesauricus IBT 34227
Aspergillus germanicus CBS 123887
Aspergillus keveii CBS 209.92
Aspergillus insuetus CBS 107.25
Aspergillus pseudodeflectus
Aspergillus calidoustus CBS 113228
Aspergillus carlsbadensis CBS 123894
−50k
−25k
0
25k
Gene functions
T3PKS
T1PKS
short−chain dehydrogenase
transport
Amino acid identity
0.4
0.6
0.8
1.0
b
BGCs as in a −>
BGCs as in a −>
log10(bitscore)
5.3
5.2
5.1
5.0
4.9
4.8

### Cluster 10

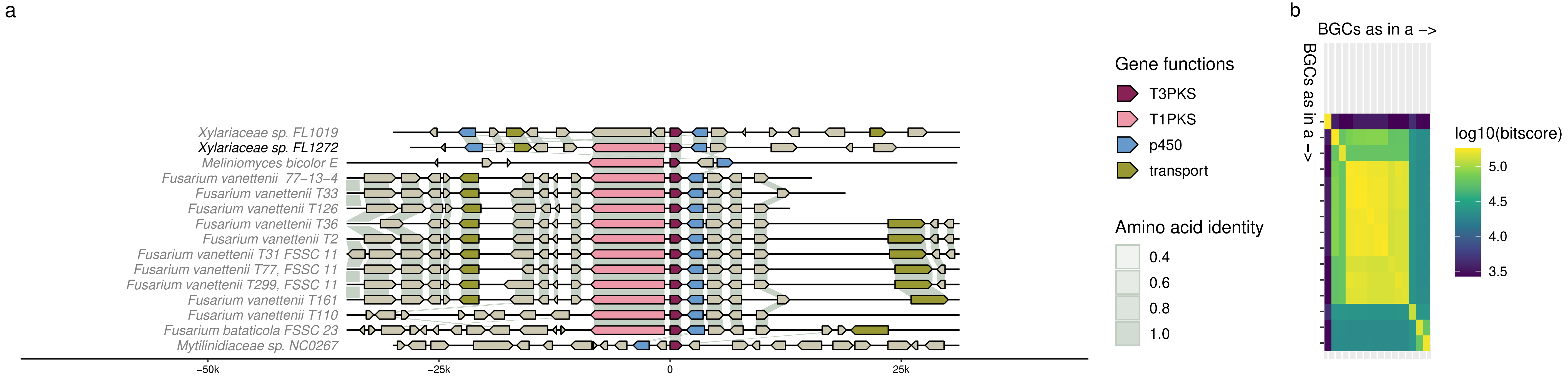
a
Xylariaceae sp. FL1019
Xylariaceae sp. FL1272
Meliniomyces bicolor E
Fusarium vanettenii 77−13−4
Fusarium vanettenii T33
Fusarium vanettenii T126
Fusarium vanettenii T36
Fusarium vanettenii T2
Fusarium vanettenii T31 FSSC 11
Fusarium vanettenii T77, FSSC 11
Fusarium vanettenii T299, FSSC 11
Fusarium vanettenii T161
Fusarium vanettenii T110
Fusarium bataticola FSSC 23
Mytilinidiaceae sp. NC0267
−50k
−25k
0
25k
Gene functions
T3PKS
T1PKS
p450
transport
Amino acid identity
0.4
0.6
0.8
1.0
b
BGCs as in a −>
BGCs as in a −>
log10(bitscore)
5.0
4.5
4.0
3.5

Cluster 11

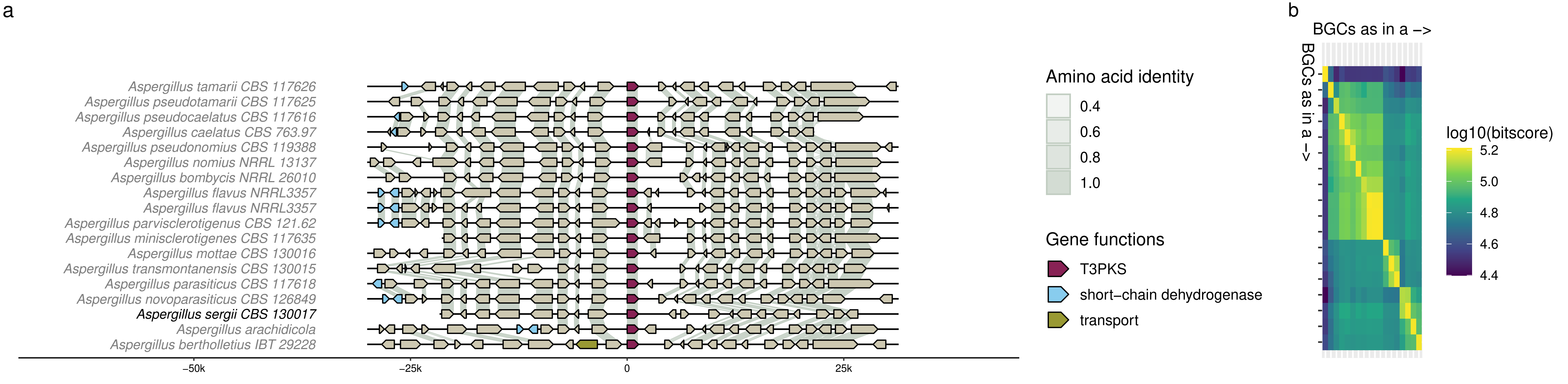

a
Aspergillus tamarii CBS 117626
Aspergillus pseudotamarii CBS 117625
Aspergillus pseudocaelatus CBS 117616
Aspergillus caelatus CBS 763.97
Aspergillus pseudonomius CBS 119388
Aspergillus nomius NRRL 13137
Aspergillus bombycis NRRL 26010
Aspergillus flavus NRRL3357
Aspergillus flavus NRRL3357
Aspergillus parvisclerotigenus CBS 121.62
Aspergillus minisclerotigenes CBS 117635
Aspergillus mottae CBS 130016
Aspergillus transmontanensis CBS 130015
Aspergillus parasiticus CBS 117618
Aspergillus novoparasiticus CBS 126849
Aspergillus sergii CBS 130017
Aspergillus arachidicola
Aspergillus bertholletius IBT 29228
−50k
−25k
0
25k
Amino acid identity
0.4
0.6
0.8
1.0
Gene functions
T3PKS
short−chain dehydrogenase
transport
b
BGCs as in a −>
BGCs as in a −>
log10(bitscore)
5.2
5.0
4.8
4.6
4.4

Cluster 12

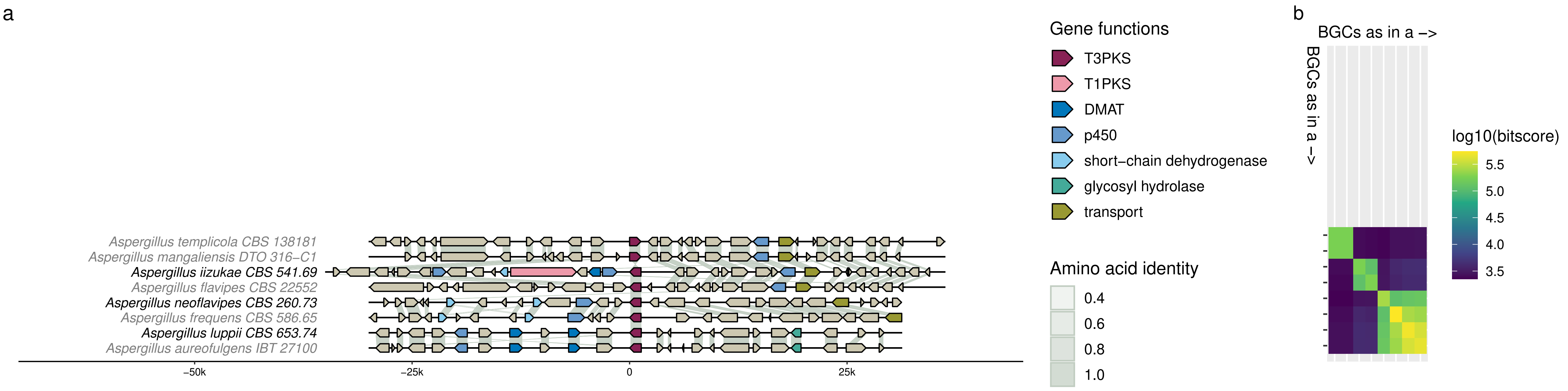
a
b
Aspergillus templicola CBS 138181
Aspergillus mangaliensis DTO 316−C1
Aspergillus iizukae CBS 541.69
Aspergillus flavipes CBS 22552
Aspergillus neoflavipes CBS 260.73
Aspergillus frequens CBS 586.65
Aspergillus luppii CBS 653.74
Aspergillus aureofulgens IBT 27100
−50k
−25k
0
25k
Gene functions
T3PKS
T1PKS
DMAT
p450
short−chain dehydrogenase
glycosyl hydrolase
transport
Amino acid identity
0.4
0.6
0.8
1.0
BGCs as in a −>
BGCs as in a −>
log10(bitscore)
5.5
5.0
4.5
4.0
3.5

Cluster 13

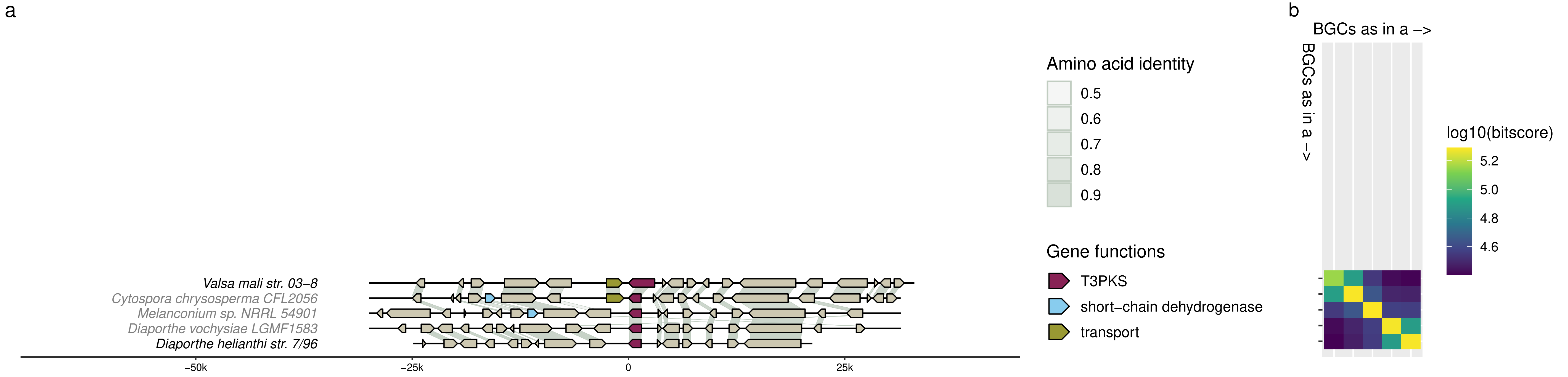

a
Valsa mali str. 03–8
Cytospora chrysosperma CFL2056
Melanconium sp. NRRL 54901
Diaporthe vochysiae LGMF1583
Diaporthe helianthi str. 7/96
–50k
–25k
0
25k
Amino acid identity
0.5
0.6
0.7
0.8
0.9
Gene functions
T3PKS
short–chain dehydrogenase
transport
b
BGCs as in a –>
BGCs as in a –>
log10(bitscore)
5.2
5.0
4.8
4.6

Cluster 14

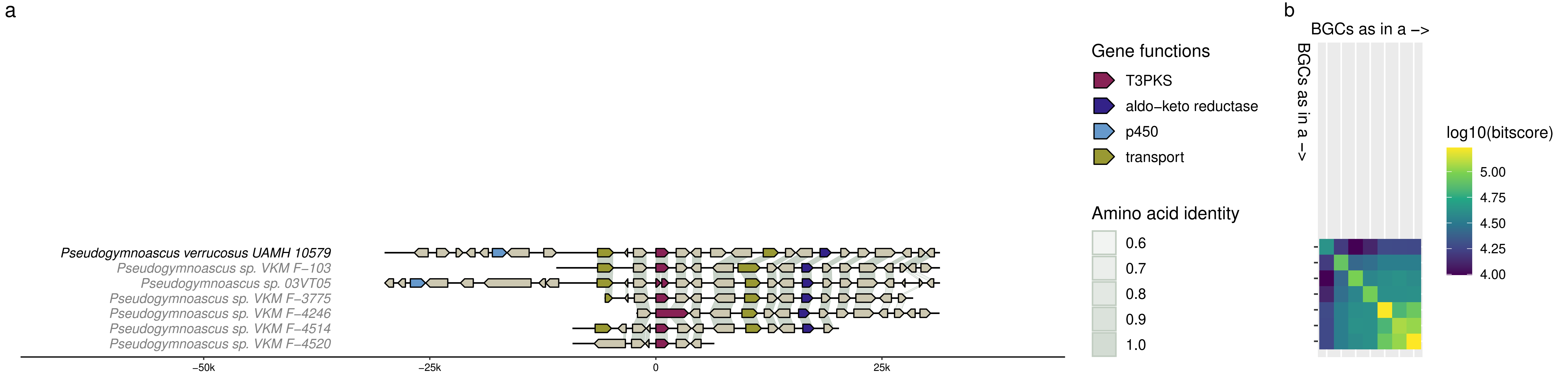
a
Pseudogymnoascus verrucosus UAMH 10579
Pseudogymnoascus sp. VKM F−103
Pseudogymnoascus sp. 03VT05
Pseudogymnoascus sp. VKM F−3775
Pseudogymnoascus sp. VKM F−4246
Pseudogymnoascus sp. VKM F−4514
Pseudogymnoascus sp. VKM F−4520
−50k
−25k
0
25k
Gene functions
T3PKS
aldo−keto reductase
p450
transport
Amino acid identity
0.6
0.7
0.8
0.9
1.0
b
BGCs as in a −>
BGCs as in a −>
log10(bitscore)
5.00
4.75
4.50
4.25
4.00

Cluster 15

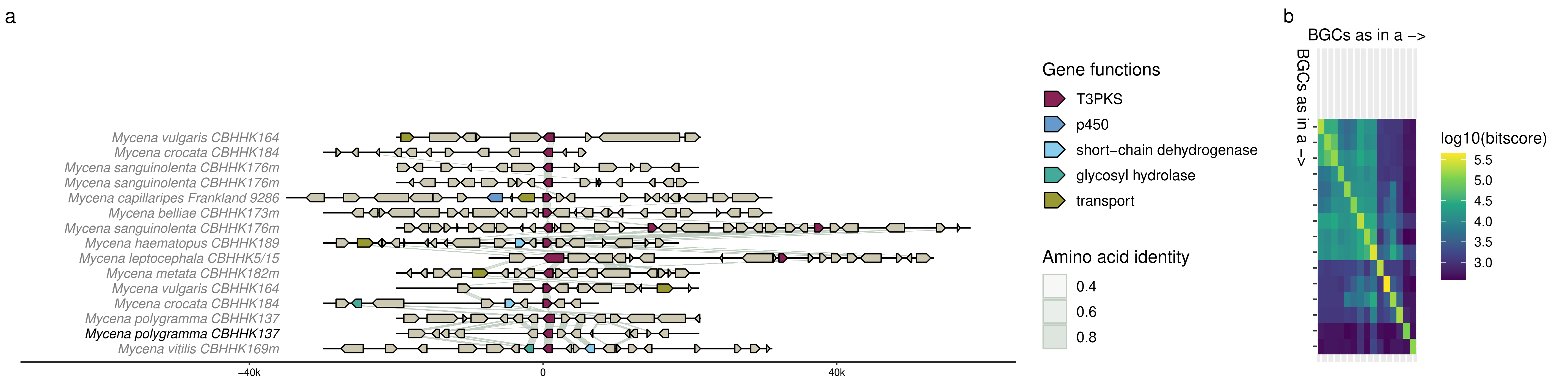

a
Mycena vulgaris CBHHK164
Mycena crocata CBHHK184
Mycena sanguinolenta CBHHK176m
Mycena sanguinolenta CBHHK176m
Mycena capillaripes Frankland 9286
Mycena belliae CBHHK173m
Mycena sanguinolenta CBHHK176m
Mycena haematopus CBHHK189
Mycena leptocephala CBHHK5/15
Mycena metata CBHHK182m
Mycena vulgaris CBHHK164
Mycena crocata CBHHK184
Mycena polygramma CBHHK137
Mycena polygramma CBHHK137
Mycena vitilis CBHHK169m
−40k
0
40k
Gene functions
T3PKS
p450
short−chain dehydrogenase
glycosyl hydrolase
transport
Amino acid identity
0.4
0.6
0.8
b
BGCs as in a −>
BGCs as in a −>
log10(bitscore)
5.5
5.0
4.5
4.0
3.5
3.0

Cluster 16

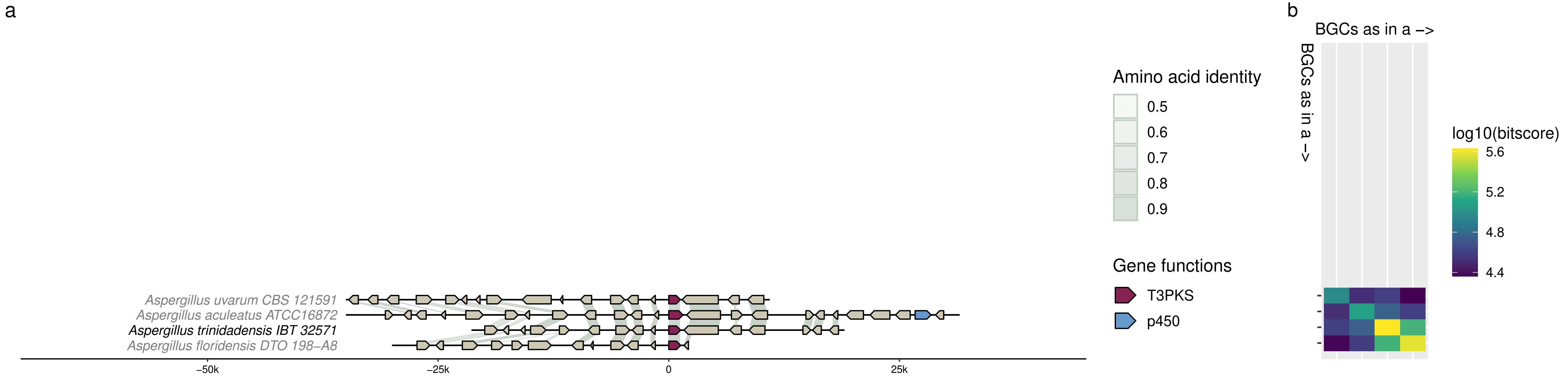

a
Aspergillus uvarum CBS 121591
Aspergillus aculeatus ATCC16872
Aspergillus trinidadensis IBT 32571
Aspergillus floridensis DTO 198–A8
−50k
−25k
0
25k
Amino acid identity
0.5
0.6
0.7
0.8
0.9
Gene functions
T3PKS
p450
b
BGCs as in a –>
BGCs as in a –>
log10(bitscore)
5.6
5.2
4.8
4.4

Cluster 17

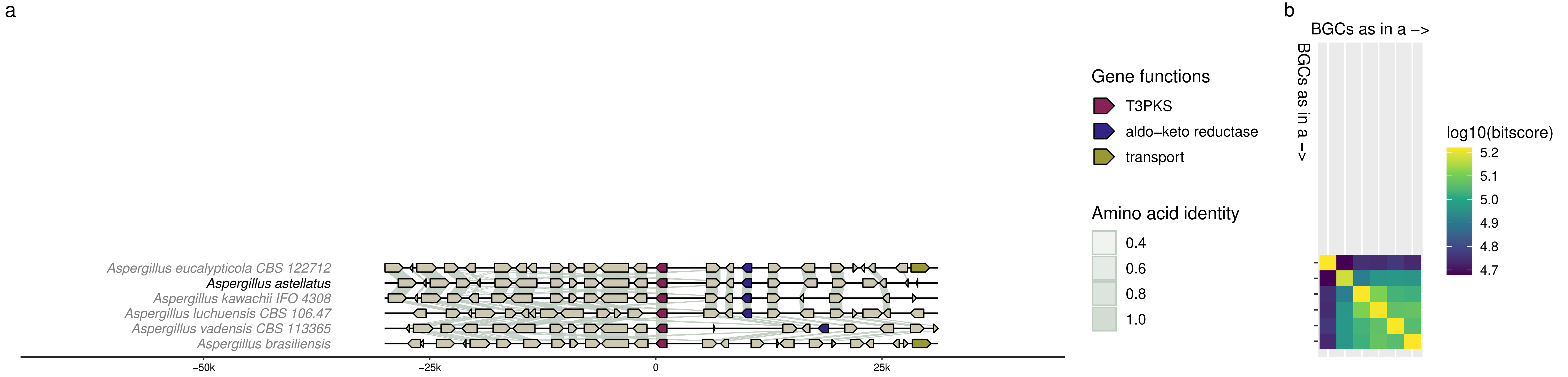
a
Aspergillus eucalypticola CBS 122712
Aspergillus astellatus
Aspergillus kawachii IFO 4308
Aspergillus luchuensis CBS 106.47
Aspergillus vadensis CBS 113365
Aspergillus brasiliensis
−50k
−25k
0
25k
Gene functions
T3PKS
aldo−keto reductase
transport
Amino acid identity
0.4
0.6
0.8
1.0
b
BGCs as in a −>
BGCs as in a −>
log10(bitscore)
5.2
5.1
5.0
4.9
4.8
4.7

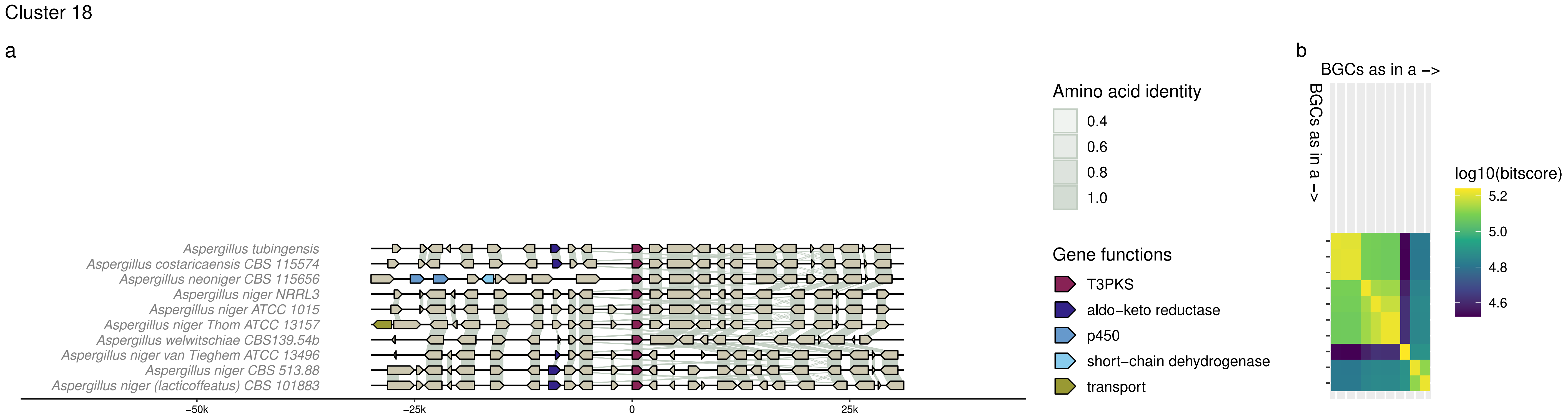

Cluster 18
a
Aspergillus tubingensis
Aspergillus costaricaensis CBS 115574
Aspergillus neoniger CBS 115656
Aspergillus niger NRRL3
Aspergillus niger ATCC 1015
Aspergillus niger Thom ATCC 13157
Aspergillus welwitschiae CBS139.54b
Aspergillus niger van Tieghem ATCC 13496
Aspergillus niger CBS 513.88
Aspergillus niger (lacticoffeatus) CBS 101883
−50k
−25k
0
25k
Amino acid identity
0.4
0.6
0.8
1.0
Gene functions
T3PKS
aldo−keto reductase
p450
short−chain dehydrogenase
transport
b
BGCs as in a −>
BGCs as in a −>
log10(bitscore)
5.2
5.0
4.8
4.6

Cluster 19

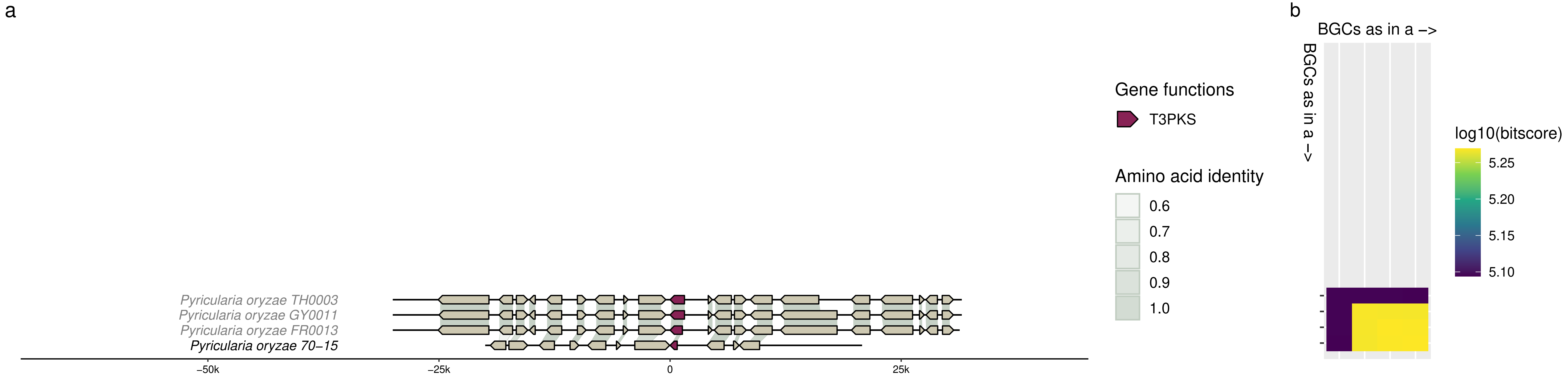

a
Pyricularia oryzae TH0003
Pyricularia oryzae GY0011
Pyricularia oryzae FR0013
Pyricularia oryzae 70−15
−50k
−25k
0
25k
Gene functions
T3PKS
Amino acid identity
0.6
0.7
0.8
0.9
1.0
b
BGCs as in a −>
BGCs as in a −>
log10(bitscore)
5.25
5.20
5.15
5.10

Cluster 20

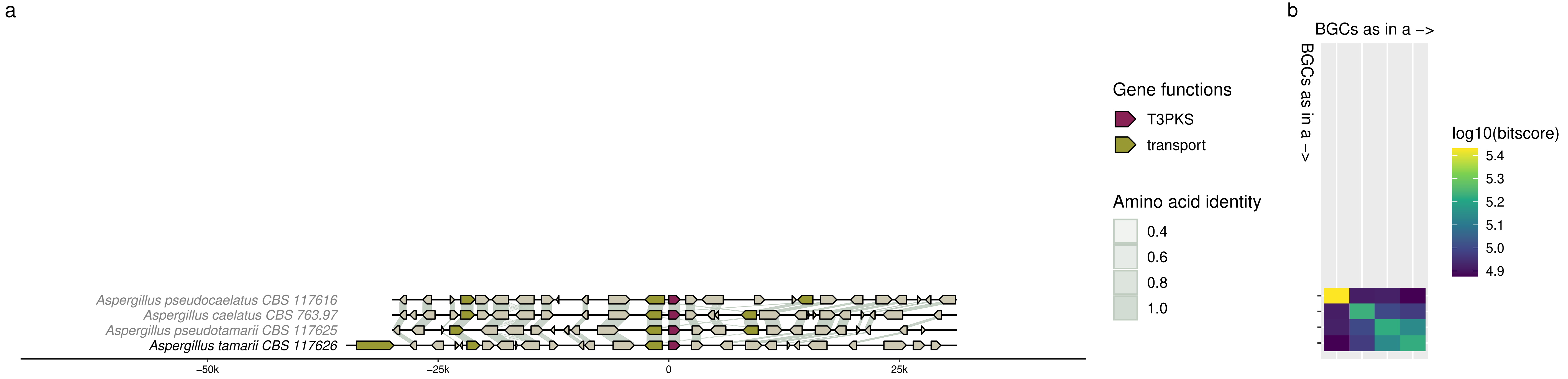

a
Aspergillus pseudocaelatus CBS 117616
Aspergillus caelatus CBS 763.97
Aspergillus pseudotamarii CBS 117625
Aspergillus tamarii CBS 117626
−50k
−25k
0
25k
Gene functions
T3PKS
transport
Amino acid identity
0.4
0.6
0.8
1.0
b
BGCs as in a −>
BGCs as in a −>
log10(bitscore)
5.4
5.3
5.2
5.1
5.0
4.9

Cluster 21

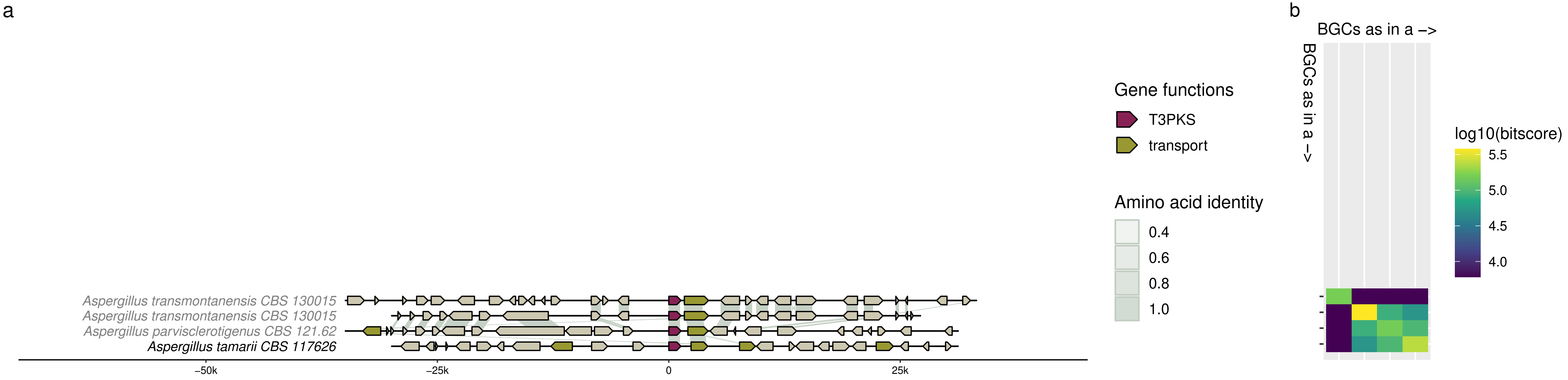
a
Aspergillus transmontanensis CBS 130015
Aspergillus transmontanensis CBS 130015
Aspergillus parvisclerotigenus CBS 121.62
Aspergillus tamarii CBS 117626
−50k
−25k
0
25k
Gene functions
T3PKS
transport
Amino acid identity
0.4
0.6
0.8
1.0
b
BGCs as in a −>
BGCs as in a −>
log10(bitscore)
5.5
5.0
4.5
4.0

Cluster 22

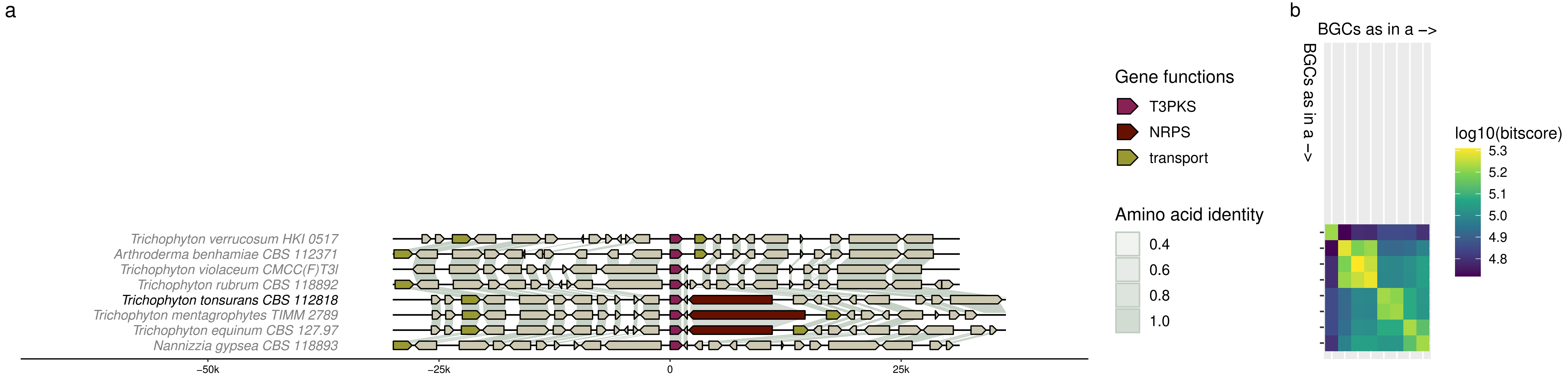

a
b
Trichophyton verrucosum HKI 0517
Arthroderma benhamiae CBS 112371
Trichophyton violaceum CMCC(F)T3l
Trichophyton rubrum CBS 118892
Trichophyton tonsurans CBS 112818
Trichophyton mentagrophytes TIMM 2789
Trichophyton equinum CBS 127.97
Nannizzia gypsea CBS 118893
−50k
−25k
0
25k
Gene functions
T3PKS
NRPS
transport
Amino acid identity
0.4
0.6
0.8
1.0
BGCs as in a –>
BGCs as in a –>
log10(bitscore)
5.3
5.2
5.1
5.0
4.9
4.8

Cluster 23

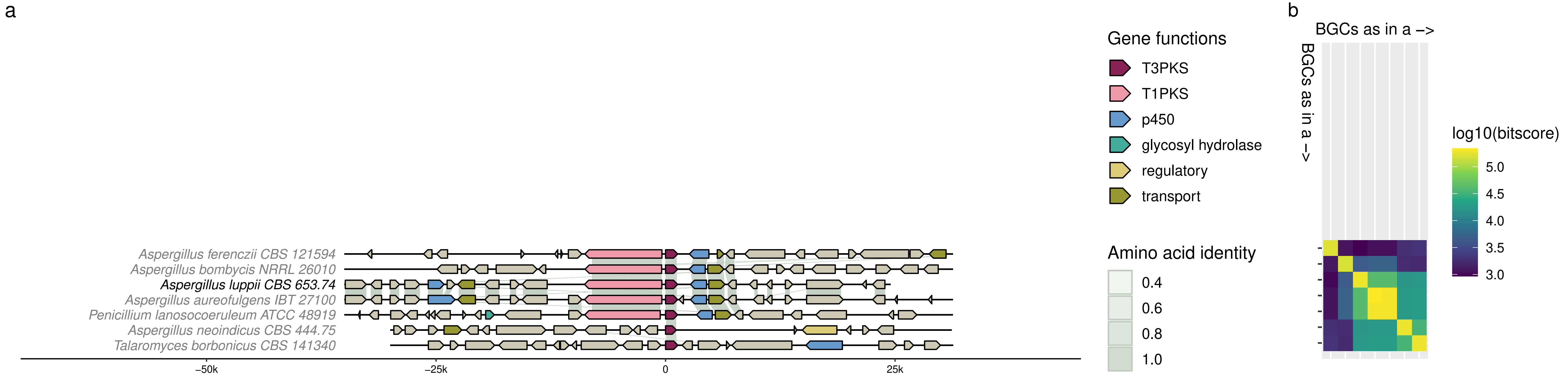
a
Aspergillus ferenczii CBS 121594
Aspergillus bombycis NRRL 26010
Aspergillus luppii CBS 653.74
Aspergillus aureofulgens IBT 27100
Penicillium lanosocoeruleum ATCC 48919
Aspergillus neoindicus CBS 444.75
Talaromyces borbonicus CBS 141340
−50k
−25k
0
25k
Gene functions
T3PKS
T1PKS
p450
glycosyl hydrolase
regulatory
transport
Amino acid identity
0.4
0.6
0.8
1.0
b
BGCs as in a –>
BGCs as in a –>
log10(bitscore)
5.0
4.5
4.0
3.5
3.0

Cluster 24

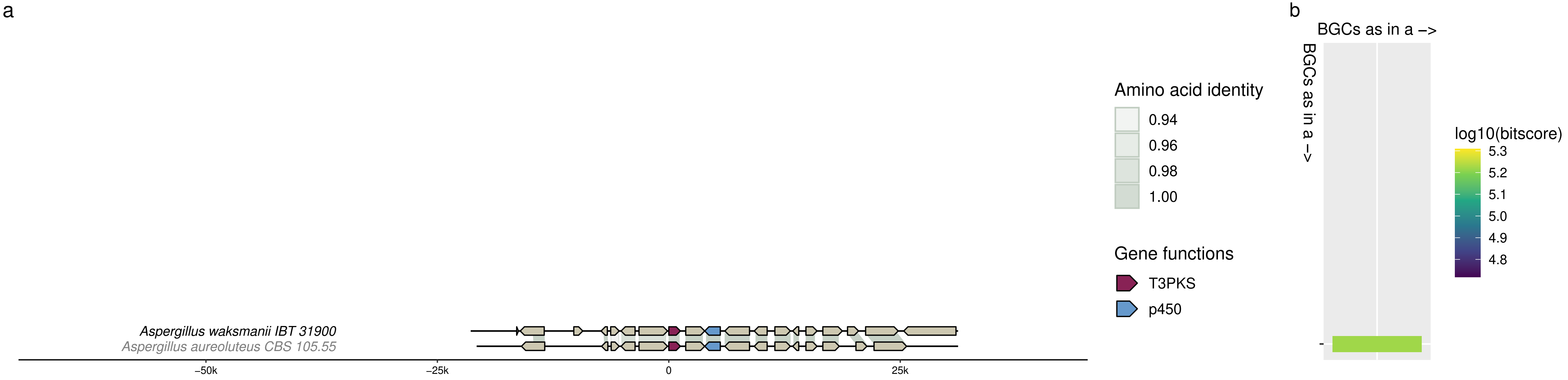
a
Aspergillus waksmanii IBT 31900
Aspergillus aureoluteus CBS 105.55
−50k
−25k
0
25k
Amino acid identity
0.94
0.96
0.98
1.00
Gene functions
T3PKS
p450
b
BGCs as in a −>
BGCs as in a −>
log10(bitscore)
5.3
5.2
5.1
5.0
4.9
4.8

### Cluster 25

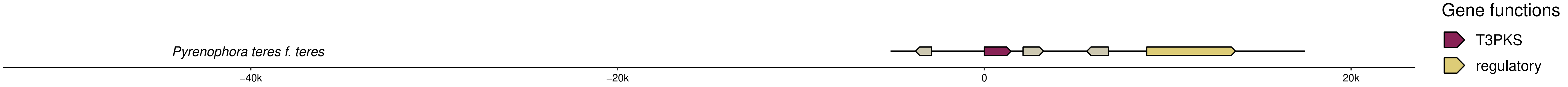
Pyrenophora teres f. teres
−40k
−20k
0
20k
Gene functions
T3PKS
regulatory
